# BRICseq bridges brain-wide interregional connectivity to neural activity and gene expression in single animals

**DOI:** 10.1101/422477

**Authors:** Longwen Huang, Justus M Kebschull, Daniel Furth, Simon Musall, Matthew T Kaufman, Anne K Churchland, Anthony M Zador

## Abstract

Comprehensive analysis of neuronal networks requires brain-wide measurement of connectivity, activity, and gene expression. Although high-throughput methods are available for mapping brain-wide activity and transcriptomes, comparable methods for mapping region-to-region connectivity remain slow and expensive because they require averaging across hundreds of brains. Here we describe BRICseq, which leverages DNA barcoding and sequencing to map connectivity from single individuals in a few weeks and at low cost. Applying BRICseq to the mouse neocortex, we find that region-to-region connectivity provides a simple bridge relating transcriptome to activity: The spatial expression patterns of a few genes predict region-to-region connectivity, and connectivity predicts activity correlations. We also exploited BRICseq to map the mutant BTBR mouse brain, which lacks a corpus callosum, and recapitulated its known connectopathies. BRICseq allows individual laboratories to compare how age, sex, environment, genetics and species affect neuronal wiring, and to integrate these with functional activity and gene expression.

## Introduction

A central problem in neuroscience is to understand how activity arises from neural circuits, how these circuits arise from genes, and how they drive animal behaviors. A powerful approach to solving this problem is to integrate information from multiple experimental modalities. Over the last decade, high-throughput approaches have enabled both gene expression (Rodriques et al., 2019; Ståhl et al., 2016; Vickovic et al., 2019) and functional neural activity (Macé et al., 2018; MacÉ et al., 2011; Musall et al., 2018; Prevedel et al., 2014; Sofroniew et al., 2016; Stirman et al., 2016; Vanni and Murphy, 2014) to be assessed at whole-brain scale in individual subjects. Unfortunately, it remains challenging to assess connectivity as rapidly and precisely. So the answers to fundamental questions of how connectivity is related to gene expression and neural activity, and how this relationship varies—in different species, genotypes, sexes and across developmental stages, as well as in animal models of neuropsychiatric disorders— remain elusive.

Historically, connectivity maps were compiled manually from results generated by many individual laboratories, each using somewhat different approaches and methods, and each presenting data relating to one or a few brain areas of interest in idiosyncratic formats (Bota et al., 2015; Felleman and Van Essen, 1991; Scannell et al., 1995). Recent studies avoid the confounds inherent in inferring connectivity across techniques and laboratories by relying on a standardized set of tracing techniques (Bohland et al., 2009; Harris et al., 2018; Markov et al., 2014; Oh et al., 2014; Zingg et al., 2014). Even with improved methods, however, such maps remain expensive and labor-intensive to generate, so region-to-region connectivity has been studied only for a small number of model organisms, typically of a single sex, age and genetic background (Markov et al., 2014; Oh et al., 2014; Zingg et al., 2014).

The major bottleneck in conventional tracing methods arises from the difficulty in multiplexing tracing experiments. In classical connectivity mapping, a single tracer—for example, a virus encoding green fluorescent protein (GFP)—is injected into a “source” brain area (Harris et al., 2018; Oh et al., 2014; Zingg et al., 2014). The brain is then dissected and imaged, and any region in which GFP-labeled axonal projections are observed is a projection “target”. Fluorescence intensity at the target is interpreted as the strength of the projection. This procedure must be performed in a separate specimen for each source region of interest, since multiple injections within a single specimen would lead to ambiguity about which injection was the source of the observed fluorescence (Figure 1A). Although multi-color tract tracing methods can achieve some multiplexing by increasing the number of fluorophores (Abdeladim et al., 2019; Zingg et al., 2014), the increase in throughput is modest because only a small number of colors can be reliably distinguished. To obtain a region-to-region connectivity map, data must be pooled across hundreds of animals, and the associated labor and costs limit the ability to generate the region-to-region connectivity maps from distinct model systems.

**Figure 1.**
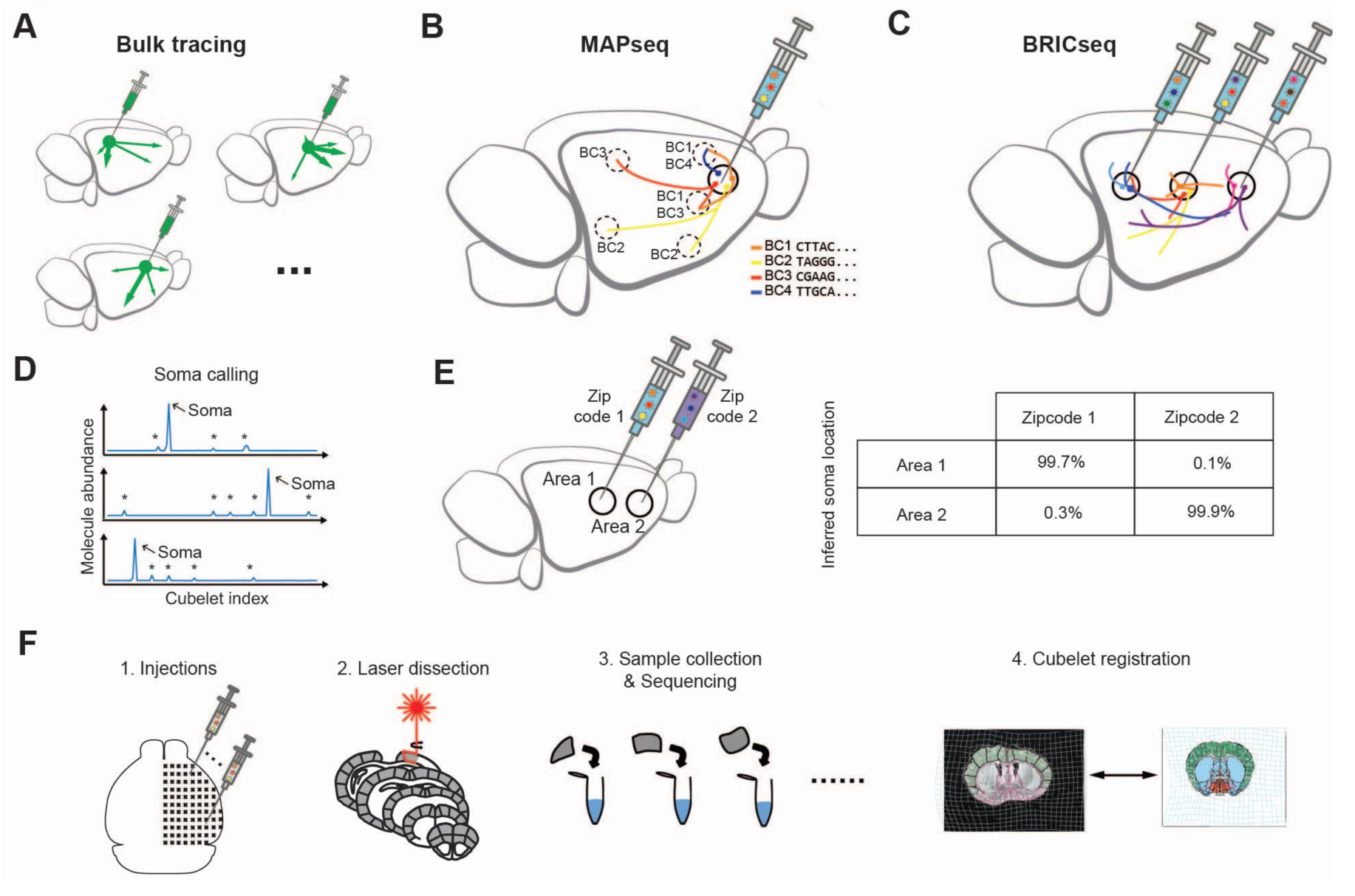
Mapping brain-wide cortico-cortical projections with BRICseq. **A**. In conventional fluorophore-based tracing, a separate brain is needed for each source area. **B**. In MAPseq, barcoded Sindbis virus is injected into a single source, and RNA barcodes from target areas of interest are extracted and sequenced. MAPseq multiplexes single neuron projections from a single source area. (*BC = barcodes*). **C**. In BRICseq, barcoded Sindbis is injected into multiple source areas. BRICseq multiplexes projections from multiple source areas, each at single neuron resolution. **D**. In the soma-max strategy for soma calling, the cubelet with the highest abundance of a particular barcode is posited to be the cubelet that contains the source (i.e. soma) of that barcode. **E**. Experimental validation of the soma-max strategy reveals an error rate <0.5%. **F**. BRICseq pipeline.

To achieve higher throughput at lower cost for mapping region-to-region connectivity in single animals, we sought to develop a method to enable multiplexing tracers for multiple source areas. Here we present BRICseq (<u>BR</u>ain-wide <u>I</u>ndividual-animal <u>C</u>onnectome <u>seq</u>uencing), which leverages barcoding and high-throughput sequencing to multiplex tracing experiments from multiple source areas, and allows for mapping of brain-wide corticocortical connectivity from individual mice in a few weeks, and at low cost. Using the map of mouse neocortex connectivity derived from BRICseq, we find that region-to-region connectivity provides a simple bridge for understanding the relationship between gene expression and neuronal activity. Applying BRICseq to the mutant BTBR mouse strain, we recapitulated its known connectopathies. The ability of BRICseq to map brain-wide connectivity from single animals in individual laboratories will foster the comparative and integrative analysis of connectivity, neural activity, and gene expression across individuals, animal models of diseases, and novel model species.

## Results

In what follows, we first describe the development of BRICseq, which allows mapping brain-wide projections from multiple sources in single animals. Next, we show that BRICseq is highly accurate and reproducible. We then show that BRICseq accurately predicts neural activity obtained by functional brain-wide calcium imaging in behaving mice, and that brain-wide gene expression predicts region-to-region connectivity. Finally, we show that BRICseq applied to the mutant BTBR mouse strain (which lacks a corpus callosum) can recapitulate its known connectopathies.

### BRICseq allows for multiplexing connectivity tracing from multiple source areas

The multi-site mapping strategy we developed, BRICseq, builds on MAPseq (Kebschull et al., 2016a). In MAPseq (Figure 1B), multiplexed single neuron tracing from a single source was achieved by labeling individual neurons with easily distinguishable nucleotide sequences, or “barcodes,” which are expressed as mRNA and trafficked into axonal processes. Because the number of nucleotide sequences, and therefore distinct barcodes, is effectively infinite—a short (30 base) random oligonucleotide has a potential diversity 4^30^≈10^18^—MAPseq can be thought of as a kind of “infinite color Brainbow” (Livet et al., 2007). Brain regions representing potential projection targets are microdissected into “cubelets” and homogenized, and the barcodes within each cubelet are sequenced, permitting readout of single cell projection patterns. The contribution of potential artifacts, including those due to degenerate labeling, fibers of passage, or non-uniform barcode transport, have been extensively quantified in previous work, and shown to be minimal (Chen et al., 2018; Han et al., 2018; Kebschull et al., 2016a) (Supplemental Note 1). MAPseq has now been validated using several different methods, including single neuron reconstruction, in multiple brain circuits (Chen et al., 2018; Han et al., 2018; Kebschull et al., 2016a).

MAPseq was originally developed to study projections from a single source. Conceptually, a straightforward generalization of MAPseq to determine the projections from many source areas in the same experiment would be to tag neurons with an additional area-specific barcode sequence—a “zipcode”—which could be used to identify the source (somatic origin) of each projection. In this approach, the overall strength of the projection from area 1 to area 2 would be determined by averaging the number of single neuron projections between those areas. In practice, however, such an approach would still be very labor intensive, because it would require the production, standardization and injection of hundreds of uniquely zipcoded batches of virus.

We therefore pursued a more convenient strategy, which requires only a single batch of virus (Figure 1C). We hypothesized that we could reliably determine the source of each projection using only sequencing, by exploiting the higher abundance of RNA barcodes in the somata compared with the axon terminals. According to this ‘soma-max’ strategy, the cubelet with the highest abundance of a given barcode of interest is assumed to be the soma (Figure 1D). To validate this soma-max strategy, we injected two distinct viral libraries, each labeled with a known zipcode, into two separate but densely connected cortical areas (primary motor area and secondary motor area). We dissected both injection sites, and sequenced the barcodes present in each. Compared to the ground truth determined by the zipcode, the soma-max strategy correctly identified the soma location for 99.2±0.2% of all cells (Figure 1E). These results indicate that the soma-max strategy would allow accurate reconstruction of connectivity even when only a single viral library is injected.

### Mapping brain-wide corticocortical region-to-region connectome with BRICseq

We first applied BRICseq to determining the region-to-region connectivity of the cortex of the adult male C57BL/6 mouse, for which there exist reference data sets (Oh et al., 2014; Zingg et al., 2014). To do so, we tiled the entire right hemicortex of each mouse with barcoded virus by making over 100 penetrations (3-6 injections/penetration at different depths) in a grid pattern with 500 μm edge length (Supplemental Table 2). Forty-four hours after viral injection, we cryosectioned the brain into 300 µm coronal slices, and used laser dissection to generate cortical (arc length ∼ 1 mm) and subcortical cubelets (Figure 1F, Figure S1C,D). The locations of all cortical cubelets were registered to the Allen Reference Atlas (2011 version, Figure 1F) (Fürth et al., 2018; Sunkin et al., 2013). We then quantified the number of each barcode sequence in each cubelet (Figure S1D).

In two adult male C57BL/6J mice (BL6-1 and BL6-2) we mapped the connections from 106±5 (mean±S.D.) source cubelets to 261.5±0.7 target cubelets (230.5±0.7 cortical, 31±0 subcortical). All dissected cubelets were potential targets; source cubelets were defined as the subset of all cubelets containing barcoded somata. From each source cubelet we obtained the sequences of several hundred somata (625±683) located therein, as well of projections from several thousand (1.5×10^3^±1.3×10^3^) neurons with somata located elsewhere. We aggregated these single neuron data to calculate region-to-region axonal projection strengths (Figure 2A,B, Figure S4A). Thus the strength of the projection from source cubelet X to target cubelet Y was defined as the number of barcodes in target Y originating from somata in source region X divided by the number of somata in X. We also estimated a confidence bound on our estimate of the strength of each connection (Figure S3G; Supplemental Note 3). All self-self projection strengths were set to 0. In addition, all the neighbor-projection strengths were also set to 0 to minimize false positives due to dendritic innervation of neighboring cubelets. Although in principle BRICseq data could have been used to determine single neuron projection patterns, in practice sequencing depth and other experimental details precluded such an analysis for this dataset (Figure S2; Supplemental Note 2.1).

**Figure 2.**
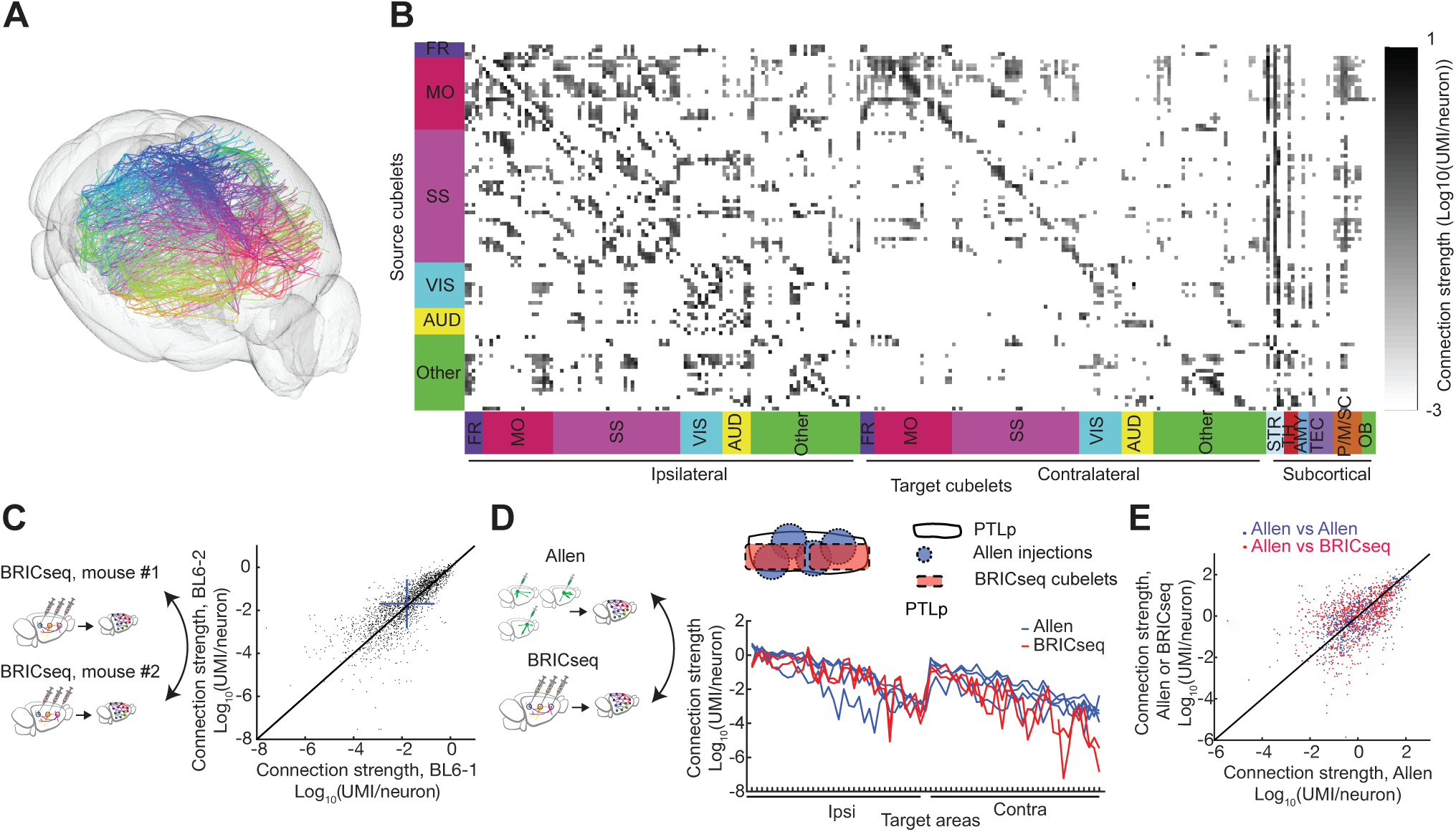
Brain-wide corticocortical projectome mapped by BRICseq and its validation. **A**,**B**. Cubelet-to-cubelet connectivity of mouse BL6-1. In **B**, Each row is a source cubelet, and each column is a target cubelet. Cubelets are assigned to their primary brain area. FR, frontal areas; MO, motor areas; SS, somatosensory areas; VIS, visual areas; AUD, auditory areas; STR, striatum; TH, thalamus; AMY, amygdala; TEC, tectum; P/M/SC, pons/medulla/spinal cord; OB, olfactory bulb. **C**. Reproducibility of brain area-to-brain area connection maps between two mice, BL6-1 and BL6-2. R = 0.7963, linear regression p < 10^−100^. The unity line is in black. Blue bars show mean±s.d. **D**,**E**. Connectivity determined by BRICseq agrees with the Allen Connectome Atlas. **D**, an example comparison of PTLp between the Allen Atlas and BRICseq of mouse BL6-1; **E**, Comparison of the whole network determined by BRICseq of BL6-2 with either the Allen Connectome or BRICseq of mouse BL6-2. **E**, Z-scores are plotted for each axis; Allen vs Allen, R = 0.5719, linear regression p < 10^−100^; Allen vs BRICseq, R = 0.5161, linear regression p < 10^−100^. The unity line is in black.

### BRICseq is reproducible and accurate

To fulfill its potential as a high-throughput method for determining connectivity, BRICseq must be both reproducible and accurate. To assess reproducibility, we compared across different BRICseq experiments. We first developed a computational pre-processing method to correct for variable experimental yields and/or sequencing depths across individual experiments (Figure S5, Supplemental Note 2.2). We next compared the two C57BL/6J connection maps, and found that the reproducibility of BRICseq was high. Estimated connection strengths were similar for the two brains tested (R = 0.83; Figure 2C; Supplemental Note 4). Differences between the measured connections in the two samples arose from some unknown combination of technical and biological variability. Major sources of technical variability likely include differences in injections and in dissection borders. We minimized biological variability by comparing subjects of the same age, sex and genetic background, but since the actual degree of animal-to-animal variability in cortical connections is unknown, these results represent an upper bound on the technical variability of BRICseq.

To assess the accuracy of BRICseq, we compared our results to the Allen Connectivity Atlas (Supplemental Table 2 in Oh et al., 2014), which was generated using conventional fluorophore-based techniques. The relationship between the ∼100 cortical BRICseq cubelets (defined by dissection) and cortical “areas” (defined by the Atlas) was not one-to-one: Each area typically spanned several cubelets, and each cubelet contributed to several areas. We therefore limited the comparison to the subset of cubelets that resided primarily (>70%) in a single source area (Supplemental Note 4). The agreement between BRICseq and the Allen Atlas was excellent (R=0.51); indeed, the agreement was comparable to inter-experiment variability within the Allen Atlas (R=0.57, Figure 2D,E). This confirms that potential MAPseq artifacts (from e.g. degenerate labeling, fibers of passage, non-uniform barcode trafficking) are minimal in BRICseq, as expected from previous work (Chen et al., 2018; Han et al., 2018; Kebschull et al., 2016a), and thus that BRICseq is a reliable method mapping region-to-region connectivity.

### Connectivity determined by BRICseq predicted neural activity

Every neuron in the cortex receives input from thousands of other neurons in other cortical and subcortical areas. Full knowledge of the detailed connections and activities of all the inputs would provide a foundation for the precise prediction of the activity of any given neuron (Bock et al., 2011; Kim et al., 2014; Seung and Sümbül, 2014; Takemura et al., 2013; Yan et al., 2017). However, BRICseq provides only region-to-region connectivity, a much lower dimensional measure. We therefore assessed whether BRICseq could predict neural activity.

We hypothesized that region-to-region anatomical connections would predict region-to-region “functional connectivity,” i.e. the statistical relationship between the neural activity in distinct brain regions (Friston, 2011). To measure functional connectivity, we performed cortex-wide wide-field calcium imaging in awake transgenic (Emx-Cre; Ai93; LSL-tTA) mice engaged in a visual/auditory decision task (Figure S6A) (Musall et al., 2018). In these mice, the calcium indicator GCaMP6f is expressed in excitatory cortical neurons. After registering calcium signals into the cubelet reference frame, the activity of each cubelet was calculated as the mean activity over all its pixels.

Figure 3 shows the relationship between anatomical connectivity measured by BRICseq and functional connectivity measured by wide-field calcium imaging. We used the correlation between pairs of cubelets as a measure of functional connectivity. Anatomical connectivity predicted functional connectivity remarkably well (R = 0.76). The excellent agreement between these two very different measurements suggests that much of the ongoing activity in the cortex can be explained by surprisingly simple interactions between connected areas.

**Figure 3.**
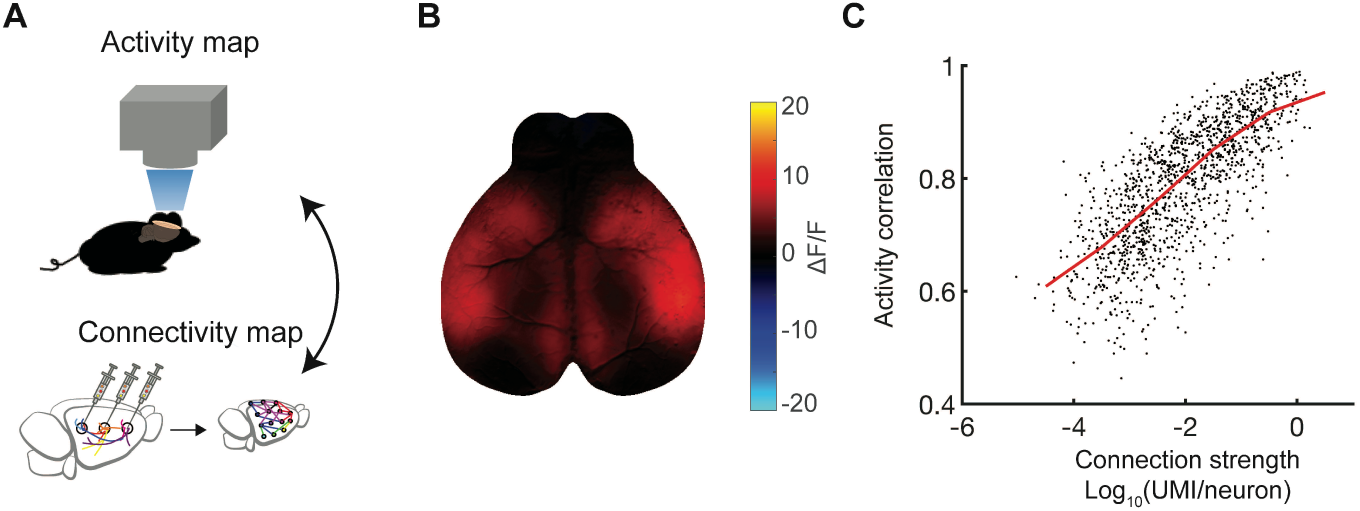
BRICseq predicts functional connectivity. **A**. BRICseq connectivity compared with cortex-wide Ca^2+^ imaging. **B**. A single frame example of cortex-wide wide-field calcium imaging in a behaving animal. **C**. Activity correlation between pairs of cubelets vs. reciprocal connection strengths between them. The median line is in *red*. n = 7225 pairs, R = 0.7637, linear regression p < 10^−100^.

### Connectivity determined by BRICseq could be predicted by low-dimensional gene expression data

We next set out to test whether gene expression could be used to predict connectivity (Fakhry and Ji, 2015; Fornito et al., 2019). We hypothesized that even though the patterns of gene expression that established wiring during development might have vanished at the time point we were examining, correlates of those patterns might persist into adulthood. We thus applied mathematical methods to search for gene expression patterns in the adult that could be used to predict the strengths of region-to-region connections (Figure 4A).

**Figure 4.**
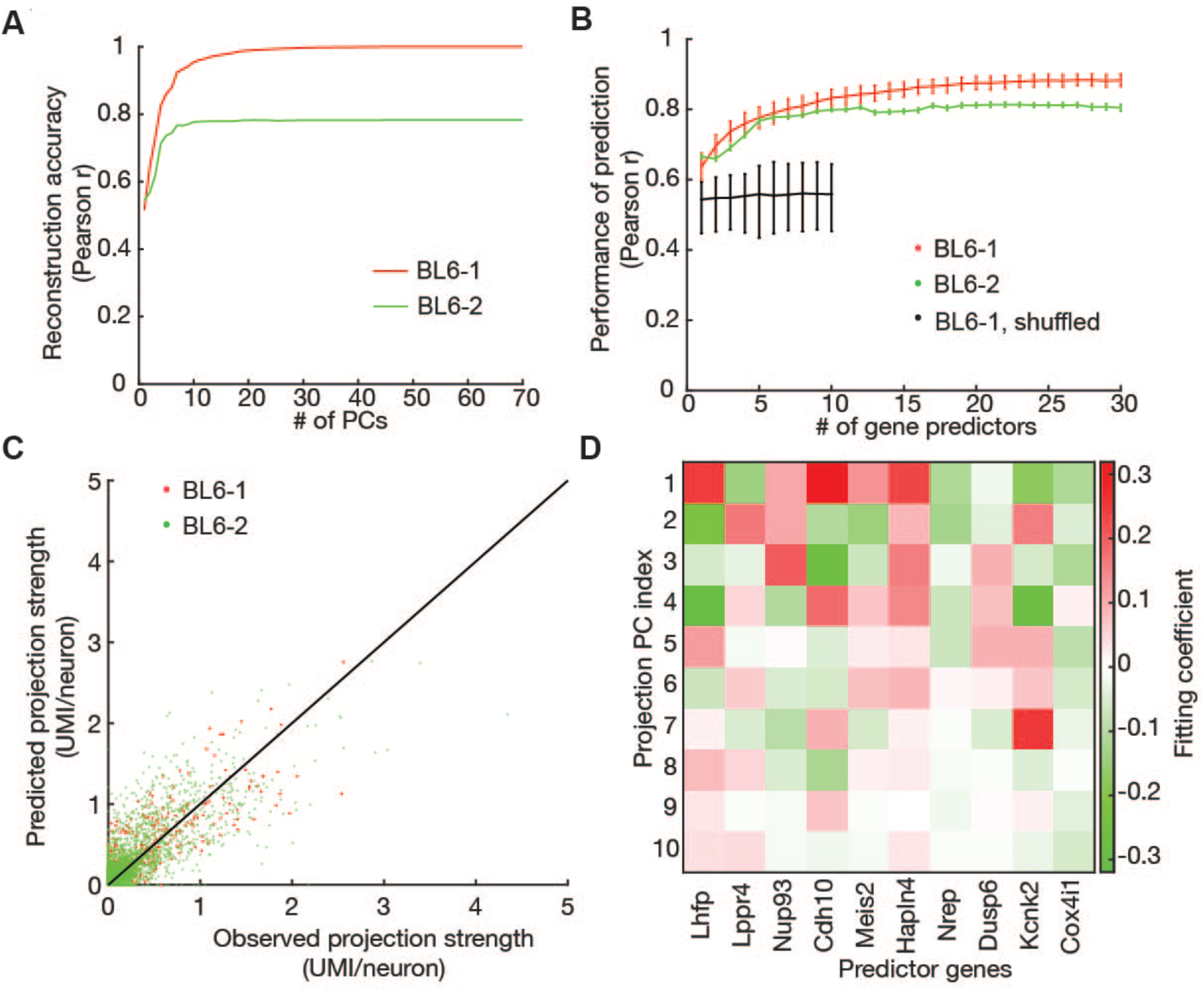
Gene expression patterns predict connectivity determined by BRICseq. **A**. PCA-based reconstruction of connectivity, using PCs and coefficients obtained from mouse BL6-1. The correlation coefficient is plotted between the connectivity reconstructed from first *n* PCs and either mouse BL6-1 (*red*) or BL6-2 (*green*). **B**,**C**. The performance of linear regression models using selected gene predictors. The linear models were trained using a training set (80%) in BL6-1, and then tested using the remaining testing set (20%) in BL6-1 as well as in BL6-2. **B**. The Pearson correlation between observed and predicted connectivity increases with the number of predictor genes. *Red*, the performance in the testing set in BL6-1. *Green*, the performance in BL6-2. *Black*, the null performance with the gene expression data shuffled before feature selection and linear regression (Supplemental Note 5.10.4). Error bars in red and green represent S.D.; error bars in black represent 95% confidence intervals. **C**. The scatter plot of observed versus predicted connectivity, using 10 gene predictors. *Red*, the testing set in BL6-1. *Green*, BL6-2. **D**. The fitting coefficients of top 10 gene predictors for top 10 connectivity PCs.

We first calculated cubelet-to-brain area connectivity based on BRICseq data, and used principal components analysis (PCA) to identify connectivity motifs shared between the two brains. In this analysis, the interpretation of each PC is a subset of correlated projection targets. Interestingly, a small number of the principal components (PCs) captured most of the variance in the connectivity data (Figure 4A; Figure S7B,C). Indeed, the reconstruction of brain connectivity based on just the first 10 PCs of brain BL6-1 was strongly correlated with both brain BL6-1 (r=0.95) and brain BL6-2 (r = 0.78). PCA can be thought of as a way of “de-noising” the brain connectivity, in the same way that low-pass filtering is a way of de-noising a periodic signal (exploiting the fact that sinusoids are the eigenvectors of a periodic signal). The motifs described by these first 10 PCs represent the components of the connectivity common to the two brains, and thus the components that could potentially be explained by gene expression data from an independent data set.

We next sought to predict the region-to-region connectivity from the gene expression in each cubelet. We first registered Allen *in situ* hybridization data, which depict the expression patterns of ∼20,000 genes in brains of male, 8 week-old, C57BL/6J mice (Lein et al., 2007), into the coordinates of BRICseq cubelets. We pre-filtered genes to only include high-quality expression data, and then used a greedy feature selection algorithm to identify 30 genes most effective for predicting connectivity using a linear model (Supplemental Note 5.10). Interestingly, prediction accuracy plateaued after only about 10 gene predictors to a surprisingly high level (BL6-1 testing set, r = 0.86±0.03; BL6-2, r = 0.83±0.004; Figure 4B-D; Figure S7D,E). Because of the highly correlated nature of gene expression, the identities these predictive genes were not unique; other sets of predictive genes performed about as well, consistent with the idea that these genes represent signatures of the genetic programs that established wiring during development. The ability of even a small number of marker genes to predict wiring agreement suggests that region-to-region connectivity patterns arise from low-dimensional genetic programs.

### BRICseq recapitulated known connectopathies in the BTBR mouse brain

A key advantage of muMPAseq is that it allows for rapid and systematic comparison of brain connectivity between model systems. We applied BRICseq to compare the cortical connectome of C57BL/6J (Figure 2B) to that of two BTBR mice (BTBR-1 and BTBR-2), an inbred strain lacking the corpus callosum and displaying social deficits (Fenlon et al., 2015; McFarlane et al., 2008; Wahlsten et al., 2003) (Figure 5A; Figure S8A). Most strikingly and as expected, BRICseq revealed a nearly complete absence of commissural cortical connections (Figure 5B,C; Figure S9A). In the C57BL/6J, commissural connections constitute 34.3±1.2% of total connections, whereas in BTBR the percentage is 1.0±0.8% (Figure 5D; Supplemental Note 2.7; the few remaining nonzero commissural connections in BTBR were found exclusively in target cubelets close to the midline, and likely represented dissection error and contamination from the ipsilateral hemisphere). Thus, the known connectopathies of the BTBR strain are recapitulated using BRICseq.

**Figure 5.**
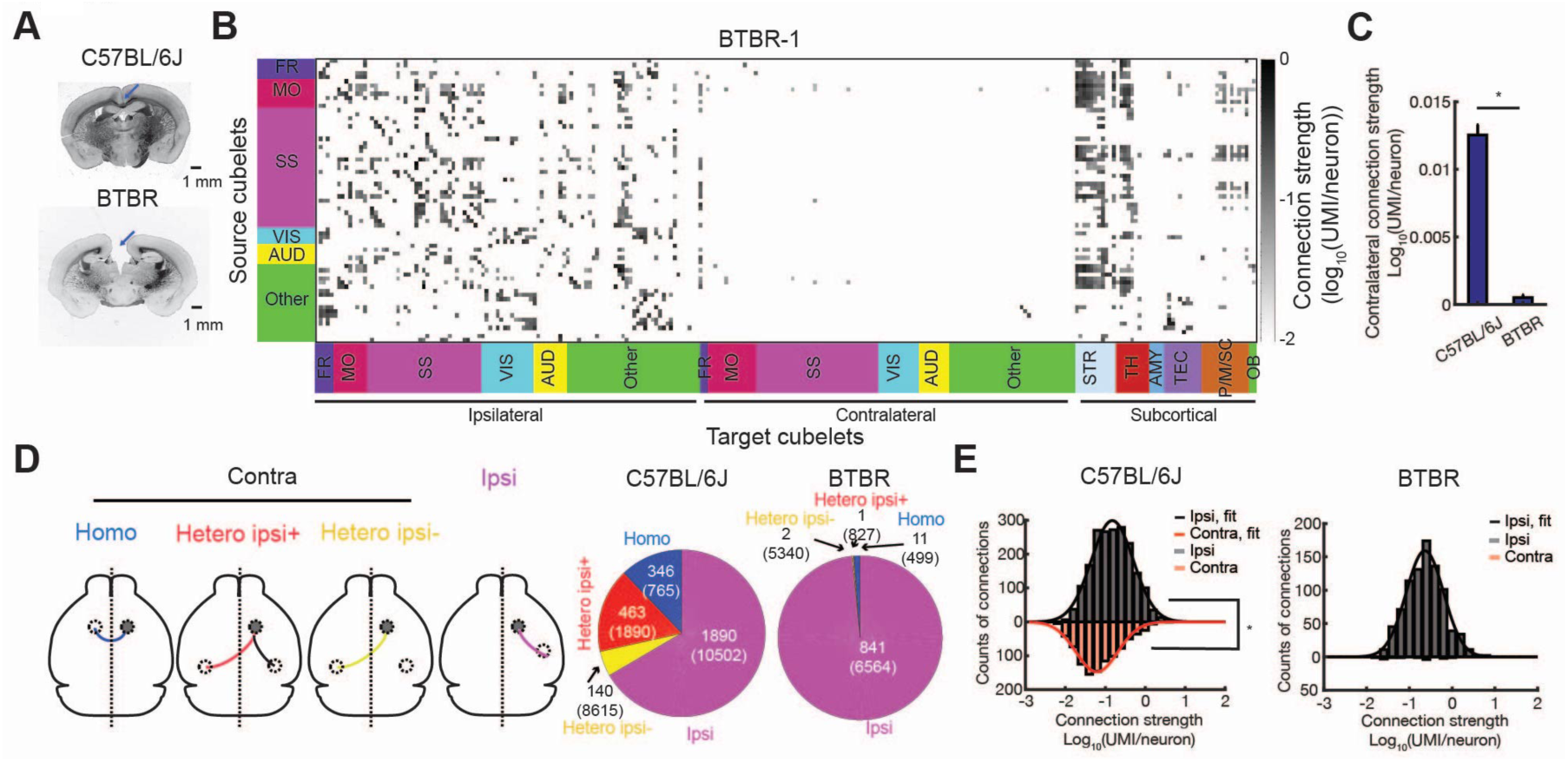
Comparison of the BTBR and C57BL/6J cortical connectivity. **A**. Bright field images of a C57BL/6J brain slice and a BTBR brain slice. Blue arrows indicate absence of the corpus callosum. **B**. Cubelet-to-cubelet connection matrix showing connection strengths in the BTBR mouse (BTBR-1). **C**. Quantification of contralateral connection strengths in C57BL/6J (BL6-1) and BTBR (BTBR-1). *, Mann-Whitney test, p < 10^−100^. Error bars represent S.E.M. **D**. Nonzero connections in C57BL/6J (BL6-1) and BTBR (BTBR-1). Numbers inside the parentheses indicate total counts of possible connections. Numbers outside the parentheses indicate total counts of non-zero connections. **E**. Distributions of ipsilateral/contralateral corticocortical connection strengths in C57BL/6J (BL6-1) and BTBR (BTBR-1). *, p < 10^−56^, Mann-Whitney test.

We next systematically compared the topological properties of the ipsilateral cortical networks of the C57BL/6J and BTBR mice in the cubelet coordinate system (Bullmore and Sporns, 2009). Network analyses of BRICseq-derived region-to-region connectivity differ from previous studies (Oh et al., 2014; Swanson et al., 2017; Zingg et al., 2014), as the natural coordinate frame is given by regularly spaced cubelets and all data were obtained from a single individual.

Consistent with previous reports (Oh et al., 2014), in the C57BL/6J, connection strengths were well fit by a log-normal distribution (Figure 5E, left). The decay of connection strength with distance (Figure S11A,D) was fit with a double exponential (λ_1_= 0.23±0.06 mm, λ_2_= 3.99±3.91 mm, 95% confidence intervals), and connection probability (Figure S11A,D) with a single exponential (λ= 0.92±0.16 mm, 95% confidence interval). Interestingly, the distribution of ipsilateral connection strengths in the BTBR was similarly fit by a log-normal distribution (Figure 5E, right), and the inferred ipsilateral region-to-region connections were not grossly disrupted (Figure S11H).

We next analyzed the topological properties of the ipsilateral cortical networks. By decomposing the network into small motifs containing 2 or 3 cubelets, and quantitatively comparing the abundance of these motifs to randomly generated networks, we found that in the C57BL/6J, the fraction of 2-cubelet motif with a reciprocally connected pair was greater than the null model, and densely connected 3-cubelet motifs were also significantly overrepresented (Figure 6A,B; Supplemental Note 2.5). Furthermore, four network modules—regions of the brain within which connections are dense, and which may reflect functional units—were revealed by connection-based clustering of cubelets in the C57BL/6J (Figure 6C,D Supplemental Note 2.6) (Zingg et al., 2014). Similar results were found in the BTBR (Figure S12E; Figure S13J-L), suggesting that these high-order topological properties were largely maintained in the BTBR strain. Thus, although the commissural corticocortical connections are completely missing, the ipsilateral network remained largely intact in the BTBR mouse.

**Figure 6.**
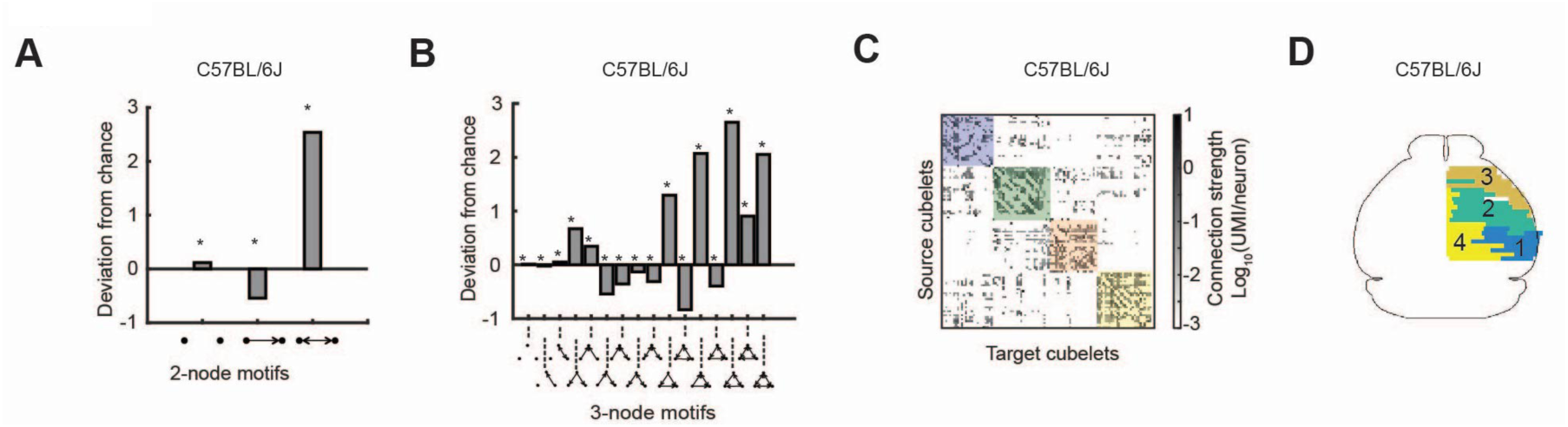
Topological properties of the ipsilateral cortical network. **A**,**B**. Abundance of 2-node and 3-node motifs in cortical network in C57BL/6J (BL6-1) compared to randomly generated networks. *, p < 0.001. **C**, Sorted cubelet-to-cubelet connection matrix based on modules in BL6-1. **D**. Connection-based modules in C57BL/6J (BL6-1). The same colors denote the same modules in **C** and **D**.

## Discussion

This study describes BRICseq, a high-throughput and low-cost method which exploits sequencing of nucleic acid barcodes for determining region-to-region connectivity in individual animals. BRICseq of the neocortex of the C57BL/6J mouse revealed that region-to region gene expression, connectivity and activity are related in a simple fashion: Spatial variations in as few as ten genes predict connectivity, and this connectivity in turn predicts correlations in neuronal activity. BRICseq of the BTBR mouse strain recapitulated the known deficits of commissural corticocortical connections. By virtue of its relatively low cost and high-throughput, BRICseq enables individual laboratories to study how age, sex, environment, genetics and species affect neuronal wiring, how these are disrupted in animal models of disease, and to integrate these with functional activity and gene expression.

### Comparison with other methods

Several approaches are available to map connectivity. Conceptually, BRICseq is closest to conventional fluorophore-based tracing techniques (Oh et al., 2014; Zingg et al., 2014). However, whereas conventional fluorophore-based approaches require pooling across hundreds of brains to map brain-wide connectivity, BRICseq multiplexes injections and is thereby able to map connectivity from individual subjects. This multiplexing reduces costs, labor, and animal-to-animal variability. The ability to generate maps from single subjects eliminates the need to register anatomical coordinate systems across animals, which increases reproducibility and accuracy. Reducing the number of subjects also leads to a substantial decrease in the total cost, both in terms of money and labor. The reduction in the number of subjects is particularly appealing for the study of non-human primates (Izpisua Belmonte et al., 2015), as well as of relatively new model systems for which connectivity maps are not yet available or individual subjects are particularly valuable, such as the Alston’s singing mouse (Banerjee et al., 2019; Okobi et al., 2019) and peromyscus (Bedford and Hoekstra, 2015; Metz et al., 2017; Weber et al., 2013).

Connectivity can also be mapped using diffusion tractography imaging (DTI), which uses 3D tracing of water diffusion pathways measured by MRI to infer the orientation of white matter tracts in the brain (Calabrese et al., 2015). Because DTI is rapid and non-invasive, it is widely used in the study of human brain connectivity. However, conventional DTI has low spatial resolution and low SNR, and has difficulty resolving subvoxel fiber complexity, so it has been much less useful in the study of small animal connectivity. Moreover, DTI requires access to specialized small animal MRI scanners, which remain relatively uncommon. Thus despite recent advances in small animal DTI, this approach has not become widely adopted.

BRICseq differs from conventional fluorophore tracing in that the spatial resolution is determined at the time of dissection, rather than as with fluorophore tracing at the time of injection. In the present study, we dissected rather large cubelets, but laser capture microdissection permits much smaller cubelets, even approaching single neuron resolution. Moreover, spatial transcriptomic methods (Rodriques et al., 2019; Ståhl et al., 2016; Vickovic et al., 2019), including *in situ* sequencing (Chen et al., 2018), raise the possibility of achieving single cell and indeed single axon or even synaptic resolution.

### Simple relationship among gene expression, connectivity and activity

At one level, our finding that there is a simple relationship (Figs. 3 & 4) among gene expression, connectivity and functional activity may not seem unexpected. The genome encodes the developmental rules for wiring up a brain—rules that are implemented in part by spatial patterns of gene expression—and this wiring in turn provides the scaffolding for resting state or “default” neuronal activity (Buckner et al., 2008). So the fact that gene expression, connectivity and functional activity are related is a direct consequence of development and brain architecture.

However, what is surprising is not that a relation exists among gene expression, connectivity and functional activity, but that this relationship is simple. Wiring could depend in complex and nearly indecipherable ways on dozens or even thousands of gene-gene interactions. Thus the fact that region-to-region connectivity of the neocortex could be predicted by the spatial expression pattern of just a small number (∼10) of genes raises the possibility that low-dimensional genetic programs determine the interregional wiring of the cortex. However, despite the predictive power of these 10 genes (Figure 4), there is no reason to expect that these predictive genes were causal in establishing wiring; they might merely be correlated with the causal genes. To establish the causal effect of genes on connectivity will likely require experiments in which gene expression is perturbed. Fortunately, BRICseq is sufficiently high-throughput that such an experimental program might not be prohibitively expensive.

We also observed that the corticocortical connectivity between two regions could predict correlations in cortical activity between them (Figure 3). Interestingly, a previous study (Honey et al., 2009) in humans found only a weak relationship between structural connectivity (assessed by DTI) and functional connectivity (inferred from resting state correlations). Whether these different results arise from methodological considerations (e.g. widefield calcium imaging and BRICseq vs. fMRI and DTI), or whether they reflect fundamental differences between mice and humans, remains to be determined.

In the present experiments, gene expression, connectivity and activity were all assessed separately, in different individuals. The data from these different experiments were then aligned to a shared coordinate system. However, because the techniques used in these experiments—widefield imaging, RNAseq of endogenous transcripts and sequencing of barcodes—are mutually compatible, it is feasible to combine them all in single individuals. Not only would this eliminate variability arising from combining data across individuals, it would also allow both connectivity and gene expression to be determined in the same coordinate system. Because the alignment to a common coordinate system represents a significant source of animal-to-animal variability, we expect that the simplicity of the relationships reported here represent a lower bound on the actual variability.

### BRICseq in the era of comparative connectomics

Growing evidence suggests that disruption of interregional connectivity leads to a variety of neuropsychiatric disorders, such as autism and schizophrenia (Geschwind and Levitt, 2007; Kubicki et al., 2007). Deciphering the circuit mechanisms underlying brain disorders requires systematic characterization of connectopathies, how they disrupt brain activity, and how they result from genetic mutations. Investigation of diverse animal models can reveal the neural mechanisms underlying species-specific behaviors, and provide a path toward discovering general brain principles (Yartsev, 2017). However, brain-wide interregional connectivity in animal models of diseases and new species remain largely unavailable, in part because of the lack of a high-throughput, inexpensive and accurate techniques. Thus, we expect that BRICseq, combined with other brain-wide individual-animal imaging or RNAseq techniques, will facilitate the creation of a systematic foundation for studying circuits in diverse animal models, opening up the possibility of a new era of quantitative comparative connectomics.

## Author Contributions and Notes

L.H. and J.M.K. performed the BRICseq experiments. S.M., M.T.K. and A.K.C. designed and performed the Ca^2+^ imaging experiments. D.F. designed whole brain visualizations. L.H., J.M.K and A.M.Z. designed the study, analyzed the data and wrote the paper. A.M.Z. is founder and equity owner in MapNeuro.

## Acknowledgments

We would like to thank Pavel Osten, Hongwei Dong and Liqun Luo for comments on the manuscript.

## STAR Methods

### Contact for reagent and resource sharing

Further information and requests for resources and reagents should be directed to and will be fulfilled by the Lead Contact, Anthony M Zador.

### Experimental model and subject details

Animal models used in the paper include: (model organism: name used in paper: genotype) Mouse: C57BL/6J: C57BL/6J; Mouse: BTBR: BTBR T^+^ Itpr3^tf^/J; Mouse: Emx-Cre: Emx1^tm1(cre)Krj^/J; Mouse: Ai93: Igs7^tm93.1(tetO-GCaMP6f)Hze^/J; Mouse: LSL-tTA: Gt(ROSA)26Sor^tm1(tTA)Roos^/J; Mouse: CamKII-tTA: CBA-Tg(Camk2a-tTA)1Mmay/J.

Animal procedures were approved by the Cold Spring Harbor Laboratory Animal Care and Use Committee and carried out in accordance with National Institutes of Health standards. For BRICseq, experimental subjects were 8-week-old male C57BL/6J mice or BTBR T^+^ Itpr3^tf^/J mice from the Jackson Laboratory. For functional imaging, triple transgenic mice Emx-Cre; Ai93; LSL-tTA were generated. A small fraction of mice used for functional imaging also harbored a CamKII-tTA allele to enhance the expression of GCaMP6f.

### Method details

#### Sindbis virus barcode libraries

The Sindbis virus used in BRICseq was made as described previously (Kebschull et al., 2016b, 2016a). Briefly, based on a dual promoter pSinEGdsp construct, we inserted MAPP-nλ after the first subgenomic promoter, and GFP-BC(barcode)-4×boxB after the second subgenomic promoter. Sequences (5’)AAG TAA ACG CGT AAT GAT ACG GCG ACC ACC GAG ATC TAC ACT CTT TCC CTA CAC GAC GCT CTT CCG ATC TNN NNN NNN NNN NNN NNN NNN NNN NNN NNN NNN GTA CTG CGG CCG CTA CCT A(3’) were inserted between MluI and NotI sites which were between GFP and 4×boxB. In barcode library 1, the 32-nt BC ended with 2 purines, while in barcode library 2, the 32-nt BC ended with 2 pyrimidines. Sindbis virus was produced using the DH-BB(5’SIN;TE12ORF) helper plasmid (Morris et al., 2011). One batch of library 1 viruses and two batches of library 2 viruses were used in the project. The viral barcode library diversity was determined by Illumina sequencing. ∼ 2 × 10^6^ barcodes were sequenced in the viral library 1, ∼ 8 × 10^6^ barcodes were sequenced in the first viral library 2 (used in BL6-1, BL6-2, BTBR-1, soma calling strategy validation experiment and template switching volume test experiment), and > 2.7 × 10^8^ barcodes were sequenced in the second viral library 2 (used in BTBR-2).

#### Injections

For BRICseq, Sindbis virus of barcode library 2 was injected into the right cortical hemispheres of experimental animals. Anesthesia was initially induced with isoflurane (4% mixed with oxygen, 0.5 L/min). Meloxican (2 mg/kg), dexamethasone (1 mg/kg) and baytril (10 mg/kg) were then administered subcutanesouly. For Sindbis injections, the whole skull above the right cortical hemisphere was removed. More than 100 injection pipette penetrations were made to cover the entire exposed brain, each spaced by 0.5 mm, both in the AP axis and ML axis. Nanoject III (Drummond Scientific) was used to inject Sindbis virus (∼2 × 10^10^ GC/mL), at 3-4 depths per penetration site (Supplemental Table 1). At each penetration site and depth, 23 nL virus was injected. The full injection surgery required about 8 hours, and constant isoflurane (1% mixed with oxygen, 0.5 L/min) was administered to maintain anesthesia. After injection, sterile Kwik-Cast (World Precision Instruments) was gently applied to cover the exposed brain region, and the skin was closed with sutures. Meloxican (2 mg/kg), dexamethasone (1 mg/kg) and baytril (10 mg/kg) were then routinely administered to animals subcutaneously every 12 hours post surgery, and animal condition was inspected every 6 – 12 hours. Similarly, we injected Sindbis virus of barcode library 1 into control animals. In control animals, instead of injecting the virus into the whole right cortex, we only made ∼6 penetrations covering a small cortical area.

For control experiments testing the soma calling strategy (Figure 1E), the same BRICseq protocol was followed, but Sindbis virus of barcode library 1 was injected into the secondary motor areas, and Sindbis virus of barcode library 2 into the primary motor areas.

For control experiments testing template switches (Figure S3B,C), we followed the BRICseq protocol above, but injected Sindbis virus of barcode library 2 into two separate animals.

For AAV CAG-tdTomato tracing experiments (Figure S8), we used coordinates AP = −4 mm, ML = 0.5 mm, 1 mm and 1.5 mm, DV = 0.25 mm and 0.5 mm for retrosplenial cortex in C57BL/6J and coordinates AP = −4 mm, ML = 0.75 mm, 1 mm and 1.5 mm, DV = 0.25 mm and 0.5 mm for retrosplenial in BTBR. In BTBR, as two hemispheres began to separate at AP = −4 mm and there was no cerebral cortex at ML = 0.5 mm, we used ML = 0.75 mm instead. In each coordinate, 20 nL of AAV1 CAG-tdTomato AAV (2×10^13^ GC/mL Penn Vector Core) was injected.

#### Cryosectioning and laser microdissection (LMD)

In BRICseq, 44 hours after Sindbis viral injection, the brain was harvested and fresh frozen at −80 °C. Olfactory bulbs and rostral spinal cord/caudal medulla were cut from the brain and collected separately. We then cut 300 µm coronal sections using a Leica CM 3050S cryostat at −12 °C chamber temperature and −10 °C object temperature. Each slice was cut with a fresh part of a blade, and the platform and brushes were carefully cleaned between slices. Each slice was immediately mounted onto a steel-framed PEN (polyethylene naphthalate)-membrane slide (Leica). After mounting on the slide, the slice was fixed in 75% ethanol at 4 °C for 3 min, washed in Milli-Q water (Millipore) briefly, stained in 0.5% toluidine blue (Sigma-Aldrich, MO) Milli-Q solution at room temperature for 30 sec, washed in Milli-Q water at room temperature for 3 times (15 sec each time), and fixed again in 75% ethanol at room temperature twice (2 min each time). The slide was then left in a vacuum desiccator for 30 min. Next, another fresh frame slide was used to sandwich the brain slice, and the two slides tightly taped to prevent the slice from falling. The sandwiched slice was stored in the vacuum desiccator at room temperature until LMD. If LMD was performed more than 1 week after cryosectioning, the sandwiched slices were stored at −80 °C in a desiccated container.

Cubelet dissection was performed with Leica LMD 7000. During LMD, cortical cubelets with ∼1 mm arc length were dissected from each coronal slice, from the surface to the deepest layer above the white matter. Orbitofrontal cortical cubelets (in rostral slices), anterior cingulate cortical cubelets, and retrosplenial cortical cubelets were also collected separately. For subcortical areas including striatum, thalamus, amygdala, tectum and pons/medulla, tissue belonging to each brain area was pooled every 1-3 consecutive slices. About 12∼21 cubelets were also collected from injection sites and contralateral homotopic areas of the injection sites in the barcode library 1 control animal, and 2 cortical cubelets in the uninjected control animal. Pictures were taken before and after every cubelet was dissected. After dissecting every 4 cubelets, we transferred them into homogenizing tubes with homogenizing beads, and added 100 µL lysis solution (RNAqueous-Micro Total RNA Isolation Kit, Thermo Fisher) into each cubelet. The collected tissues were stored temporally on dry ice and then at −80 °C.

#### Sequencing library preparation

After LMD, each cubelet was homogenized in lysis solution with a tissue lyser (Qiagen) at 20 Hz for 6 min. Then we extracted RNA molecules from each cubelet with RNAqueous-Micro Total RNA Isolation Kit (Thermo Fisher). We did not treat products with DNase I as DNA did not influence following experiments. The final product was eluted in 20 µL elution solution.

After RNA extraction, we performed reverse transcription (RT) with barcoded RT primers using SuperScript IV (Thermo Fisher). Barcoded RT primers were in the form of (5’)CTT GGC ACC CGA GAA TTC CAX XXX XXX XXX XXZ ZZZ ZZZ ZTG TAC AGC TAG CGG TGG TCG(3’), where Z_8_ is one of 288 CSIs (cubelet-specific identifiers) and X_12_ is the UMI (unique molecular identifier). 1 µL of 1 × 10^−9^ µg/µL spike-in RNAs were also added. The sequence of spike-in RNAs were (5’)GUC AUG AUC AUA AUA CGA CUC ACU AUA GGG GAC GAG CUG UAC AAG UAA ACG CGU AAU GAU ACG GCG ACC ACC GAG AUC UAC ACU CUU UCC CUA CAC GAC GCU CUU CCG AUC UNN NNN NNN NNN NNN NNN NNN NNN NAU CAG UCA UCG GAG CGG CCG CUA CCU AAU UGC CGU CGU GAG GUA CGA CCA CCG CUA GCU GUA CA(3’).

We then cleaned up RT products with 1.8×SPRI select beads (Beckman Coulter), synthesized double-stranded cDNA with previously described methods (Morris et al., 2011), cleaned up 2^nd^ strand synthesis products again with 1.8× SPRI select beads, and treated the eluted ds cDNA with Exonuclease I (New England Biolabs) (incubated the mix at 37°C for 1 hr and inactivated the enzyme at 80°C for 20 min). As cDNA molecules from different cubelets were already CSI-barcoded after RT, we pooled every 12 RT products for 1^st^ bead purification and 2^nd^ strand synthesis, and pooled all the products for 2^nd^ bead purification and Exonuclease I treatment.

We next amplified the cDNA library by nested PCR using primers (5’)GGA CGA GCT G(3’) and (5’) CAA GCA GAA GAC GGC ATA CGA GAT CGT GAT GTG ACT GGA GTT CCT TGG CAC CCG AGA ATT CCA(3’) for the first PCR and primers (5’)AAT GAT ACG GCG ACC ACC GA(3’) and (5’) CAA GCA GAA GAC GGC ATA CGA(3’) for the second PCR in Accuprime Pfx Supermix (Thermo Fisher). First PCR was performed for 5 cycles in 720 µL; after Exonuclease I treatment (incubated the mix at 37°C for 30 min and inactivated the enzyme at 80°C for 20 min), 1/4 of the first PCR products were used for second PCR. Second PCR was performed for 5-10 cycles in 12 mL. Standard Accuprime protocol was used for PCR except that the extension time in each cycle was set to 2 min to reduce incomplete elongation and template switches.

Nested PCR products were then purified and eluted in 600 µL with a Wizard SV Gel and PCR Clean-Up System (Promega), and further concentrated with Ampure XP beads (Beckman Coulter) in 25 µL Milli-Q H2O. After running in a 2% agarose gel, the 230 bp band was cut out and cleaned up with the Qiagen MinElute Gel Extraction Kit (Qiagen). We sequenced the library on an Illumina Nextseq500 high output run at paired end 36 using the SBS3T sequencing primer for paired end 1 and the Illumina small RNA sequencing primer 2 for paired end 2.

Most of the molecular experiments were performed according to the reagent manufacturer’s protocol unless otherwise stated.

#### Sequencing

We sequenced the pooled libraries prepared as above on an Illumina Nextseq500 high output run at paired end 36 using the SBS3T sequencing primer for paired end 1 and the Illumina small RNA sequencing primer 2 for paired end 2.

#### Confocal imaging

In AAV tracing experiments, brains were harvested 14 days after viral injection, fixed in 4% paraformaldehyde, washed in phosphate-buffered saline, and cut into 100 µm slices with a vibrotome (LeicaVT1000S, Leica). Slices were then mounted onto slides in Fluoroshield (Sigma-Aldrich), and imaged in a Laser Scanning Microscope 710 system (Leica).

#### Wide-field calcium imaging

Wide-field calcium imaging experiments in Figure 3 and Figure S6 are as described in Musall et al., 2018. All surgeries were performed under 1-2 % isoflurane in oxygen anesthesia. After induction of anesthesia, 1.2 mg/kg of meloxicam was injected subcutaneously and lidocaine ointment was topically applied to the skin. After making a medial incision, the skin was pushed to the side and fixed in position with tissue adhesive (Vetbond, 3M). We then created an outer wall using dental cement (Ortho-Jet, Lang Dental) while leaving as much of the skull exposed as possible, then a circular headbar was attached to the dental cement. After carefully cleaning the exposed skull we applied a layer of cyanoacrylate (Zap-A-Gap CA+, Pacer technology) to clear the bone. After the cyanoacrylate was cured, cortical blood vessels were clearly visible.

Widefield imaging was done using an inverted tandem-lens macroscope in combination with an sCMOS camera (Edge 5.5, PCO) running at 60 fps. The top lens had a focal length of 105 mm (DC-Nikkor, Nikon) and the bottom lens 85 mm (85M-S, Rokinon), resulting in a magnification of 1.24x. The total field of view was 12.4 × 10.5 mm and the spatial resolution was ∼20µm/pixel. To capture GCaMP fluorescence, a 500 nm long-pass filter was placed in front of the camera. Excitation light was coupled in using a 495 nm long-pass dichroic mirror, placed between the two macro lenses. The excitation light was generated by a collimated blue LED (470 nm, M470L3, Thorlabs) and a collimated violet LED (405 nm, M405L3, Thorlabs) that were coupled into the same excitation path using a dichroic mirror (#87-063, Edmund optics). From frame to frame, we alternated between the two LEDs, resulting in one set of frames with blue and the other with violet excitation at 30 fps each. Excitation of GCaMP at 405 nm results in non-calcium dependent fluorescence, and we could therefore isolate the true calcium-dependent signal by rescaling and subtracting frames with violet illumination from the preceding frames with blue illumination. All subsequent analysis was based on this differential signal at 30 fps.

#### Behavior task

For Figure 3 and Figure S6, the behavior has previously been described in Musall et al., 2018. Briefly, ten mice were trained on a delayed 2-alternative forced choice (2AFC), spatial discrimination task. Mice initiated trials by touching either of two handles. After one second of holding the handle, mice were presented with either a sequence of auditory clicks, or repeated presentation of a visual moving bar (3 repetitions, 200 ms each), for a total of 600 ms. Stimuli were positioned either left or right of the animal. After the 600 ms period, the animals experienced a 500 ms delay with no stimulus, followed by a second 600 ms period containing a repeat of the sensory stimuli. A 1000 ms delay was then imposed, after which servo motors moved two lick spouts into close proximity of the animal’s mouth. Licks to the spout corresponding to the stimulus presentation side were rewarded with water. After one spout was contacted, the opposite spout was moved out of reach to force the animal to commit to its initial decision. Animals were trained over the course of approximately 30 days, and reached stable detection performance levels of 80% or higher.

Each animal was trained exclusively on a single modality (6 vision, 4 auditory). During imaging sessions, both modalities were presented in different trials.

### Quantification and statistical analysis

#### Wide-field calcium imaging processing

To preprocess widefield data, we used SVD to compute the 500 highest dimensions accounting for more than 88 % of the variance in the data. The original data matrix M (of size pixels × frames) was decomposed as

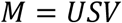

which returns ‘spatial components’ U (of size pixels × components), ‘temporal components’ V (of size components × frames) and singular values S (of size components × components) to scale components to match the original data. Further analysis methods are described in Supplemental Note 5.9.

#### LMD (laser microdissection) Image processing

Wholebrain toolbox (by Daniel Fürth, http://www.wholebrainsoftware.org) was used to register Toluidine Blue-stained coronal slices into Allen Reference Atlas semi-automatically. Using Matlab, we determined the coordinates of each cubelet by processing pictures taken before and after each cubelet was dissected. Combining image registration results and cubelet coordinates, we mapped each cubelet into one or multiple brain areas.

#### BRICseq data analysis

The details on BRICseq analysis, including bioinformatics, statistics and computational methods can be found in Supplemental Note 5.

#### BRICseq data visualization

BRICseq data were visualized in a 3D brain in Figure 2A. To reconstruct the cubelet-to-cubelet connection pathways, the position in stereotactic coordinates for each registered cubelet source node was used to query Allen Mouse Brain Connectivity Atlas (Oh et al., 2014) for injection sites within 500 µm from each source node. Out of all the injection sites the injection with largest injection volume was used to download projection density volumes with 200 µm voxel resolution. 92 out of 99 cubelet source nodes could be mapped to a unique projection density volume. Next, we used A* search algorithm (Sur and Taipale, 2016) implemented in C/C++ to find the optimal path between BRICseq source and target cubelet nodes using binary projection density volume to represent graph nodes and blocked obstacles. The optimal path for 1677 out of 3015 non-zero connection could be determined (56%). The remaining either didn’t have a corresponding projection density volume, alternatively target and source cubelets were not connected in the projection density volume. Each projection path was then smoothed as a spline using a Generalized Additive Model (GAM) (Chambers and Hastie, 2017). Each path was rendered in 3D with a unique color given by the position of the path’s target cubelet. The color-coding of target cubelet locations was based on a red-green-blue (RGB) spatial color cube code where red represents medio-lateral, green represents anterior-posterior, and blue represents dorso-ventral axis.

## Data and software availability

All sequencing datasets are publicly available under SRA accession codes SRA: PRJNA541990. Further information and requests for data should be directed to and will be fulfilled by the Lead Contact, Anthony M Zador.

