## Supplemental Notes for "BRICseq bridges brain-wide interregional connectivity to neural activity and gene expression in single animals"

### Supplemental Figures

**Figure S1, related to Figure 1. Additional BRICseq protocol. A**. Engineered Sindbis virus for BRICseq. **B**. A 300 µm coronal slice in a mouse brain 44 hours after Sindbis injection. Scale bar, 500 µm. **C**. A brain slice was stained with Toluidine Blue for laser micro-dissection. Red lines show contours of dissected cubelets. **D**. BRICseq pipeline.

**Figure S2, related to Figure 2. Single cell projection patterns. A**. Single cell projection patterns from over 600 cells residing in an example cubelet (cubelet 53) in BL6-1. Each row is one cell, and each column is a target cubelet. Cells are clustered by non-negative matrix factorization (Supplemental Note 2.1). **B**. Dorsal view of projections of 3 example cells in cubelet 53. The circles indicate the location of the source cubelet. **C**. Distribution of the number of single cell projection targets in BL6-1. **D**. A cartoon illustrating that low sequencing depth may influence single-cell but not bulk projection patterns.

**Figure S3, related to Figure 2. Error sources of BRICseq. A**. Template switching is one error source in BRICseq. During PCR, a DNA strand being synthesized may use a new template during elongation, due to a shared sequence among all the templates. Details are discussed in Supplemental Note 3.1. **B,C**. Increasing the PCR volume from 25µL to 2mL dramatically decreased template switching rates. BRICseq was performed with samples from two brains, and 96 cubelets (48 from each) were pooled together for PCR and sequencing. **D**. In BL6-1, PCR was performed in a 12 mL system. Cubelets from a different BRICseq brain were added during PCR to estimate template switching rates. Increasing UMI threshold helped further reduce errors caused by template switching. **E**. Re-used barcode is another error source in BRICseq. If the same barcode is used more than once to infect multiple neurons, somata of some of the infected neurons are mis-interpreted as axons. Details are discussed in Supplemental Note 3.2. **F**. By simulating the process of sampling barcodes from the viral library, ratio of re-used barcodes can be estimated. More specifically, if we define each barcode's firstmax and secondmax as the highest and second highest UMI counts among all the cubelets, the joint distribution of (firstmax, secondmax) of re-used barcodes can also be calculated (Supplemental Note 5.5). Here we show the ratio of estimated re-used barcodes to total number of recovered barcodes in the (firstmax, secondmax) space. Black lines indicate axon threshold ($\theta_{axon}$, 20) and soma threshold ($\theta_{soma}$, 250) used to reduce re-used barcode errors. Neurons are included for analysis only when $\text{firstmax}>\theta_{soma}$ and $\text{secondmax}<\theta_{axon}$. **G**. Volcano plots of connection strengths and significance in BL6-1. To calculate p values, errors caused by template switching, re-used barcodes, and baseline contaminations are considered (Supplemental Note 3.3, 3.4).

**Figure S4, related to Figure 2. Comparing connection maps between animals. A**. Cubelet-to-cubelet connetion strengths in BL6-2. **B**. The method to infer brain area-to-brain area connections by weighted averaging for source and targets. Data in Figure 2C and Figure S11H are calculated in this way. **C**. Brain area-to-brain area connection maps in BL6-1 and BL6-2 inferred by weighted averaging. Figure 2C is generated based on them. **D**. The method to infer cubelet-to-brain area connections by weighted averaging. Data in Figure 2D,E are calculated in this way. **E-G**. The method to infer brain area-to-brain area connections by constrained optimization. **E**. Constrained optimization assumptions and methods. **F**. Brain area-to-brain area connection in BL6-1 and BL6-2 inferred by constrained optimization. **G**. Comparison of brain area-to-brain area connection between BL6-1 and BL6-2 inferred by constrained optimization (Pearson R = 0.7595, linear regression $p < 10^{-100}$).

**Figure S5, related to Figure 2. Normalize connection maps between animals by undersampling sequencing results. A.** Many factors including RNA extraction efficiency and sequencing depth could potentially affect the measured connection strength in different experiments. We therefore normalized BRICseq data before cross-brain comparisons (Supplemental Note 2.2) to counteract these effects. To do so, we assumed that the underlying distribution of barcode counts at infected somas are consistent across animals, and any variation in the observed distributions is caused by technical variation between experiments. We then undersampled the detected RNA molecules of a given brain to best approximate the soma barcode count distribution of another. As an illustration, the black line shows the histogram of the barcode counts at each soma of mouse BL6-2. The blue line shows the same histogram for mouse BTBR-2. Due to the smaller number of infected neurons in BTBR-2 and the resulting increased sequencing depth of this library, we detected on average higher barcode counts in BTBR-2 than BL6-2. The other three lines (cyan, green and yellow) indicate the results of undersampling mouse BTBR-2 with 3 different rates. **B**. The optimal undersampling rate is determined by minimizing the sum of squared errors of distributions of barcode counts at soma between BL6-2 and undersampled BTBR-2. In this example, to compare BTBR-2 to BL6-2, undersampling rate 0.31 is optimal.

**Figure S6, related to Figure 3. Function-connection relationship.** **A**. The behavior task. **B.** Scatter plot of input correlations and activity correlations. The input correlation is Pearson correlation of input to each pair of cublets. **C.** As the distance between a pair of cubelets has a strong effect on the connection strength, input correlation, and activity correlation, we decompose connection strengths, input correlations and activity correlations into distance-dependent components and distance-independent components (Supplemental Note 2.3). Here the relationship between distance-independent connection strengths/input correlations and activity correlations are plotted. **D-G**. The scatter plots of spontaneous/noise correlations versus connection strengths/input correlations. **D** and **F** show the original data; **E** and **G** show the data after removal of distance-dependent components. **B-G**. Red lines show medians.

**Figure S7, related to Figure 4. Gene expression-connection relationship. A,** Analysis pipeline. **B,C**. The bases and loadings of projection PCs in BL6-1. **D**. Comparisons between the predicted loadings with linear regression models using top 10 predictor genes and the observed loadings for projection PCs 1-10. **E**. For projection PCs 1-10, the Pearson correlation coefficients (Pearson r) between predicted loadings and observed loadings increase with the numbers of predictor genes. Error bars represent S.E.M.

**Figure S8, related to Figure 5. Anterograde tracing in C57BL/6J and BTBR mice**. **A.** Anterograde fluorescence tracing from right retrosplenial cortice. Black arrows indicate injection sites. Yellow arrows indicate projection sites. Images are shown in inverted grayscale.

**Figure S9, related to Figure 5. Connectopathies in BTBR mice**. **A**. Quantification of contralateral connection strengths in all 4 animals. Kruskal-Wallis test, $p < 10^{-100}$. Tukey multiple comparison test: n.s., $p > 0.05$; *, $p < 10^{-8}$. Error bars represent S.E.M. **B.** Distributions of nonzero commissural connections in BTBR mice. They were found exclusively in target cubelets close to the midline, and thus likely represented dissection error and contamination from the ipsilateral hemispheres.

**Figure S10, related to Figure 5. Additional analyses of 4 types of corticocortical projections. A**. Cumulative density distributions of connection strengths in the 4 types of connections in BL6-1. Projection strengths of zeroes are plotted as $10^{-3}$ (-3 in the log scale). **B.** Correlation between ipsilateral projection strength and contralateral projection strength for each target pair in heterotopic ipsi+ projections in BL6-1. **C**-**E**. The analyses of contralateral projections in BL6-1 when multiple comparison correction is not performed for the significance test. **C.** Cumulative density distributions of connection strengths. **D.** Piechart of positive projections in BL6-1. **E.** Correlation between ipsilateral projection strength and contralateral projection strength for each target pair in heterotopic ipsi+ projections.

**Figure S11, related to Figure 5. Additional analyses of cortical networks. A.** Distance-dependent connection strength and connection probability in BL6-1. The lines are fitting curves. (Left, double exponential fitting; right, single exponential fitting.) **B**. Histograms of output/input correlation between pairwise cubelets in BL6-1. Output(input) correlations are defined as Pearson correlations between output(input) from(in) pairs of cubelets. **C**. Distance-dependent output/input correlation in BL6-1. Red lines show medians. **D,E**. Distance-dependent connection strengths and output/input correlation in pairwise cubelets in BL6-2. Lines in **D** show fitting results (left, double exponential fitting; right, single exponential fitting), and red lines in **F** show medians. **F.** Left, distance-dependent connection strength and connection probability in BTBR (BTBR-1). The lines are fitting curves. (Left, double exponential fitting; right, single exponential fitting) **G**. Distance-dependent input/output correlation in BTBR (BTBR-1). The red lines show median input/output correlation. **H**. Comparison of ipsilateral brain area-to-brain area connection strengths between BL6-1 and BL6-2/BTBR (BTBR-1). Black: BL6-1 vs BL6-2, Pearson R = 0.7979, linear regression $p < 10^{-100}$; green: BL6-1 vs BTBR (BTBR-1), Pearson R = 0.6912, linear regression $p < 10^{-100}$.

**Figure S12, related to Figure 6. Additional analysis on network motifs. A**. Distribution of 2-node and 3-node motifs in cortical network in BL6-1 compared to randomly generated networks generated with the observed distance-dependent low-order properties, $RN_{dd}$ (Supplemental 4.4). *, $p < 0.001$. **B**. Fractions of 2-node and 3-node motifs in cortical networks in BL6-1. **C**. Distribution of 2-node and 3-node motifs in cortical network in BL6-2 compared to randomly networks generated with the observed global low-order properties, $RN_g$. *, $p < 0.001$ (in 10,000 simulation trials). **D.** Fractions of 2-node and 3-node motifs in cortical networks in BL6-2. **E**. Distribution of 2-node and 3-node motifs in cortical network in BTBR (BTBR-1) compared to randomly networks

$RN_g$. *, $p < 0.001$ (in 10,000 simulation trials). **F.** Fractions of 2-node and 3-node motifs in cortical networks in BTBR (BTBR-1). **G.** Clustering coefficients of cortical networks in BL6-1, BL6-2 and BTBR (BTBR-1) compared to random networks $RN_g$.

**Figure S13, related to Figure 6. Additional analysis on network modules. A**. In the clustering algorithm, a parameter, $\gamma$, can be tuned to adjust the size and number of modules (Supplemental Note 2.6). Here To determine $\gamma$, the original data are undersampled for 100 times, and the average number of modules and Rand index (inconsistency) are calculated for each $\gamma$ in BL6-1. $\gamma = 0.83$ was used. **B**. Connection ($M_c$)/input correlation ($M_{ic}$)/output correlation ($M_{oc}$) based modules in BL6-1. **C.** Comparison of $M_c$, $M_{ic}$, and $M_{oc}$ in BL6-1. **D**. As the distance between a pair of cubelets has a strong effect on the connection strengths, the original connection matrix can be decomposed into a distance-dependent connection matrix and a distance-independent connection matrix (Supplemental Note 2.6). Modules based on original ($M_o$), distance-dependent ($M_{dd}$), and distance-independent ($M_{di}$) connection matrices in BL6-1 are shown. **E**. Comparison of $M_o$, $M_{dd}$, and $M_{di}$ in BL6-1. $M_o$ is more similar to $M_{di}$. **F**. Modules clustered with different $\gamma$ in BL6-1. **G-H**. Connection/input correlation/output correlation-based modules in BL6-2. **I**. Modules clustered with different $\gamma$ in BL6-2. **J**. Connection-based modules, input correlation-based modules, and output correlation-based modules in BTBR (BTBR-1). Note there is one blank row where only axons but no somata were detected in any of its cubelets, probably due to missing of viral injections. **K**. Comparison of $M_c$ and $M_{ic}/M_{oc}$ in BTBR (BTBR-1). **L**. Modules clustered with different $\gamma$ in BTBR (BTBR-1).

### Supplemental Notes

#### Supplemental Note 1: Potential MAPseq artifacts are minimal and MAPseq efficiency is high

Below we discuss several classes of potential MAPseq artifacts.

**Degenerate and double barcode labeling**. In MAPseq, barcodes drawn from a high-diversity viral pool are used to uniquely label neurons. Ideally, every infected neuron would have a single, unique barcode. As discussed in detail in (Kebschull et al., 2016), there are two potential deviations from this ideal scenario: (i) multiple neurons per barcode, and (ii) multiple barcodes per neuron. The former, multiple neurons per barcode, i.e. re-used barcodes, is problematic as it leads to incorrect results. Because the probability that two neurons are infected by the same barcode is determined by the diversity of the barcode library and the total number of infected cells, we generated a high diversity viral library with over $8\times10^6$ barcodes for BRICseq (about $5\times10^4$ neurons). We also computationally inferred and calculated false positive projections caused by re-used barcodes (see details in Supplemental Note 3). The consistency between BRICseq and Allen Connectivity Atlas (Figure 2D,E) confirmed that the effects were minimum. The second scenario, multiple barcodes per neuron, has much less severe consequences. If a neuron expresses more than one barcode, BRICseq will overestimate the number of traced neurons. However, the relative abundance of each projection type and the bulk connection strength remain unchanged. In practice, we also aimed to infect neurons at close to 1 barcode per neuron by using appropriate volume and titer of Sindbis viruses.

**Non-uniform barcode transport**. In MAPseq, we interpret the number of barcodes from source cubelet X in target cubelet Y as a measure of the strength of projection from X to Y, analogous to GFP intensity. For this assumption to be valid, we must implicitly assume that barcode transport is uniform; in particular, we must assume that nearby and distal targets are equally filled. This assumption was rigorously validated in previous work (see Figure 2 in (Kebschull et al., 2016)).

**Fibers of passage**. Although the MAPseq carrier protein is derived from the synaptic protein neurexin (Kebschull et al., 2016), it does not exclusively target to presynaptic terminals. Thus MAPseq does not distinguish between synaptic connections (axon terminals) and fibers of passage. In this respect it is analogous to using GFP intensity in a conventional connectivity Atlas to measure the strength of the connection from the injection site to a target. To minimize potential confounds due to fibers of passage, we avoided white matter when dissecting cortical cubelets.

**MAPseq efficiency**. The efficiency of MAPseq has also been quantified in (Kebschull et al., 2016). By injecting red retrobeads into the olfactory bulb, and infecting the locus coeruleus with GFP-barcode Sindbis viruses, 91.4±6% of all barcodes from cells that projected to the olfactory bulb as determined by bead labeling also appeared to project to the bulb by sequencing (Figure S6 in (Kebschull et al., 2016)).

**End-to-end assessment of artifacts**. The agreement between BRICseq and the Allen Atlas reported in Figure 2D,E also confirmed that potential MAPseq artifacts above were minimal, consistent with previous validations (Chen et al., 2018; Han et al., 2018; Kebschull et al., 2016).

### Supplemental Note 2. Supplemental results and discussions

#### 2.1 Single cell projection data

In principle, by determining soma locations of individual barcodes, we were able to reconstruct projection patterns at single cell resolution. As an example, Figure S2A shows projection patterns of 712 cells in one source cubelet (cubelet #53 in PTLp); the cells are clustered with non-negative least squares-based sparse non-negative matrix factorization (cluster number is 10). Three of them are highlighted in dorsal views (Figure S2B). In the current data, individual cells displayed a wide range of node degrees (i.e. numbers of projecting target; Figure S2C). However, due to re-used barcodes (Supplemental Note 3.2), template switching (Supplemental Note 3.1) and inadequate sequencing depth (Figure S2D), further analyses regarding single cell projections were not performed with the current data.

#### 2.2 Normalize connection maps between animals by undersampling sequencing results

Many experimental factors including RNA extraction efficiency and sequencing depth could vary between individual experiments. For instance, due to variations of virus injections, the number of infected cells might vary between animals. A lower number of infected cells resulted in a lower count of total molecules, and thus an increase in sequencing depth (read per molecule), given the fact that the sequencing depth is generally low in BRICseq. In such cases, more barcode molecules (UMIs) in the axon and soma were sequenced per barcoded neuron, causing experimental biases. To compensate these variations and make different experimental results comparable, we sought a normalization method. We first assumed that the real distributions of molecule counts of barcode RNA at each neuron's soma (DOMCAS) were consistent between animals. We then reasoned that if we were able to undersample the sequencing result of a given experiment so that its DOMCAS matched another experiment, the data from these two experiments would be comparable (i.e., the same net efficiency of barcode detection). As an example, we undersampled the sequencing result of BTBR-2 to make it consistent with BL6-2. As shown in Figure S5A, the originally measured DOMCAS had a much longer tail in BTBR-2 (black) than BL6-2 (blue), due to a lower count of infected neurons and higher sequencing depth. By downsampling the BTBR-2 result, the DOMCAS was left-shifted (Figure S5A, gray lines). To find the optimal undersampling rate, we minimized the sum of squared errors of DOMCAS between undersampled BTBR-2 and BL6-2 (Figure S5B, optimal rate = 0.31). The data were pre-processed to normalize the net efficiency of barcode detection, and next used for further analyses. All the figures and calculations that compared connection maps between experiments were generated based on pre-processed data, including Figure 2C, Figure 5C-E, Figure S4C,F,G, Figure S9, Figure S10 and Figure S11H.

#### 2.3 Comparing BRICseq data with functional imaging data

To compare BRICseq data with functional imaging data, we trained animals with a perceptual decision making task (Figure S6A; see details in methods) (Musall et al., 2018), and calculated the reciprocal connection strength, input correlation (Pearson correlation of input vectors), activity correlation, noise correlation and spontaneous correlation for each pair of cubelets (Supplemental Note 5.9). Activity correlations (Figure 3C; Figure S6B), noise correlations (Figure S6F) and spontaneous correlations (Figure S6D) were all strongly correlated with reciprocal connection strengths or input correlations. As the distance between cubelets had a large effect on the connection strength (Figure S11A), input correlation (Figure S11C) and activity correlation (data not shown), we further removed distance-dependent components from connection strengths, input correlations, and activity correlations (Supplemental Note 5.9). The residual distance-independent components showed weaker, but still significant correlations (Figure S6C). Moreover, similar results were found between connection strength/input correlation and noise/spontaneous correlation (Figure S6E,G). The consistency between connection data and functional data not only validated BRICseq as a functionally relevant measure of cortical connectivity, but also suggested that intracortical connections are highly related to cortical activity.

#### 2.4 Input/output correlation

The input correlation (Pearson correlation of a pair of input vectors) and output correlation (Pearson correlation of a pair of output vectors) were calculated for each pair of cubelet. We found that both the input correlation and output correlation showed asymmetric distributions and no strongly negatively correlated input/output patterns were observed. This is consistent with sparseness of connections in the cortex. Furthermore, not surprisingly, both input correlation and output correlation decayed as the distance between two cubelets increased (Figure S11C,E), suggesting that proximal cubelets share more similar connection patterns than distal cubelets.

#### 2.5 Motif analysis

The statistical properties of a connected network can be decomposed into a series of mathematical terms that quantify increasingly complex motifs in the network, analogous to a Taylor expansion. Specifically, considering a binary directed network $\boldsymbol{G}$ (represented by its adjacent matrix), we define its following properties with increasing statistical orders:

1$^{st}$ order property: connection probability. Let $f_{1,1}(\boldsymbol{G})$ denote the connection probability in network $\boldsymbol{G}$.

2$^{nd}$ order properties: probability of 2-node motifs. There are 3 possible 2-node motifs: non-connected 2-node motif, uni-connected 2-node motif, and bi-connected 2-node motif, so (3-1=2) degrees of freedom are needed to describe 2$^{nd}$ order properties. Let $f_{2,1}(\boldsymbol{G})$, $f_{2,2}(\boldsymbol{G})$ and $1 - f_{2,1}(\boldsymbol{G}) - f_{2,2}(\boldsymbol{G})$ denote the probabilities of 2-node motifs in $\boldsymbol{G}$.

3$^{rd}$ order properties: probability of 3-node motifs. There are 16 possible 3-node motif, so (16-1=15) degrees of freedom are needed to describe 3$^{rd}$ order properties. Let $f_{3,1}(\boldsymbol{G})$ … $f_{3,15}(\boldsymbol{G})$ and $1 - \sum_{i=1}^{15} f_{3,i}(\boldsymbol{G})$ denote the probabilities of 3-node motifs in $\boldsymbol{G}$.

…

n$^{th}$ order properties: probability of n-node motifs. Let $M_n$ denote the number of n-node motifs, then we need ($df_n = M_n - 1$) degrees of freedom to describe n$^{th}$ order properties.

Let $f_{n,1}(\boldsymbol{G}) \dots f_{n,df_n}(\boldsymbol{G})$ and $1 - \sum_{i=1}^{df_n} f_{n,i}(\boldsymbol{G})$ denote the probabilities of n-node motifs in $\boldsymbol{G}$.

Given n^th order properties of $\boldsymbol{G}$, we are able to infer the probability distribution of $\boldsymbol{G}$, $P_n(\boldsymbol{G})$, using the maximum entropy principle (Jaynes, 1957):

$$P_n(\boldsymbol{G}) = \frac{1}{\ln Z_n} \exp\left(-\sum_{i=1}^{df_n} \mu_{n,i} f_{n,i}(\boldsymbol{G})\right)$$

, where

$$Z_n = \sum_{\boldsymbol{G}} \exp\,(-\sum_{i=1}^{df_n} \mu_{n,i} f_{n,i}(\boldsymbol{G}))$$

$$\frac{\partial Z_n}{\partial \mu_{n,i}} = < f_{n,i}(\boldsymbol{G}) >$$

. Here $< f_{n,i}(\boldsymbol{G}) >$ represents observed probability of a given n-node motif.

Define

$$I_n = \begin{cases} \sum_{i=1}^{df_n} \mu_{n,i} f_{n,i}(\boldsymbol{G}) - \sum_{i=1}^{df_{n-1}} \mu_{n-1,i} f_{n-1,i}(\boldsymbol{G}) & (n > 1) \\ \mu_{1,1} f_{1,1}(\boldsymbol{G}) & (n = 1) \end{cases}$$

, then we get an expansion form of $P_n(\boldsymbol{G})$:

$$P_n(\boldsymbol{G}) = \frac{1}{\ln Z_n} \exp\big(-(I_1 + I_2 + \cdots + I_{n-1} + I_n)\big)$$

. In the above formula, if we only take the first $k$ terms and recalculate the partition function $Z$, we get $P_k(\boldsymbol{G})$, i.e., the probability distribution of $\boldsymbol{G}$ given only $k^{th}$ order statistical properties. Studying $P_k(\boldsymbol{G})$ with increasing $k$ reveals the effect of higher order statistical properties on $\boldsymbol{G}$. In particular, the second and third terms represent the interactions between pairs and triplets of elements, respectively.

We first studied 2-cubelet motifs in the C57BL/6J cortical network. Random networks, $RN_g$ were generated based on observed global first-order properties ($< f_{1,1}(\boldsymbol{G}) >$). In our C57BL/6J cortical network, the probability that a pair of cubelets $x$ and $y$ was reciprocally connected was greater than predicted by the null hypothesis (Figure 6A; Figure S12B-D). Three-cubelet motifs were also highly non-random, with a tendency for densely connected motifs to be particularly overrepresented (Figure 4G; Figure S12B-D), compared to $RN_g$, random networks generated based on observed global second-order properties (i.e. probabilities of 2-cubelet motifs, $< f_{2,i}(\boldsymbol{G}) >, i = 1{,}2$). The most under-represented motif was a unidirectional cycle, similar to what has been reported at the cellular level (Dechery and MacLean, 2018). Consistent with 3-cubelet motif statistics, the observed clustering coefficient was high compared to random networks (Figure S12G). Interestingly, the distribution of 3-cubelet motifs was strikingly similar to statistics of connections among single neurons in the rat visual cortex (Song et al., 2005), suggesting that a common rule might govern the organization of neural circuits at both microscale (intra-neuron) and mesoscale (intra-area) levels. These analyses reveal that the network architecture was highly structured, deviating sharply from simple random connectivity.

As the distances between cubelets may affect the probability of connections or motifs, we also considered random networks generated with the observed distance-dependent low-order properties, $RN_{dd}$. Comparing $RN_{dd}$ to observed networks (Supplemental Note 5.8), similar overrepresented/underrepresented motifs were found (Figure S12A).

**2.6 Module analysis**

Previous analyses of the connectivity between cytoarchitecturally defined brain areas (Harris et al., 2018; Zingg et al., 2014) revealed "modules"— regions of the brain within which connections are dense, and which may reflect functional units. Because the basic unit in BRICseq is a cubelet, defined by dissection without regard to functionally defined regions, we wondered whether similar modules would emerge, or whether we could reveal structure within classical brain areas that were previously obscured by their labeling as one homogeneous area.

To analyze modules of the ipsilateral cubelet-to-cubelet connection matrix, we utilized a community structure-finding algorithm (Rubinov and Sporns, 2010). In the algorithm, a resolution parameter, $\gamma$ can be tuned to get smaller/more or larger/fewer modules. To choose a proper $\gamma$, we undersampled from all the projection neurons, and used the algorithm to find modules. The optimal $\gamma$ was chosen so that the Rand index (inconsistency) was low and the average number of modules was stable (Figure S12A; Supplemental Note 5.7).

In the C57BL/6J mouse #BL6-1, with the optimal $\gamma$=0.87, we recovered four major modules (Figure 6C,D), of which module 1 belonged to visual-auditory areas, modules 2 and 3 belonged to somatosensory/motor areas, and module 4 belonged to anterior cingulate/retrosplenial areas. Interestingly, the two modules belonging to somatosensory/motor areas were not clustered according to brain areas defined in the Allen atlas (i.e. SSp, SSs, MOp, MOm), but were clustered according to the represented body parts. Roughly, module 2 corresponded to somatosensory and somatomotor areas associated with sensation and movement of limbs, trunk and whiskers, whereas module 3 corresponded to areas associated with mouth and nose. Interestingly, the modules obtained by this analysis of connectivity closely match those obtained by brain-wide calcium imaging and clustering, but not the partitioning based on cytoarchitecture (Vanni and Murphy, 2014).

In addition to the connection matrix, we could perform a similar analysis on the input (or output) correlation (Pearson correlation between input to a pair of cubelets or between output from a pair of cubelets) matrix. The modular organization of these matrices was similar to that of the connections themselves (Figure S13B,C), suggesting that inter-connected modules tend to receive similar inputs and send similar outputs.

As the distance between two cubelets strongly affected their connection strength (Figure S11A), we asked whether the observed connection modules in the connection matrix were a result of simple distance dependence, or due to specific connection patterns between brain areas. To address this question, we generated a distance-dependent connection matrix ($M_{dd}$) with observed average connection strengths at various distances, and performed clustering analysis (Figure S13D, middle; Figure S13E, left). A distance-independent connection matrix ($M_{di}$) was also generated by subtracting the distance-dependent connection matrix from the original connection matrix, and analyzed in a similar way (Figure S13D, right; Figure S13E, right). The original connection modules were more similar to the distance-independent connection modules, suggesting the cortical modules were not simply organized with a distance-dependent rule, but reflected specific connection patterns between certain brain areas.

We also performed module analysis with various $\gamma$ (Figure S13F). With lower $\gamma$, we found fewer modules, which consisted of one or more modules that were determined with $\gamma$=0.83. However, we failed to detect fine modules from the connection network with higher $\gamma$, probably due to limited spatial resolution of dissected cubelets (300μm×1mm×1mm).

Similar analysis was done in mouse BL6-2 (Figure S13G-I). With the optimal resolution parameter $\gamma$=0.8, five major modules were recovered: two somatosensory-somatomotor area modules (similar with the two modules in BL6-1), a visual-auditory area module, an anterior cingulate area module, and a retrosplenial area module. The global modular organizations between BL6-1 and BL6-2 were similar, and the subtle differences may result from limited spatial resolution and cubelet dissection variations.

#### 2.7 Analysis of contralateral projections

Commissural connections play important roles in regulating a variety of behaviors, and disruption of these projections might play a role in autism and other neurological dysfunction (Paul, 2011; Sperry et al., 1969). However, the network properties of commissural projections are much less studied than their ipsilateral counterparts. We therefore examined the structure of commissural connections in our dataset. In the C57BL/6J mouse #BL6-1, homotopic projections, i.e. projections from one area of cortex to the corresponding area in the other hemisphere (Figure 5D, left), are the most likely to be positive (Figure 5D, middle; Figure S10A, blue; 47.2±0.6% of all possible homotopic projections are positive), and made up a substantial fraction of all commissural projections (36.9±0.6% of all non-zero commissural projections are homotopic), consistent with previous reports (Bota et al., 2015). The remaining projections were heterotopic projections from one area of cortex to a non-corresponding area in the contralateral hemisphere, but not to its corresponding contralateral area. To understand the structure of these projections, we further divided heterotopic projections into those with projections to both the ipsi and contralateral versions of a target area (heterotopic ipsi+) and those that projected only to the contralateral version of one area, but not the ipsilateral one (heterotopic ipsi-; Figure 5D, left). 71.1±4.6% of positive heterotopic projections were of the ipsi+ kind. Heterotopic ipsi- projections accordingly made up only 18.2%±2.7% of the all commissural projections. Our findings therefore support a largely symmetric model of the mouse cortex, where a given area in one hemisphere often directly projects to its corresponding area in the contralateral hemisphere, and moreover preferably projects to both ipsi- and contralateral versions of other target areas.

Heterotopic ipsi+ commissural projections constituted a substantial fraction of total commissural projections and they represented bifurcated projections to two hemispheres. To further study these projections, we defined the source area S, the ipsilateral target area T, and the contralateral target area T'. Note T and T' are homotopic, according to the definition. For all the positive heterotopic ipsi+ commissural projections, the correlation between S-T projection strength and S-T' projection strength was weak (Figure S10B).

All the previous analysis of commissural projections was based on projections that have passed the significance test with multiple comparison. As commissural projections were usually weaker than association projections (Figure 5E), the false negative rate for commissural projections might be higher. To examine how the false negative rate may affect results, we also did parallel analysis with projection data that have passed the significance test but without multiple comparison. Homotopic commissural projection was still the major commissural projection type, while the number of heterotopic ipsi+ commissural projection increased (Figure S10C,D). For heterotopic ipsi+ projections, weak but significant correlation was observed between the association projection strength and the commissural projection strength (Figure S10E).

**2.8 The BTBR brain**

In the BTBR brain (BTBR-1), the connection strength and the input/output correlation between a pair of cubelets decreased as the distance increased (Figure S11F,G), similar to what was observed in BL6-1 and BL6-2.

By performing module analysis, we found modules in the BTBR-1 connection matrix ($M_c$; Figure S13L). Four major modules were found in the connection matrix at the optimal resolution parameter $\gamma$=1.1 (Figure S13J, left): three modules in the somatosensory-somatomotor area (roughly corresponding to orofaciophryngeal, upper limb, lower limb – whisker areas respectively), and one module in the anterior cingulate-retrosplenial-visual area. The differences in the somatosensory-somatomotor areas between the BTBR brain and C57BL/6J brains might be due to limited spatial resolution and cubelet dissection variations. The failure to get the visual-auditory area module is likely explained by injection artifacts: In the BTBR brain, probably due to lack of corpus callosum, the two cortical hemispheres are physically separated much more rostrally than a C57BL/6J mouse. Thus, some cortical brain areas including the auditory cortex and part of the visual cortex are more lateralized and difficult to be targeted by viral injection from the dorsal surface. Actually, very few, if any, somata were found in these areas from the sequencing results. As cubelets with too few infected cells (less than 50) were excluded for analysis, the visual-auditory area was not recovered as a module, as seen in BL6-1. We also performed module analysis with the input correlation matrix ($M_{ic}$; Figure S13J, middle) and the output correlation matrix ($M_{oc}$; Figure S13J, right), and the results were similar to modules in the connection matrix (Figure S13K).

Topological properties of the ipsilateral cubelet-to-cubelet connectivity network was also examined in the BTBR brain (BTBR-1). All the results were very similar to C57BL/6J brains (Figure S12E-G). Briefly, among all the 2-node motifs, bidirectional connections were overrepresented; the clustering coefficient was significantly higher than random networks generated with the same second-order properties; the distribution of 3-node motifs was highly non-random, and densely connected motifs were overrepresented. The results suggested that these topological properties of the ipsilateral connection network were not disrupted in the BTBR brain.

All the results observed in BTBR-1 were also similarly seen in BTBR-2 (data not shown).

**Supplemental Note 3: Sources of errors and calculation of cubelet-to-cubelet connections in BRICseq**

**List of variables in Supplemental Notes 3**

| | |
|---|---|
| $l_1$ | Number of molecules in cubelet 1 |
| $l_2$ | Number of molecules in cubelet 2 |
| $c$ | Template switching rate constant |
| $h_{12}$ | Number of cubelet 1-cubelet 2 hybrid molecules |
| $N(i)$ or $N_1(i)$ | Number of projection neurons (type 1 neurons, Supplemental Note 3.2) residing in cubelet $i$ |
| $N_3(i)$ | Number of type 3 neurons (Supplemental Note 3.2) residing in cubelet $i$ |
| $N_t$ | Total number of barcodes in the BRICseq result (type 1-4, Supplemental note 3.2) |
| $N_{re}$ | Total number of re-used barcodes |
| $n(i,j)$ | Total number of molecules of $j$th neuron in $i$th cubelet (soma molecules + all axon molecules) |
| $n_{soma}(i,j)$ | The number of soma molecules of $j$th neuron in $i$th cubelet |
| $n_{axon}(k)$ | The number of axon molecules detected in $i$th cubelet |
| $p(i,j,k)$ | The probability that molecules of $j$th neuron in $i$th cubelet were detected in $k$th cubelet due to template switching |
| $m_k$ | Number of error molecules from neurons in experimental cubelets that were detected in $k$th control cubelet due to template switching |
| $b$ | Number of error molecules in each cubelet due to baseline contamination |
| $P_\theta(i,j,k)$ | The probability that > $\theta$ error molecules of $j$th neuron in $i$th cubelet were detected in $k$th cubelet due to template switching |
| $r_{ts}(i,k)$ | The average probability that a false projection from a neuron in $i$th cubelet to $k$th cubelet was detected due to template switching |
| $r_{re}(i,k)$ | The average probability that a false projection from a neuron in $i$th cubelet to $k$th cubelet was detected due to re-used barcodes |

| | |
|---|---|
| $r_{ba}(i,k)$ | The average probability that a false projection from a neuron in $i$th cubelet to $k$th cubelet was detected due to baseline contamination |
| $v_{ik}$ | p value (false positive probability) of cubelet $i$-to-cubelet $k$ projection |
| $N_{pro}(i,k)$ | Observed number of neurons in cubelet $i$ that projected to cubelet $k$ |
| $C$ | Cubelet-to-cubelet connection matrix |

There are two major error sources that affected BRICseq data: template switching and re-used barcodes. In this section, we first describe how these error sources cause false positive results in BRICseq and how we reduce these effects experimentally. Next, we establish quantitative models about false positive projections in the experimental data. Finally, according to the mathematical model of false positive projections, we correct the observed cubelet-to-cubelet connection strength, and calculate a p-value for each cubelet-to-cubelet connection, indicating its significance level.

The following terms are defined before further discussion. 1) Barcode: a barcode is a unique 32nt sequence delivered by the Sindbis virus. One barcode theoretically corresponds to a neuron. 2) Molecule: here a molecule is defined as a unique BC-CSI-UMI (32nt + 8nt + 12nt) sequence. A molecule should correspond to a single RT product. Due to barcode amplification in a neuron, one barcode has multiple molecules. 3) Molecule copy: a molecule copy is defined as a final product after PCR. A large number of molecule copies are generated from one molecule during PCR. 4) Read: reads are the sequencing product. Not considering sequencing errors, all the reads constitute a subset of all the molecule copies.

#### 3.1 Template switching

##### 3.1.1 How template switching affects BRICseq and experimental solutions

Template switching may occur when DNA templates share a common sequence during PCR (Figure S3A). In BRICseq, cDNA from all the cubelets was pooled together for PCR, and they all shared a common RT primer annealing sequence. The hybrid products of template switching caused barcode molecules to appear in erroneous cubelets (in Figure S3A, BC2 is detected in cubelet 1 due to template switching). Template switching is usually considered to be rare, and might be corrected by setting a read threshold for molecules (Kebschull and Zador, 2015). However, low sequencing depth disabled the use of read threshold to remove error molecules. Moreover, as molecules of a barcode in a soma usually outnumbered molecules in axons by ~100 fold, template switching molecules might constitute a large proportion in axon barcodes, albeit rare compared to total molecules. Thus, template switching had a significant influence in measuring projection strengths in BRICseq.

As DNA concentration is a major factor determining the template switching rate, we proposed we could reduce template switch molecules by increasing the PCR volume. To systematically evaluate template switching and test our hypothesis, we designed an experiment to perform BRICseq from two brains. We injected similar amounts of barcoded viruses into two animals, collected cubelets, and performed RT from individual cubelets. Then single-strand DNA molecules were pooled (48 cubelets from each animal, 96 in total) for second-strand synthesis, PCR and sequencing. Thus 'inter-brain' projection molecules reflected template switching. To measure the effect of DNA concentration on template switching, the same sample was separated to perform PCR either in a 25 µL volume or in a 2 mL volume. In the 25 µL PCR experiment, a large number of molecules that were detected in both brains ('inter-brain' molecules) as well as stripe-like patterns in the barcode heatmap indicated a high rate of template switching (Figure S3B, left). By increasing PCR volume to 2 mL, 'inter-brain' molecules were dramatically decreased (Figure S3B, right). The rate of template switching could be further reduced by raising the UMI threshold that was used to determine a real projection (Figure S3C). In addition to the high reaction volume, we also set the PCR extension time in each cycle to 2min to reduce incompletely elongated products, another possible source of template switching.

To reduce template switching, we chose to perform the final PCR in 12 mL volume for BRICseq experiments BL6-1, BL6-2, BTBR-1 and BTBR-2. While Sindbis viruses harboring barcode library 2 were used to label experimental animals, we also injected Sindbis viruses harboring barcode library 1 into a few brain areas in a separate animal. After RT and second-strand synthesis, DNA molecules from experimental animals (261 cubelets in BL6-1, 262 cubelets in BL6-2, 258 cubelets in BTBR-1, and 262 cubelets in BTBR-2) were mixed with DNA molecules from library 1 virus-injected control animals (21 cubelets in BL6-1 control, 12 cubelets in BL6-2 control, 12 cubelets in BTBR-1, and 16 cubelets in BTBR-2) for PCR and sequencing, so the number of 'inter-brain' projection molecules was an internal measurement of template switching. In BL6-1, when we set UMI threshold to 1 (i.e. a projection was positive when its UMI count was greater than 1), 2004 out of 63107 barcodes were detected in the control brain (Figure S3D). Similar results were also found in other animals (data not shown).

#### 3.1.2 Calculating false-positive projections caused by template switching

With PCR volume = 12mL and UMI threshold = 1, the probability that a barcode was detected in a non-projecting cubelet due to template switching on average was reasonably low ($\frac{2004}{63107\times 21}$< 1%). To further determine whether a bulk projection was significant, we calculated the distribution of false positive projections caused by template switching. The calculations included the following steps:

1. Determine the template switching coefficient by linear regression.
2. Determine the probability that a neuron in source cubelet $i$ had a false positive projection to target cubelet $j$.
3. Determine the distribution of the number of neurons in source cubelet $i$ that false positively 'projected' to target cubelet $j$.

**Step 1. Determine the template switching coefficient by linear regression.**
First consider a general scenario. Let $l_1$ denote the number of molecules in cubelet 1, and $l_2$ denote the number of molecules in cubelet 2. If we pool these molecules to perform PCR, we assume the number of hybrid molecules after PCR $h_{12}$ can be written as:

$$h_{12} = 2cl_1l_2 \quad (1)$$

, where $c$ is called template switching rate constant, and should be dependent on the total number of initial molecules, PCR cycle number and PCR volume. As we pooled all the samples together for PCR, $c$ was a constant in one BRICseq experiment.

Specifically, in BRICseq, let $N(i)$ denote the number of neurons in cubelet $i$, $n(i,j)$ denote the number of molecules (including both soma molecules and axon molecules) for the $j$th neuron in cubelet $i$, $n_{soma}(i,j)$ denote the number of soma molecules for the $j$th neuron in cubelet $i$, and $n_{axon}(i)$ denote the number of axon molecules detected in cubelet $i$. The probability that the $j$th neuron in cubelet $i$ had a false positive molecule in cubelet $k$, $p(i,j,k)$ was:

$$p(i,j,k) = cn(i,j)\left(\sum_{l=1}^{N(k)} n_{soma}(k,l) + n_{axon}(k)\right) \quad (2)$$

.

In order to estimate the template switching coefficient $c$ in Eq. (2), we calculated the number of 'inter-brain' projection molecules as the ground truth of template switching molecules. If we considered template switching across two brains, then the number molecules that were from neurons residing in the experimental brain and found in the control brain cubelet $k$, $m_k$ was:

$$m_k = c\sum_{\substack{i\ in\\ exp.}}\sum_{j=1}^{N(i)} n(i,j)\,(\sum_{l=1}^{N(k)} n_{soma}(k,l) + n_{axon}(k)) \quad (3)$$

, where $i$ visited all the cubelets in the experimental brain and $j$ visited all the neurons in each experimental brain cubelet.

In the real experiment, there was an extra baseline contamination term (this term can also be inferred from molecules in additional control cubelets from a brain without viral injection), so Eq (3) was modified as:

$$m_k = c\sum_{\substack{i\ in\\ exp.}}\sum_{j=1}^{N(i)} n(i,j)\,(\sum_{l=1}^{N(k)} n_{soma}(k,l) + n_{axon}(k)) \quad (4)$$

, where $b$ was the baseline contamination constant.

In Eq. (4), the term $\sum_{\substack{i\ in\\ exp.}}\sum_{j=1}^{N(i)} n(i,j)$ is equal to the total amount of barcode molecules in the experimental brain, the term $\sum_{l=1}^{N(k)} n_{soma}(k,l) + n_{axon}(k)$ is equal to the total amount of barcode molecules in the control brain cubelet $k$, and $m_k$ is equal to number of library-2 barcode molecules in the control brain cubelet $k$. As all these numbers were known, we were able to use a linear regression model to fit Eq. (4) to estimate $b$ and $c$. As an example, in BL6-1, we got:

$$c = 1.12\times10^{-11}$$
$$b = 3.90\times10^{3}$$

.

**Step 2. Determine the probability that a neuron in source cubelet $\boldsymbol{i}$ had a false positive projection to target cubelet $\boldsymbol{j}$.**

With estimated $c$ and $b$, we could predict intra-brain template switching probability, $p(i,j,k)$ with Eq. (2) when $i$ and $k$ were both from the experimental brain. However, as we further filtered the data by setting a UMI threshold $\theta$ (Figure S3D), a false-positive projection was detected only when at least $(\theta + 1)$ template switching molecules from a given neuron to a given cubelet were seen. Let $P_\theta(i,j,k)$ denote the probability that the $j$th neuron in cubelet $i$ falsely projected to cubelet $k$ with UMI threshold = $\theta$, then according to Poisson distribution, we had

$$P_\theta(i,j,k) = \sum_{l=\theta+1}^{\infty} e^{-p(i,j,k)} \frac{p(i,j,k)^l}{l!} \quad (5)$$

.

As an approximation, we only calculated the first three terms when $\theta = 1$, as $p(i,j,k)$ was small enough and false projections with high UMI counts were extremely rare. We got:

$$P_1(i,j,k) \approx e^{-p(i,j,k)} \frac{p(i,j,k)^2}{2!} + e^{-p(i,j,k)} \frac{p(i,j,k)^3}{3!} + e^{-p(i,j,k)} \frac{p(i,j,k)^4}{4!}$$
$$\approx \frac{p(i,j,k)^2}{2!} + \frac{p(i,j,k)^3}{3!} + \frac{p(i,j,k)^4}{4!} \quad (6)$$

. With Eq. (6), we were able to calculate the probability that a given neuron in cubelet $i$ falsely ‘projected’ to cubelet $k$.

**Step 3. Determine the distribution of the number of neurons in source cubelet $\boldsymbol{i}$ that false positively ‘projected’ to target cubelet $\boldsymbol{j}$.**

In step 2, we were able to determine the probability that a given neuron in cubelet $i$ that falsely ‘projected’ to cubelet $k$. As cubelet $i$ consisted of $N(i)$ neurons, and each neuron had a different template switching probability ($P_1(i,j,k)$ is different for each $j$), the total number of $i$-to-$k$ false-positive neurons caused by template switching obeyed a Poisson binomial distribution. Note it was neither a Poisson distribution nor a binomial distribution, but a distribution of the sum of Bernoulli trials with different probabilities.

To calculate the distribution of the number of false positive projection neurons, we sought to calculate the Poisson binomial cumulative probability distribution. In BRICseq, there were over 30000 possible cubelet-to-cubelet projections, and for each of these projections, there were 500~1000 cells in the source cubelet (corresponding to 500~1000

Bernoulli trials). To our knowledge, there does not exist a fast and precise way to calculate the cumulative probability of the Poisson binomial distribution for each cubelet-to-cubelet projection. Particularly, when multiple comparison correction was considered, the p value was as small as $0.05/36018 \approx 1.66\times10^{-6}$; even for Monte-Carlo methods, a large number of simulation trials are required. Thus, we chose to use binomial distributions to approximate Poisson binomial distributions, assuming the probability of any given neuron in cubelet $i$ falsely projected to cubelet $k$, $r_{ts}(i,k)$, was the mean probability over all the neurons in cubelet $i$:

$$r_{ts}(i,k) = \frac{\sum_{j=1}^{N(i)} P_1(i,j,k)}{N(i)} \quad (7)$$

. Thus, the number of neurons in source cubelet $i$ that false positively 'projected' to target cubelet $k$ due to template switching was modeled as a binomial distribution with $N(i)$ experimental trials and probability of $r_{ts}(i,k)$.

Note when the required p value was not too small (for example, p = 0.05, without multiple comparison), we used Monte-Carlo method (10000 trials each) to estimate the cumulative probability of the Poisson binomial distribution for each cubelet-to-cubelet projection.

To summarize, template switching could be a detrimental error source when DNA concentration during PCR is high and sequencing depth is low. By using a large volume of the reaction system for PCR, setting a UMI threshold, and rejecting false positive projections, we have greatly reduced template switching errors to a very low level.

#### 3.2 Re-used barcodes

##### 3.2.1 How re-used barcodes affects BRICseq

To scale up MAPseq, it is crucial to use a barcode library with a sufficiently high diversity. Otherwise, the same barcode might label two (or more) different cells causing misinterpretation of the data (Figure S3E). The rate of re-used barcodes was determined by barcode diversity and the total number of infected neurons. In BRICseq for BL6-1, BL6-2 and BTBR-1, the diversity of the barcode library was no less than $8.26\times10^{6}$, according to the viral library sequencing result (note as we used a different viral library with much higher diversity for BTBR-2, re-used barcodes in BTBR-2 are ignorable). However, the total number of neurons expressing barcodes was much higher than the number of recovered neurons (~50000) due to a large number of 'non-projection' neurons. For example, in BL6-1, over 600000 'non-projection neurons' were recovered. Some of these 'non-projection' neurons might belong to local inhibitory or excitatory neurons, but a large number of them expressed RNA barcodes at very low levels. It was likely that due to variations of RNA expression levels, some projection neurons expressed very small amount of RNA barcodes, which couldn't be efficiently trafficked to axon terminals. These low expressed barcodes were almost all in the right cortical cubelets (injection site), and usually fewer than 20 molecules were detected in somata (cubelets with the highest molecule abundance), and no molecules above the UMI threshold (=1) were detected in axons (other cubelets). Moreover, these barcodes were also found in the viral library, suggesting they were unlikely due to sequencing errors. Although these 'non-projection' neurons were not included for data analysis, they might harbor re-used barcodes shared with other projection neurons, resulting in false projections (Figure S3E).

#### 3.2.2 Calculating false-positive projections caused by re-used barcodes

To quantify errors caused by re-used barcodes and remove them from connection results, we first set an additional set of thresholds to reduce re-used barcode errors. We nexted estimated the rate of re-used barcodes based on the thresolds, and determined the distribution of false positive projection neurons caused by re-used barcodes.

**Step 1. Reduce re-used barcodes by thresholding**

For each barcode, we defined its firstmax and secondmax as the highest and second highest abundance among all the cubelets. If a barcode corresponded to one neuron, then its firstmax was the count of molecules in its soma, and its secondmax was the count of molecules in its strongest axon. If a barcode was used in two neurons, then firstmax and secondmax were the highest two of UMI counts in two somata and two strongest axons. As the molecules in somata statistically outnumbered molecules in axons, secondmax of a re-used barcode was likely to be the amount of molecules in one of the two somata. According to this, we reasoned that re-used barcodes might have distinct distribution in the (firstmax, secondmax) space from barcodes used only once. To quantify this, we simulated the barcode sampling process (details in Supplemental Note 5.5, we modeled viral infection as a process where neurons randomly sampled barcodes from the barcode library), and calculated the number of re-used barcodes in the (firstmax, secondmax) space, given the observed joint distributions of (firstmax, secondmax) and the known barcode library. The ratio of simulated re-used barcodes to the total barcodes was plotted in the (firstmax, secondmax) space (Figure S3F). Not surprisingly, a higher ratio of re-used barcode was present close the diagonal line in the (firstmax, secondmax) space.

We next set a soma threshold (=250) and an axon threshold (=20) (Figure S3F), and defined 4 types of barcodes according to the thresholds:

Type 1 barcode: firstmax > soma threshold AND secondmax > UMI threshold AND secondmax < axon threshold.

Type 2 barcode: firstmax > soma threshold AND secondmax ≤ UMI threshold.

Type 3 barcode: firstmax < axon threshold AND firstmax > UMI threshold.

Type 4 barcode: secondmax > axon threshold.

To reduce the effect of re-used barcodes, we only included type 1 barcode for projection pattern analysis. Based on simulation results, in BL6-1, 8.77% of type 1 barcodes were re-used barcodes (8.27% in BL6-2 and 8.62% in BTBR-1). As there were 115 cubelets in the injection site of BL6-1, if a source cubelet and a target cubelet were both in the injection site (right hemisphere), then the probability of a type 1 neuron in the source cubelet that falsely projected to the target cubelet was on average $\frac{8.77\%}{115} \approx 0.0763\%$, which was reasonably low.

**Step 2. Distribution of false positive projection neurons caused by re-used barcodes**

To quantify false positive projection neurons caused by re-used barcodes for each cubelet-to-cubelet connection, we calculated $r_{re}(i,k)$, the probability that a type 1 neuron in cubelet $i$ that falsely projected to cubelet $k$ due to re-used barcodes. In BL6-1, for example, because a re-used type 1 barcode could only occur when a type 1 or type 2 neuron in the source cubelet and a type 3 neuron in a target cubelet shared the same barcode, we could estimate $r_{re}(i,k)$ with:

$$r_{re}(i,k) = \frac{8.77\% * N_3(k)}{\sum_{l\ in\ all} N_3(l)} \quad (8)$$

, where $N_3(k)$ represents the number of type 3 barcodes in cubelet $k$. Thus, the number of neurons in source cubelet $i$ that false positively 'projected' to target cubelet $k$ due to re-used barcodes was modeled as a binomial distribution with $N(i)$ experimental trials and probability of $r_{re}(i,k)$.

To conclude, with the viral barcode diversity used in BL6-1, BL6-2 and BTBR-1, the probability of re-used barcodes cannot be ignored. By setting thresholds for soma and axon identification, and rejecting false positive projections, we have removed most of errors due to re-used barcodes. In BTBR-2, a new viral barcode library with over $2.7\times10^8$ was used and this problem was completely circumvented.

#### 3.3 Calculating cubelet-to-cubelet connection strength

The projection strength from a source cubelet to a target cubelet was defined as the total count of UMIs in the target cubelet from all the neurons residing in the source cubelet divided by total number of projection neurons in the source cubelet. Considering the projection from cubelet $i$ to cubelet $k$, let $N(i)$ denote number of projection neurons in cubelet $i$ and $UMI(i,j,k)$ denote the UMI count in cubelet $k$ from $j$th neuron in cubelet $i$, then the UMI count in cubelet $k$ from an average neuron in cubelet $i$, $UMI(i,*,k)$ could be written as:

$$UMI(i,*,k) = \frac{\sum_{j=1}^{N(i)} UMI(i,j,k)}{N(i)} \quad (9)$$

. However, noise caused by template switching, re-used barcodes, and baseline contaminations could also contribute to $UMI(i,*,k)$. The noise level of the $i$-to-$k$ projection, $Noise(i,k)$, was calculated as:

$$Noise(i,k) = UMI_{ts}(i,*,k) + r_{re}(i,k) * UMI_{type3}(k) + r_{ba}(i,k) * UMI_{ba} \quad (10)$$

, where $UMI_{ts}(i,*,k)$ is the expected UMI count in cubelet $k$ from an average neuron in cubelet $i$ due to template switching, $UMI_{type3}(k)$ is the average UMI count of type 3 neurons in cubelet $k$ (after UMI thresholding), $UMI_{ba}$ is the average UMI count of a barcode in cubelets from the uninjected control brain (baseline contamination, after UMI thresholding), and the $r_{ba}(i,k)$ is the probability that a neuron in cubelet $i$ falsely projected to cubelet $k$ due to baseline contaminations, (estimated from non-injected control cubelets). These three terms corresponded to the template switching noise, re-used barcode noise, and baseline contamination noise. Particularly, $UMI_{ts}(i,*,k)$ was calculated with:

$$UMI_{ts}(i,*,k) = \frac{\sum_{j=1}^{N(i)} \sum_{l=\theta+1}^{\infty} l e^{-p(i,j,k)} \frac{p(i,j,k)^l}{l!}}{N(i)}$$
$$\approx \frac{2e^{-p(i,j,k)} \frac{p(i,j,k)^2}{2!} + 3e^{-p(i,j,k)} \frac{p(i,j,k)^3}{3!} + 4e^{-p(i,j,k)} \frac{p(i,j,k)^4}{4!}}{N(i)}$$
$$\approx \frac{\frac{p(i,j,k)^2}{1!} + \frac{p(i,j,k)^3}{2!} + \frac{p(i,j,k)^4}{3!}}{N(i)} \quad (11)$$

. The projection strength from cubelet $i$ to cubelet $j$, $C(i,j)$ was then calculated with:

$$C(i,k) = \max\{UMI(i,*,k) - Noise(i,k), 0\} \quad (12)$$

.

In addition to calculate the projection strength, we also calculated p value for each cubelet-to-cubelet connection, as noted in Supplemental Note 3.4.

**3.4 Calculating p values**

In addition to removing the noise estimate from the projection strength, we also calculated the p value for each cubelet-to-cubelet projection. For a source cubelet $i$ and a target cubelet $k$, we calculated the probability that a neuron in cubelet $i$ falsely projected to cubelet $k$ due to template switching, $r_{ts}(i,k)$ (Supplemental Note 3.1), the probability that a neuron in cubelet $i$ falsely projected to cubelet $k$ due to re-used barcodes, $r_{re}(i,k)$ (Supplemental Note 3.2), and the probability that a neuron in cubelet $i$ falsely projected to cubelet $k$ due to baseline contaminations, $r_{ba}(i,k)$. Note that $r_{ts}(i,k)$, $r_{re}(i,k)$, and $r_{ba}(i,k)$ were all very small, so the overall false-positive probability could be calculated additively. If there were $N(i)$ neurons in cubelet $i$, and $N_{pro}(i,k)$ neurons in cubelet $i$ were found to project to cubelet $k$, then the p value of $i$-to-$k$ connection, $v_{ik}$ was calculated with:

$$v_{ik} = 1 - f\left(N_{pro}(i,k), N_i, r_{ts}(i,k) + r_{re}(i,k) + r_{ba}(i,k)\right) \quad (13)$$

, where $f$ was the binomial cumulative distribution function:

$$f(n,N,p) = \sum_{l=0}^{n} \binom{N}{l} p^l (1-p)^{N-l} \quad (14)$$

.

With p-values, we were able to determine whether a given cubelet-to-cubelet connection was significant. Volcano plots of ipsilateral connections and contralateral connections in BL6-1 are shown in Figure S3G. All the analyses in the manuscript only included significant projections after multiple comparison correction (Bonferroni correction for multiple comparison, p-value < $\frac{0.05}{N}$, where N is the total number of possible projections) unless otherwise stated.

**Summary of error sources**

| Error sources | Effects | Solutions |
|---|---|---|
| Barcode base substitution | Generate barcodes with 1 or very few counts in 1 or very few cubelets | Collapse barcodes with up to 3 mismatches.<br>Set UMI threshold.<br>Set soma threshold. |
| Barcode base insertion/deletion | Generate barcodes with 1 or very few counts in 1 or very few cubelets | Set UMI threshold.<br>Set soma threshold. |
| CSI sequencing errors | Generate barcodes in 'non-existing' cubelets | CSIs that did not match any of the 288 used CSIs were excluded for further analysis |
| UMI sequencing errors | Cause overestimated barcode counts | Not corrected (But errors should be rare and uniformly randomly distributed) |
| Template switching | False projections | PCR with a large volume.<br>Set UMI threshold.<br>Calculate false-positive rates. |
| Re-used barcodes | False projections | Use a high diversity barcode library.<br>Exclude over-represented barcodes in the barcode library.<br>Set axon/soma threshold.<br>Calculate false-positive rates |
| Non-collected soma | Strongest projections were detected as somata | Set soma threshold. |

### Supplemental Note 4: Comparing BRICseq projectome with Allen Connectivity Atlas, and comparing between BRICseq mapped brains

#### List of variables in Supplemental Notes 4

| | |
|---|---|
| $A_k$ | Brain area-to-brain area connection matrix, type $k$ ($k$=1,2,3,4) |
| $C_k$ | Cubelet-to-cubelet connection matrix, type $k$ ($k$=1,2,3,4) |
| $P_k$ | Cubelet-to-brain area connection matrix, type $k$ ($k$=1,2,3,4) |
| $M$ | Cubelet-to-brain area mapping matrix |
| $M_a$ | Cubelet-to-brain area mapping matrix, normalized to total size of each brain area |
| $M_c$ | Cubelet-to-brain area mapping matrix, normalized to total size of each cubelet |
| $S_a$ | Brain area size matrix, diagonal |
| $S_c$ | Cubelet size matrix, diagonal |
| ρ | Neuron density, number of neurons per unit area size |

With BRICseq, we were able to map cubelet-to-cubelet connections from one individual brain. In order to compare between BRICseq data and Allen data, we utilized brain registration results to infer cubelet-to-brain area connections and brain area-to-brain area connections from cubelet-to-cubelet connections. Here we describe and discuss 2 models underlying connection inference: weighted averaging (Supplemental Note 4.1) and constrained optimization (Supplemental Note 4.2).

The following terms and variables are defined before further discussion:
Considering the connection from cubelet $i$ to cubelet $j$, $\{C\}_{ij}$, we could quantify its strength by calculating the average counts of UMIs (molecules) in cubelet $j$ per neuron in cubelet $i$ (See Supplemental Note 3.3). This described the projection strength (axon volume) from an average neuron in cubelet $i$ to the whole cubelet $j$, and thus was called 'unit-to-total' connection here. By considering the physical sizes of cubelet $i$ and cubelet $j$, we could also define and calculate 'unit-to-unit' connection (connection from a neuron in cubelet $i$ to a unit area size in cubelet $j$), 'total-to-unit' connection (connection from the whole cubelet $i$ to a unit area size in cubelet $j$), and 'total-to-total' connection (connection from the whole cubelet $i$ to the whole cubelet $j$), as summarized in the table below (similar to Supplemental Figure 2 in (Oh et al., 2014).

| Connection type | Connection source | Connection target | Definition | Formula |
|---|---|---|---|---|
| Type 1, $C_1$ | Cubelet | Cubelet | Unit neuron-to-unit area size | $C_1$ |

| | | | | |
|---|---|---|---|---|
| Type 2, $C_2$ | Cubelet | Cubelet | Unit neuron-to-total | $C_2 = C_1 S_c$ |
| Type 3, $C_3$ | Cubelet | Cubelet | Total-to-unit area size | $C_3 = \rho S_c C_1$ |
| Type 4, $C_4$ | Cubelet | Cubelet | Total-to-total | $C_4 = \rho S_c C_1 S_c$ |

Here $S_c$ is a diagonal matrix, whose element $\{S_c\}_{ii}$ represents the physical size of cubelet $i$, and ρ represents the number of neurons per unit area size, or neuron density. We assume that ρ is uniform, so the average connection strength from a unit area size in a source cubelet to a target is ρ times the average connection strength from a neuron in the source cubelet to the target.

In conventional fluorescence tracing, projection strength is usually quantified as the normalized fluorescence intensity in the target area to the fluorescence intensity in the injection area (Oh et al., 2014). This was analogous to the type 2 connection, as defined above. Connections mentioned in this manuscript all referred to type 2 connections, unless otherwise stated.

Similar to cubelet-to-cubelet connections, $C_k$ ($k$=1,2,3,4), we also defined 4 types of brain area-to-brain area connections, $A_k$ ($k$=1,2,3,4), and cubelet-to-brain area connections, $P_k$ ($k$=1,2,3,4), as summarized below.

| Connection type | Connection source | Connection target | Definition | Formula |
|---|---|---|---|---|
| Type 1, $A_1$ | Brain area | Brain area | Unit neuron-to-unit area size | $A_1$ |
| Type 2, $A_2$ | Brain area | Brain area | Unit neuron-to-total | $A_2 = A_1 S_a$ |
| Type 3, $A_3$ | Brain area | Brain area | Total-to-unit area size | $A_3 = \rho S_a A_1$ |
| Type 4, $A_4$ | Brain area | Brain area | Total-to-total | $A_4 = \rho S_c A_1 S_a$ |

| Connection type | Connection source | Connection target | Definition | Formula |
|---|---|---|---|---|
| Type 1, $P_1$ | Cubelet | Brain area | Unit neuron-to-unit area size | $P_1$ |
| Type 2, $P_2$ | Cubelet | Brain area | Unit neuron-to-total | $P_2 = P_1 S_a$ |
| Type 3, $P_3$ | Cubelet | Brain area | Total-to-unit area size | $P_3 = \rho S_c P_1$ |
| Type 4, $P_4$ | Cubelet | Brain area | Total-to-total | $P_4 = \rho S_c P_1 S_a$ |

Here $S_a$ is a diagonal matrix, and its element $\{S_a\}_{ii}$ represents the physical size of brain area $i$.

We also calculated a cubelet-to-brain area mapping matrix, $M$, based on cubelet registration results. $\{M\}_{ij}$ represents the physical size of the intersection of cubelet $i$ and brain area $j$. The mapping matrix $M$ was also normalized to either the total size of each brain area or to the total size of each cubelet:

$$M_a = M S_a^{-1} \quad (15)$$

$$M_c = S_c^{-1} M \quad (16)$$

. In $M_a$, the sum of each column is 1; in $M_c$, the sum of each row is 1.

**4.1 Inferring cubelet-to-brain area connections/brain area-to-brain area connections by weighted averaging** (Figure 2C-E; Figure S4B-D; Figure S11H).

While we have dissected the cortex into ~ 230 cubelets, there are ~ 70 brain cortical areas according to Allen atlas (2011 version). The size of a cortical area was much larger than a cubelet, and an area on average consisted of 10 cubelets. Thus we considered the cubelets as building blocks of brain connectivity and assumed connections between brain areas were weighted averages of cubelets contained (Figure S4B,D). With such an assumption, we had:

$$P_2 = C_1 M \quad (17)$$

$$A_3 = \rho M^T P_1 \quad (18)$$

, where $M^T$ denotes the transpose of $M$.

With Eq. (16) and (17), we got

$$P_2 = C_1 M = C_2 S_c^{-1} M = C_2 M_c \quad (19)$$

. With Eq. (15) and (18), we got

$$A_2 = \rho^{-1} S_a^{-1} A_3 \, S_a = \rho^{-1} S_a^{-1} \rho M^T P_1 S_a = (M S_a^{-1})^T \, P_1 S_a = M_a^T P_2 \quad (20)$$

. With Eq. (19) and (20), we got

$$A_2 = M_a^T P_2 = M_a^T C_2 M_c \quad (21)$$

. We inferred cubelet-to-brain area connections with (19) in Figure 2D,E; and inferred brain area-to-brain area connections with Eq. (21) in Figure 2C, Figure S11H.

To reduce the variations brought by dissection and registration errors, we downsampled the cubelet-to-cubelet connection matrix for analyses here. If $\alpha_0$ and $\beta_0$ were two cubelets, $\alpha_1, \alpha_2 \ldots \alpha_m$ were neighbors of $\alpha_0$, and $\beta_1, \beta_2 \ldots \beta_n$ were neighbors of $\beta_0$, then the projection strength from $\alpha_0$ to $\beta_0$, $C_{\alpha_0 - \beta_0}$ was downsampled as:

$$C_{\alpha_0-\beta_0} = \begin{pmatrix} 0.9 & \frac{0.1}{m} & \cdots & \frac{0.1}{m} \end{pmatrix} \begin{pmatrix} C_{\alpha_0-\beta_0} & C_{\alpha_0-\beta_1} & \cdots & C_{\alpha_0-\beta_n} \\ C_{\alpha_1-\beta_0} & C_{\alpha_1-\beta_1} & \cdots & C_{\alpha_1-\beta_n} \\ \vdots & \vdots & \ddots & \vdots \\ C_{\alpha_m-\beta_0} & C_{\alpha_m-\beta_1} & \cdots & C_{\alpha_m-\beta_n} \end{pmatrix} \begin{pmatrix} 0.9 \\ \frac{0.1}{n} \\ \vdots \\ \frac{0.1}{n} \end{pmatrix} \quad (22).$$

For the analysis in 4.1, all the non-significant cubelet-to-cubelet connections were set to 0. As multiple comparison had a high false negative rate particularly for weak projections, p value = 0.05 (no multiple comparison) was used for the criterion of significance here. For comparison between cubelets and injections in the same source brain area (Figure 2D,E), we require the cubelets reside primarily (>70%) in the brain area. When calculating brain area-to-brain area connections, only well-infected brain areas are included as source areas. A well-infected brain area is defined as an area where > 50% of the area's size is covered by cubelets infected with >5 neurons.

**4.2 Inferring brain area-to-brain area connections by constrained optimization** (Figure S4E-G).

In contrast to assuming cubelets, which were smaller in size, were building blocks of brain connections, connections of brain areas could also be inferred assuming input and output per unit area size win each brain area were homogeneous (Figure S4E) (Oh et al., 2014). With this assumption, we had:

$$P_3 = \rho M A_1 \quad (23)$$

$$C_2 = P_1 M^T \quad (24)$$

. The Eq. (23) and (24) corresponded to output homogeneity and input homogeneity, respectively.

With Eq. (16) and (23), we got

$$P_2 = \rho^{-1} S_c^{-1} P_3 S_a = \rho^{-1} S_c^{-1} \rho M A_1 S_a = M_c A_2 \quad (25)$$

. With Eq. (15) and (25), we got

$$C_2 = P_1 M^T = P_2 S_a^{-1} M^T = P_2 M_a^T \quad (26)$$

. With Eq. (25) and (26), we got

$$C_2 = M_c A_2 M_a^T \quad (27)$$

.

According to Eq. (27), we could estimate $A_2$ (least-squares solution) with:

$$\widetilde{A_2} = M_c^+ C_2 (M_a^+)^T \quad (28)$$

, where $\widetilde{A_2}$ is estimated $A_2$, and $M_c^+$ ($M_a^+$) is the pseudo-inverse matrix of $M_c$ ($M_a$). However, this might result in negative connection values. Thus, we determined to estimate $A_2$ with constrained optimization:

$$\widetilde{A_2} = argmin_{A_2}(\|C_2 - M_c A_2 M_a^T\|) \quad (29)$$

, with the constraint

$$A_2 \geq 0 \quad (30)$$

. With Eq. (29) and formula (30), we inferred brain area-to-brain area connections in Figure S4F,G.

To reduce the variations brought by registration errors, downsampling was also performed here for the cubelet-to-cubelet connection matrix with Eq. (22).

For the analysis in 4.2, all the non-significant cubelet-to-cubelet connections were set to 0. As multiple comparison had a high false negative rate particularly for weak projections, p value = 0.05 (no multiple comparison) was used for the criterion of significance here. Only well-infected brain areas are included as source areas. A well-infected brain area is defined as an area where > 50% of the area's size is covered by cubelets infected with >5 neurons.

**4.3 Discussions**

It remains a challenge to infer the underlying connections between brain areas from neural tracing experiments (Knox et al., 2018). In the real scenario, neither cubelets nor brain areas were necessarily homogeneous, and thus we did not aim to develop a method to precisely quantify brain area-to-brain area connection patterns here. However, by inferring brain area connections with abovementioned assumptions, we argue that it provided a fair approach to validate BRICseq and screen for long-range connection disruptions in neuropsychiatric disorders.

In the 'weighted averaging' approach, we assumed that individual cubelets were homogeneous, and broke down brain areas into cubelets contained. Ideally this would be correct if the size of cubelets was small enough so that each cubelet was a homogeneous unit. However, with the current experiment protocol, inferring connections with this method was still an estimation. In the 'constrained optimization' approach, the assumption that input and output of all the regions within a brain area were homogeneous was also imprecise. For example, the connections between primary visual cortex and higher visual areas are organized according to retinotopic maps (Wang and Burkhalter, 2007). Moreover, our results also indicated the rank of $C$ was much higher than the total number of brain areas (data not shown), arguing against the brain area homogeneity assumption (according to Eq. (27), the rank of $C$ should not be greater than the rank of $A$). Future work should be done to better define, quantify and calculate brain connections.

### Supplemental Note 5: Bioinformatics, statistics and computational methods

#### 5.1 Processing of raw sequencing data.

Raw Illumina sequencing results consisted of two .fastq files: 32-nt BC sequences were in paired end 1, and 12-nt UMI and 8-nt CSI sequences were in paired end 2. The full BC-UMI-CSI sequences were merged and then de-multiplexed based on CSIs (cubelets). All the sequences with ambiguous bases (shown as N in the sequencing results) were removed. We then collapsed all the identical reads. As the current sequencing depth was too low and most of the sequences only had 1 read each, we didn't set any threshold for read counts to remove errors (but see Supplemental Note 3). Unique sequences were next sorted into barcode library 1 (BC ended with 2 purines), barcode library 2 (BC ended with 2 pyrimidines), and spike-in (BC ended with ATCAGTCA). We then counted the number of unique UMIs for each BC-CSI, which represented the molecule count of a given barcode in a given cubelet.

#### 5.2 Substitution error correction.

Base substitution is one of the major error sources. As the theoretical diversity of a random barcode of $N_{30}YY$ or $N_{30}RR$ is $4^{30} \times 2^2 \approx 10^{18}$, an error barcode due to substitution should be very similar to one of the real barcodes, while any two real barcodes should be very different. To correct substitution errors, we first found all the barcode pairs with up to 3 mismatches using the short read aligner *bowtie* (http://bowtie-bio.sourceforge.net/index.shtml) (Langmead et al., 2009). We next collapsed all the barcodes into a large number of clusters, such that for any barcode (BC1) in a given cluster, there existed another barcode (BC2) in the same cluster with less than 3 mismatches. As a simple algorithm, mathematically it could cause very different barcodes to be collapsed into the same cluster; however, this did not happen in the real scenario due to the high theoretical diversity. The barcode with the highest UMI counts in each cluster was used to represent the cluster, and the summed UMI count of all the barcodes in the cluster was calculated as the corrected UMI count of the barcode. After substitution correction, we generated a barcode-cubelet matrix, where each element represented the molecule count of a given barcode in a given cubelet after collapsing.

#### 5.3 Reconstruction of single cell projections.

5.3.1 Viral abundance thresholding. To reduce re-used barcode errors, barcodes whose counts were greater than 4 in the viral library sequencing result were excluded for analysis in the barcode-cubelet matrix. See full details in Supplemental Note 5.5.

5.3.2 UMI thresholding. To remove noises, we set all the no-greater-than-1 (UMI threshold) elements in the matrix to 0.

5.3.3 Soma/axon thresholding. After barcode abundance thresholding and UMI thresholding, we determined the soma location of each barcode using the 'soma-max' strategy. To exclude local dendritic innervations, for each barcode, the UMI counts of all the cubelets neighboring to the soma cubelet were set to 0. Firstmax and secondmax were then calculated as the highest and second highest UMI counts for each barcode. We chose soma threshold to be 250 and axon threshold to be 20, and only analyzed barcodes whose firstmax was greater than soma threshold and secondmax was between UMI threshold and axon threshold. See full details in Supplemental Note 3.2.

5.3.4 Filter right cortical neurons. We remove the barcodes whose somas did not reside in the right cortical hemisphere. Cells not in the right cortex were extremely rare, and they were likely due to virus spread.

With these steps, we were able to determine each cell's location and its projection pattern.

**5.4 Calculating bulk projection patterns.**

To calculate bulk projection patterns, we pooled all the projection cells that resided in the same cubelets together, and calculated their average projection patterns. Projection strengths due to noise were evaluated and subtracted from the uncorrected projection strengths. P values were also calculated for each projection. See details in Supplemental Note 3.

In the manuscript, '(non-)significant connections (no multiple comparison)' refer to connections with p value (≥) < 0.05; '(non-)significant connections (multiple comparison)' refer to connections with p value (≥) < 0.05/N, where N is total number of possible connection (the number of right cortical cubelets times the number of all the cortical and subcortical cubelets).

Some of the RT primers were found to be cross-contaminated at low levels *post hoc*. Thus, we didn't analyze the projections between these contaminated cubelets. These projections include: BL6-1, cubelet 97-to-cubelet 68, cubelet 115-to-cubelet 130, cubelet 21-to-cubelet 268; BL6-2, cubelet 75-to-cubelet 13, cubelet 13-to-cubelet 75; BTBR-1, cubelet 60-to-cubelet 81, cubelet 81-to-cubelet 60.

**5.5. Simulation of re-used barcodes**

To set thresholds to reduce re-used barcodes and quantify false positive errors caused by re-used barcodes, we need to simulate re-used barcodes and calculate their distribution in the (firstmax, secondmax) space (details in Supplemental Note 3.2.2).

Step 1. Exclude overrepresented barcodes in the barcode library.

The distribution of barcode abundance in the barcode library was not uniform, so barcodes with higher abundance in the library were more likely to be re-used in multiple neurons. Moreover, as we did not sequence the full viral barcode library, we also found barcodes present in the BRICseq result but absent in the viral library sequencing result. We set a viral barcode threshold (=4), and classified barcodes according to their abundance: overrepresented barcodes (present and over 4 counts in the library sequencing result), underrepresented barcodes (present but no-greater-than 4 counts in the library sequencing result), and non-sequenced barcodes (absent in the library sequencing result, but present in the BRICseq result). To reduce the chance of re-used barcodes, we included underrepresented barcodes and non-sequenced barcodes for neuronal projection analysis. But for re-used barcodes simulation as follows, we only included underrepresented barcodes; because non-sequenced barcodes have lower copies in the viral library than underrepresented barcodes, the re-used barcode chance in the non-sequenced barcodes should be lower than the chance in the underrepresented barcodes.

Step 2. Simulate re-used barcodes and calculate their distributions in the (firstmax, secondmax) space.
To simulate re-used barcodes, we assumed 1) most of the observed barcodes were single-cell barcodes (re-used barcodes were rare) and 2) the barcode expression level and projection patterns of an infected cell were independent of the barcode sequence itself. The simulation steps were:
1. Estimate total number of re-used barcodes. We calculated the total amount of underrepresented barcodes in the MAPseq results (including type 1-4 barcodes) $N_t$, and sampled $N_t$ barcodes from underrepresented barcodes in the barcode library sequencing result using the observed abundance distribution. We estimated the total number of re-used barcodes, $N_{re}$ from the sampling simulation.
2. Estimate the distribution of re-used barcodes in the (firstmax, secondmax) space. As most re-used barcodes were used twice, we randomly sampled firstmax and secondmax from the measured distributions for two neurons related to each re-used barcode, and calculated the new firstmax and secondmax for this barcode. By doing this $N_{re}$ times, we generated a distribution of re-used barcodes in the (firstmax, secondmax) space.
3. Given the simulated distribution of re-used barcodes, we calculated the ratio of re-used barcodes to the total number of barcodes in the BRICseq result in the (firstmax, secondmax) space. (Figure S3F).

**5.6 Distribution of connection strengths and projection types**
To calculate distributions of connection strengths and distance-dependent connection properties (connection strength, connection probability, and input/output correlation), only significant non-zero (with Bonferroni multiple comparison correction) cubelet-to-cubelet connections were included. The distance between 2 cubelets was defined as the distance of their centroids.

The contralateral homotopic cubelet for a given cubelet was determined based on the symmetry of laser dissection. Considering sectioning/dissection variations, we also generalized contralateral homotopic cubelets to include all the neighbor cubelets of the exact contralateral homotopic cubelet. Generalized contralateral homotopic cubelets were used for all the analysis regarding contralateral projections.

**5.7 Analysis of modules.**
We utilized the Brain Connectivity Toolbox (https://sites.google.com/site/bctnet/) for module analysis in Matlab. *modularity_dir.m* was used to find modules in the connectivity matrix (directed graph), and *modularity_und.m* was used to find modules in the input/output correlation matrix (undirected graph). In input/output correlation matrix, negative values were set to 0 before clustering. A resolution parameter $\gamma$ can be tuned to get smaller/more or larger/fewer modules. To determine the optimal $\gamma$, we undersampled

half of the total projection neurons for 100 times, and performed clustering with various γ. For each γ, we calculated the average number of modules over 100 undersampling trials, and quantified the inconsistency of clustering that was defined as the mean of Rand indices between pairwise trials' clustering results. The optimal γ was chosen so that the inconsistency was low and the average number of modules was stable (Figure S13A). All the analyses were done with the optimal γ unless otherwise stated.
To generate the distance-dependent connection matrix, we first calculated connection strengths and physical distances for all cubelet pairs. We next grouped cubelet pairs into bins according to the distances (50 μm each bin), and calculated the mean connection strength in each bin. Then in the distance-dependent connection matrix, each element was set to the mean connection strength of the bin it belonged to. To calculate the distance-independent connection matrix, the distance-dependent connection matrix was subtracted from the original connection matrix. Negative values in the distance-independent connection matrix were set to 0 before clustering. The distance between 2 cubelets was defined as the distance of their centroids.

Clustering results were compared with Rand indices (Rand, 1971).

For module analysis, non-significant (with Bonferroni multiple comparison correction) cubelet-to-cubelet projections were set to 0.

**5.8 Analysis of motifs.**
*clustering_coef_bd.m* in the Brain Connectivity Toolbox was used to calculate the clustering coefficient. The connection matrix was binarized for this analysis. For comparison, we generated random connection networks based on distance-dependent connection probability rule: in the real network, we calculated the probability that cubelet $i$ projected to cubelet $j$ if their distance was $d$ (in 50 μm bins); then the measured probabilities were used to generate 10000 random networks assuming each connection was independent.

Three types of 2-node motifs and 16 types of 3-node motifs were counted in real cortical networks. Random networks were also simulated to calculate the relative abundance of each motif in real networks. The relative abundance was calculated with:

$$\frac{Count_{real}(motif\ i) - Count_{random}(motif\ i)}{Count_{random}(motif\ i)}.$$

Different models were used to generate random networks, and 10000 random networks were generated each:
In 2-node motif comparison, $RN_g$ was generated based on a global connection probability rule: in the real network, we calculated the probability that cubelet $i$ projected to cubelet $j$; then the measured probability was used to generate $RN_g$ assuming each connection was independent.
In 2-node motif comparison, $RN_{dd}$ was generated based on a distance-dependent connection probability rule: in the real network, we calculated the probability that cubelet $i$ projected to cubelet $j$ if their distance was $d$ (in 50 μm bins); then the measure probabilities were used to generate $RN_{dd}$ assuming each connection was independent.

In 3-node motif comparison, $RN_g$ was generated based on a global 2-node motif probability rule: in the real network, we calculated the probability of each 2-node motif between cubelet $i$ and cubelet $j$, then the measured probability was used to generate $RN_g$ assuming each 2-node motif was independent.
In 3-node motif comparison, $RN_{dd}$ was generated based on a distance-dependent 2-node motif probability rule: in the real network, we calculated the probability of each 2-node motif between cubelet $i$ and cubelet $j$ if their distance was $d$ (in 50 μm bins), then the measured probabilities was used to generate $RN_{dd}$ assuming each 2-node motif was independent.

For all the analysis in 5.7, the distance between 2 cubelets was defined as the distance of their centroids.

For motif analysis, non-significant (with Bonferroni multiple comparison correction) cubelet-to-cubelet projections were set to 0.

**5.9 Analysis of function-connection relationship**

With the functional imaging data, we first performed singular-value decomposition with the activity matrix (pixel-by-time). The first 500 components were included for following analysis. To determine the activity of each cubelet, we calculated the mean activity over all pixels belong to the same cubelet. The activity correlation was calculated using activity data in all the time frames of all the trials. The spontaneous correlation was calculated using activity data from 0-1s of all the trials (note the initialization of each trial was at 2±0.2 s). To calculate the noise correlation, we grouped them into auditory-left-correct (modality-choice-result), auditory-right-correct, visual-left-correct, visual-right-correct, auditory-left-incorrect, auditory-right-incorrect, visual-left-incorrect, visual-right-incorrect trial groups. The mean activity at a given time point over all the trials in the same group was subtracted from the original activity data belonging to the corresponding trial group to calculate noises. All the correlations were calculated as Pearson correlations.

The down-sampled connection matrices (Supplemental Note 4.1) were used for the connection analysis. The reciprocal connection strength was calculated as the mean of logarithm of connection strengths in two directions. To compare function data with connection data, we only included cubelet pairs that satisfied 1) number of infected cells in both cubelets were greater than 50 in BRICseq, 2) both cubelets were well imaged (excluding non-surface areas like orbitofrontal cortex/anterior cingulate cortex/retrosplenial cortex, and lateral areas like insular cortex), 3) the two cubelets in a pair were not neighbors (neighbor connections were not analyzed in BRICseq).

To remove distance-dependent components from activity correlations, spontaneous correlations, noise correlations, connection strengths, and input correlations, we grouped cubelets pairs into bins according to the distances (50 μm each bin), and calculated the mean value of each variable in each bin. The mean value of each variable was then subtracted from the original data in the corresponding bins to calculate distance-independent components. The averaging and subtraction of connection strengths were performed in the logarithm form. The distance between 2 cubelets was defined as the distance of their centroids.

For the analysis in 5.9, all the non-significant cubelet-to-cubelet connections were set to 0. P value = 0.05 (no multiple comparison) was used for the criterion of significance here.

**5.10 Analysis of gene expression-connection relationship**

5.10.1 Pre-processing of the *in situ* hybridization data

The Allen *in situ* hybridization data were downloaded and registered to the coordinates of BRICseq cubelets in BL6-1 and BL6-2. The expression of gene X in cubelet Y was calculated as the average expression of gene X in all the voxels located in cubelet Y. The expression was quantified as the sum of intensity of expressing pixels divided by the total number of pixels (defined as energy in Allen *in situ* hybridization database). Only *in situ* hybridization data from coronal sections were used because typically expression data in lateral brain areas are missing in sagittal sections. To select genes with high quality expression data for later analysis, we calculated the correlation coefficients of the expression levels of the same genes between data from saggital and coronal sections across the shared cubelets, and only included genes with Pearson r > 0.8. The selected genes also had higher expression levels and dispersion metrics (variance divided by mean) than the rest (data not shown), suggesting that these genes were with high signal-to-noise ratios and high variance. The pre-processing of gene expression data resulted in gene expression matrices $\boldsymbol{G}$ where each row represented a cubelet, and each column represented a filtered gene for BL6-1 and BL6-2.

5.10.2 Principal component analysis (PCA) of the connectivity data

To identify features that explained most of the connectivity data and were invariant between two brains (BL6-1 and BL6-2), we first calculated cubelet-to-brain area connectivity matrix $\boldsymbol{C}$ based on BRICseq data of BL6-1 and BL6-2 (Supplemental Note 4.1; each source cubelet was considered as an observation represented in each row, and the projection strength to each target brain area was considered as a feature represented in each column), and performed principal component analysis (PCA) on $\boldsymbol{C}_1$ (in what follows, the subscript 1 denotes BL6-1 and the subscript 2 denotes BL6-2). The eigenvector matrix $\boldsymbol{W}_1$ consisted of eigenvectors of $\boldsymbol{C}_1^T\boldsymbol{C}_1$, and the loading matrix $\boldsymbol{P}_1$ was determined with $\boldsymbol{P}_1 = \boldsymbol{C}_1\boldsymbol{W}_1$(Figure S7B,C). Next, we reconstructed cubelet-to-brain area connectivity $\widetilde{\boldsymbol{C}_1}$ using a subset of top PCs $\widetilde{\boldsymbol{P}}_1$ with $\widetilde{\boldsymbol{C}_1} = \widetilde{\boldsymbol{P}}_1\boldsymbol{W}_1^{-1}$, where $\boldsymbol{W}_1^{-1}$ denotes the inverse of $\boldsymbol{W}_1$. To quantify how the subset of PCs explained the full data in BL6-1, we calculated the Pearson r between $\widetilde{\boldsymbol{C}_1}$ and $\boldsymbol{C}_1$. To quantify how the subset of PCs explained the shared connectivity patterns between BL6-1 and BL6-2, we first did coordinate transformation to predict cubelet-to-brain area connectivity of BL6-1 cubelets, $\boldsymbol{C}_1^*$, using cubelet-to-brain area connectivity data in BL6-2, $\boldsymbol{C}_2$, assuming cubelets in BL6-2 are homogeneous (similar to Supplemental Note 4). Then the Pearson r between reconstructed connectivity data in BL6-1, $\widetilde{\boldsymbol{C}_1}$ and the BL6-2-predicted connectivity data of BL6-1, $\boldsymbol{C}_1^*$ was calculated to quantify the shared connectivity patterns between reconstructed BL6-1 and BL6-2. We found that top 10 PCs were able to explain a large fraction of the data in BL6-1 as well as shared data between BL6-1 and BL6-2 (Figure 4A). Thus, in the following analysis, top 10 PC loadings were used to represent projection patterns for all the cubelets in BL6-1 and BL6-2: $\boldsymbol{P}_1 = \boldsymbol{C}_1\boldsymbol{W}_1$, $\boldsymbol{P}_2 = \boldsymbol{C}_2\boldsymbol{W}_1$, and $\widetilde{\boldsymbol{P}_1}$ and $\widetilde{\boldsymbol{P}_2}$ are top 10 dimensions of $\boldsymbol{P}_1$ and $\boldsymbol{P}_2$.

5.10.3 Feature selection and linear regression

A greedy feature selection algorithm was applied to find feature gene set $S$, which predicted the loadings of top 10 projection PCs. The feature selection started from an empty feature set $S = \emptyset$, and in each iteration, one more feature $g$ was selected and added to the feature set $S = S \cup \{g_i\}$, to minimize the mean squared error of a linear regression model that fit the PC loadings $\widetilde{\boldsymbol{P}}$ with the expression data of genes in the feature set $\boldsymbol{G}_{S \cup \{g_i\}}$:

$$g = argmin_{g_i}\left(min_{\boldsymbol{U},\boldsymbol{\Lambda}}\left\|\boldsymbol{G}_{S \cup \{g_i\}}\boldsymbol{U} + \boldsymbol{\Lambda} - \widetilde{\boldsymbol{P}}\right\|_2\right)$$

, where $\boldsymbol{G}_{S \cup \{g_i\}}$ denotes the expression of genes in the set $S \cup \{g_i\}$, $\boldsymbol{U}$ and $\boldsymbol{\Lambda}$ denote the coefficients and intercepts of the linear regression model, and $\|\boldsymbol{X}\|_2$ denotes the L2-norm of the matrix $\boldsymbol{X}$.

The feature selection process was stopped when 30 gene features were selected. To avoid overfitting, 5-fold cross-validation was performed for the linear regression model to calculate the mean squared error during feature selection. Both the training data and the testing data used for feature selection were from the mouse BL6-1. After feature selection, a linear regression model was used to fit the PC loadings $\widetilde{\boldsymbol{P}}$ with the expression of the selected feature genes $\boldsymbol{G}_S$ with a training set (80%) from BL6-1 (Figure 4D). To quantify the predictability of the linear model, the coefficients were used to predict the connectivity data in the testing set of BL6-1 as well as in all the data of BL6-2 (Figure 4B,C; Figure S7D,E). The reconstructed cubelet-to-brain area projection data $\widetilde{\boldsymbol{C}}$ was calculated as $\widetilde{\boldsymbol{C}} = max\left(0, \widetilde{\boldsymbol{P}}\boldsymbol{W}^{-1}\right)$, where $\boldsymbol{W}^{-1}$ is the inverse of $\boldsymbol{W}$.

5.10.4 Data shuffling and the null distribution

To determine the null performance of the feature selection and the linear prediction model, we shuffled the gene expression matrix $\boldsymbol{G}$ within each column (each gene) for BL6-1. Next, we used the same algorithm as in 5.10.3 to find a feature set $S^*$ that could predict connectivity $\widetilde{\boldsymbol{P}}$ with shuffled gene expression $\boldsymbol{G}^*$. Similarly, the feature selection was performed using data from BL6-1, with 5-fold cross-validation. After finding the gene predictors, we fit the connectivity data $\widetilde{\boldsymbol{P}}$ with the expression of the selected genes $\boldsymbol{G}_S^*$ using a training set (80%) from BL6-1, and quantified the predictability (Pearson r) of the linear model by using the fitting coefficients to predict the connectivity data in the testing set of BL6-1. The whole process was repeated for 100 times, to determine the 95% confidence interval of the null performance.

**Supplemental Table 1. Sindbis viral injections**
**Supplemental Table 2. Cubelet-to-cubelet projection strengths**
**Supplemental Table 3. Cubelet registration result**
**Supplemental Table 4. Inferred brain area-to-brain area connection strengths**
**Supplemental Table 5. Area names in Figure S4**

**Supplemental Movie 1. Summary of BRICseq results**

Bota, M., Sporns, O., and Swanson, L.W. (2015). Architecture of the cerebral cortical association connectome underlying cognition. Proc. Natl. Acad. Sci. *112*, E2093–E2101.

Chen, X., Zhan, H., Kebschull, J.M., Sun, Y., Zador, A.M., Zhan, H., Sun, Y., and Zador, A.M. (2018). Spatial organization of projection neurons in the mouse auditory cortex identified by in situ barcode sequencing. BioRxiv 294637.

Dechery, J.B., and MacLean, J.N. (2018). Functional triplet motifs underlie accurate predictions of single-trial responses in populations of tuned and untuned V1 neurons. PLoS Comput. Biol. *14*.

Han, Y., Kebschull, J.M., Campbell, R.A.A., Cowan, D., Imhof, F., Zador, A.M., and Mrsic-Flogel, T.D. (2018). The logic of single-cell projections from visual cortex. Nature.

Harris, J.A., Mihalas, S., Hirokawa, K.E., Whitesell, J.D., Knox, J., Bernard, A., Bohn, P., Caldejon, S., Casal, L., Cho, A., et al. (2018). The organization of intracortical connections by layer and cell class in the mouse brain. BioRxiv 292961.

Jaynes, E.T. (1957). Information theory and statistical mechanics. II. Phys. Rev. *108*, 171–190.

Kebschull, J.M., and Zador, A.M. (2015). Sources of PCR-induced distortions in high-throughput sequencing data sets. Nucleic Acids Res. *43*.

Kebschull, J.M., Garcia da Silva, P., Reid, A.P., Peikon, I.D., Albeanu, D.F., and Zador, A.M. (2016). High-Throughput Mapping of Single-Neuron Projections by Sequencing of Barcoded RNA. Neuron *91*, 975–987.

Knox, J.E., Harris, K.D., Graddis, N., Whitesell, J.D., Zeng, H., Harris, J.A., Shea-Brown, E., and Mihalas, S. (2018). High-resolution data-driven model of the mouse connectome. Netw. Neurosci. *3*, 217–236.

Langmead, B., Trapnell, C., Pop, M., and Salzberg, S.L. (2009). Ultrafast and memory-efficient alignment of short DNA sequences to the human genome. Genome Biol. *10*.

Musall, S., Kaufman, M.T., Gluf, S., and Churchland, A. (2018). Movement-related activity dominates cortex during sensory-guided decision making. BioRxiv 308288.

Oh, S.W., Harris, J.A., Ng, L., Winslow, B., Cain, N., Mihalas, S., Wang, Q., Lau, C., Kuan, L., Henry, A.M., et al. (2014). A mesoscale connectome of the mouse brain. Nature *508*, 207–214.

Paul, L.K. (2011). Developmental malformation of the corpus callosum: A review of typical callosal development and examples of developmental disorders with callosal involvement. J. Neurodev. Disord. *3*, 3–27.

Rand, W.M. (1971). Objective criteria for the evaluation of clustering methods. J. Am. Stat. Assoc. *66*, 846–850.

Rubinov, M., and Sporns, O. (2010). Complex network measures of brain connectivity: Uses and interpretations. Neuroimage *52*, 1059–1069.

Song, S., Sjöström, P.J., Reigl, M., Nelson, S., and Chklovskii, D.B. (2005). Highly nonrandom features of synaptic connectivity in local cortical circuits. In PLoS Biology, pp. 0507–0519.

Sperry, R.W., Gazzaniga, M.S., and Bogen, J.E. (1969). Interhemispheric relationships: the neocortical commissures; syndromes of hemisphere disconnection. Handb. Clin. Neurol. 273–290.

Vanni, M.P., and Murphy, T.H. (2014). Mesoscale Transcranial Spontaneous Activity Mapping in GCaMP3 Transgenic Mice Reveals Extensive Reciprocal Connections between Areas of Somatomotor Cortex. J. Neurosci. *34*, 15931–15946.

Wang, Q., and Burkhalter, A. (2007). Area map of mouse visual cortex. J. Comp. Neurol. *502*, 339–357.

Zingg, B., Hintiryan, H., Gou, L., Song, M.Y., Bay, M., Bienkowski, M.S., Foster, N.N., Yamashita, S., Bowman, I., Toga, A.W., et al. (2014). Neural networks of the mouse neocortex. Cell *156*, 1096–1111.
