## Supplemental Figures for "BRICseq bridges brain-wide interregional connectivity to neural activity and gene expression in single animals"

Figure S1

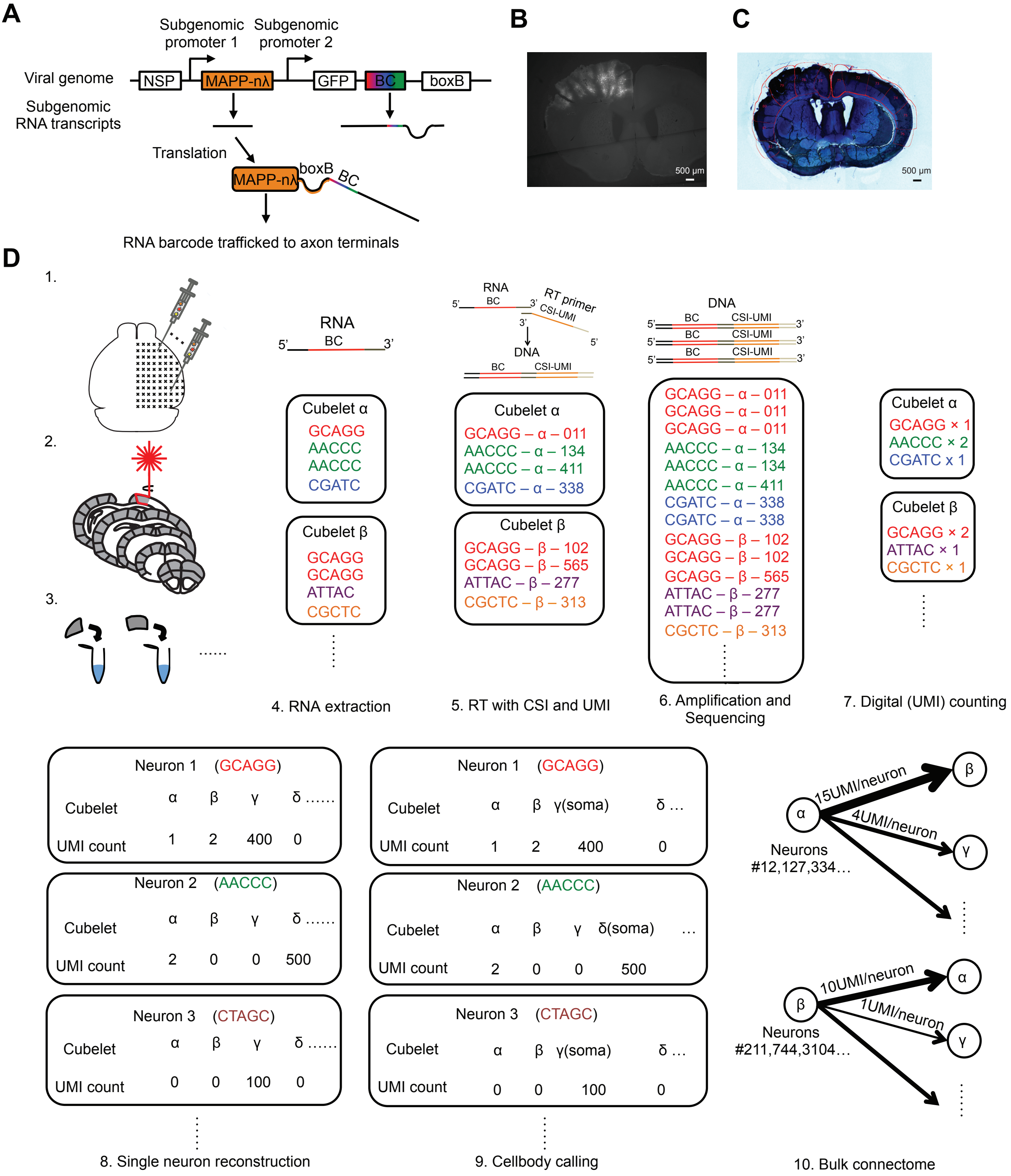

A
Subgenomic promoter 1
Subgenomic promoter 2
Viral genome
NSP
MAPP-nλ
GFP
BC
boxB
Subgenomic RNA transcripts
Translation
MAPP-nλ
boxB
BC
RNA barcode trafficked to axon terminals
B
500 μm
C
500 μm
D
1.
2.
3.
......
RNA
BC
5'
3'
Cubelet α
GCAGG
AACCC
AACCC
CGATC
Cubelet β
GCAGG
GCAGG
ATTAC
CGCTC
4. RNA extraction
RNA
BC
RT primer
CSI-UMI
DNA
BC
CSI-UMI
Cubelet α
GCAGG – α – 011
AACCC – α – 134
AACCC – α – 411
CGATC – α – 338
Cubelet β
GCAGG – β – 102
GCAGG – β – 565
ATTAC – β – 277
CGCTC – β – 313
5. RT with CSI and UMI
DNA
BC
CSI-UMI
GCAGG – α – 011
GCAGG – α – 011
GCAGG – α – 011
AACCC – α – 134
AACCC – α – 134
AACCC – α – 411
CGATC – α – 338
CGATC – α – 338
GCAGG – β – 102
GCAGG – β – 102
GCAGG – β – 565
ATTAC – β – 277
ATTAC – β – 277
CGCTC – β – 313
6. Amplification and Sequencing
Cubelet α
GCAGG × 1
AACCC × 2
CGATC x 1
Cubelet β
GCAGG × 2
ATTAC × 1
CGCTC × 1
7. Digital (UMI) counting
Neuron 1 (GCAGG)
Cubelet α β γ δ ......
UMI count 1 2 400 0
Neuron 2 (AACCC)
Cubelet α β γ δ ......
UMI count 2 0 0 500
Neuron 3 (CTAGC)
Cubelet α β γ δ ......
UMI count 0 0 100 0
8. Single neuron reconstruction
Neuron 1 (GCAGG)
Cubelet α β γ(soma) δ ...
UMI count 1 2 400 0
Neuron 2 (AACCC)
Cubelet α β γ δ(soma) ...
UMI count 2 0 0 500
Neuron 3 (CTAGC)
Cubelet α β γ(soma) δ ...
UMI count 0 0 100 0
9. Cellbody calling
α
β
γ
15UMI/neuron
4UMI/neuron
Neurons #12,127,334...
β
α
γ
10UMI/neuron
1UMI/neuron
Neurons #211,744,3104...
10. Bulk connectome

Figure S2

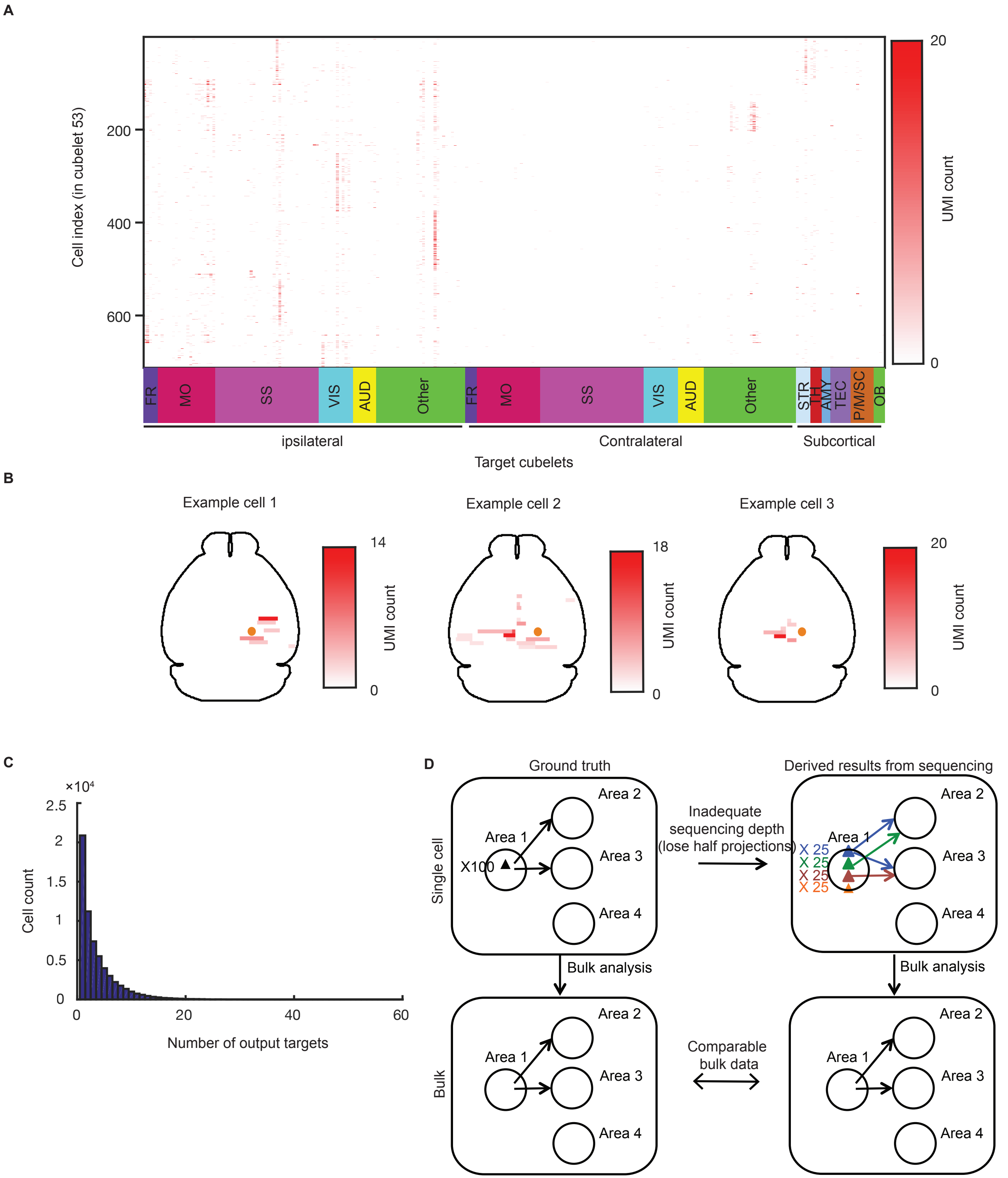
A
Cell index (in cubelet 53)
200
400
600
20
0
UMI count
FR
MO
SS
VIS
AUD
Other
FR
MO
SS
VIS
AUD
Other
STR
TH
AMY
TEC
P/M/SC
OB
ipsilateral
Contralateral
Subcortical
Target cubelets
B
Example cell 1
14
0
UMI count
Example cell 2
18
0
UMI count
Example cell 3
20
0
UMI count
C
×10⁴
2.5
2
1.5
1
0.5
0
Cell count
0
20
40
60
Number of output targets
D
Ground truth
Derived results from sequencing
Single cell
Area 1
Area 2
Area 3
Area 4
X100
Inadequate
sequencing depth
(lose half projections)
Area 1
Area 2
Area 3
Area 4
X 25
X 25
X 25
X 25
Bulk analysis
Bulk analysis
Bulk
Area 1
Area 2
Area 3
Area 4
Comparable
bulk data
Area 1
Area 2
Area 3
Area 4

Figure S3

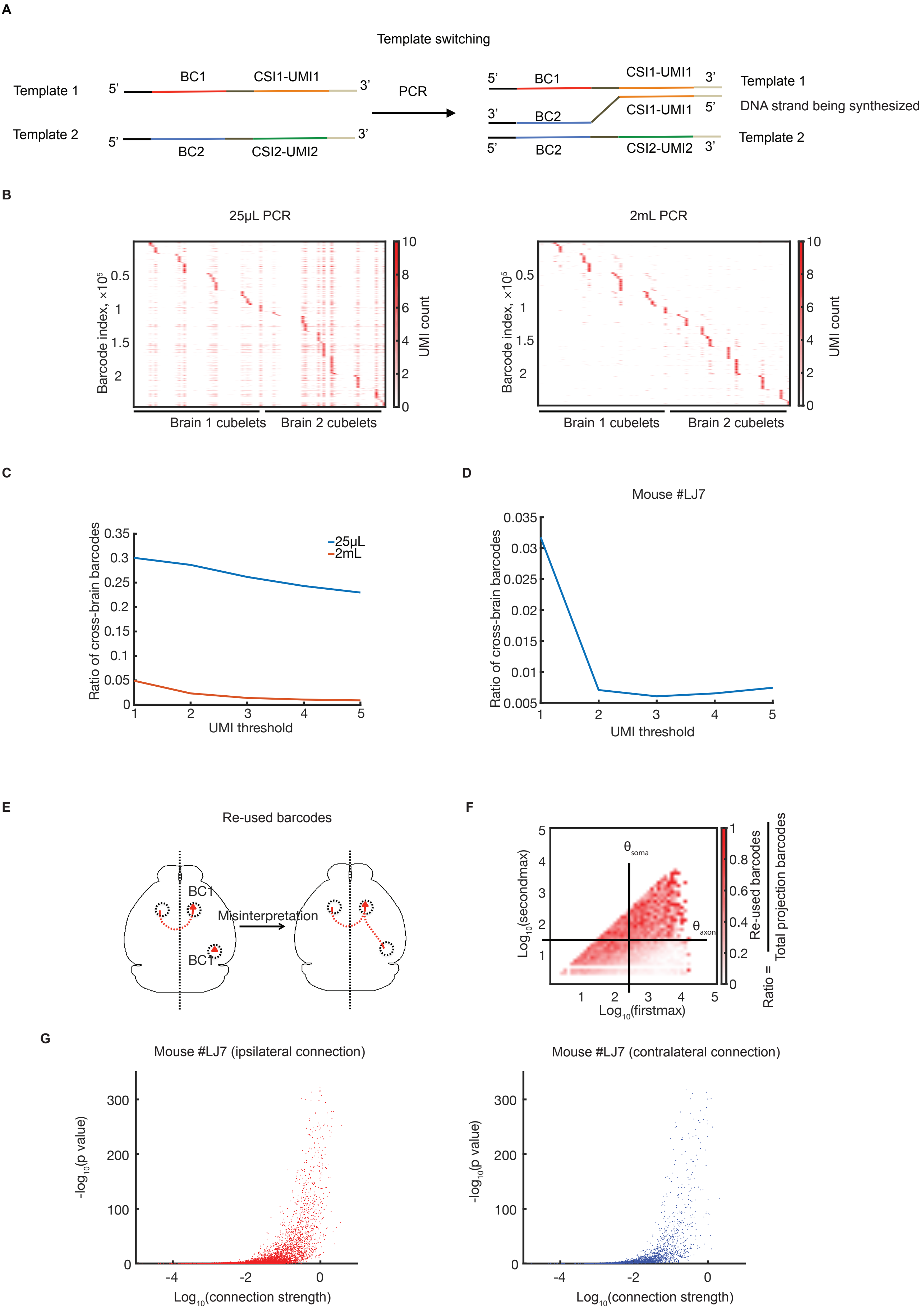
A
Template switching
Template 1
Template 2
5'
3'
BC1
CSI1-UMI1
BC2
CSI2-UMI2
PCR
DNA strand being synthesized
B
25μL PCR
2mL PCR
Barcode index, ×10^5
0.5
1
1.5
2
UMI count
10
8
6
4
2
0
Brain 1 cubelets
Brain 2 cubelets
C
Ratio of cross-brain barcodes
0.35
0.3
0.25
0.2
0.15
0.1
0.05
0
25μL
2mL
UMI threshold
1
2
3
4
5
D
Mouse #LJ7
0.035
0.03
0.025
0.02
0.015
0.01
0.005
E
Re-used barcodes
BC1
Misinterpretation
F
Log10(secondmax)
Log10(firstmax)
θsoma
θaxon
Ratio = Re-used barcodes / Total projection barcodes
1
0.8
0.6
0.4
0.2
0
G
Mouse #LJ7 (ipsilateral connection)
Mouse #LJ7 (contralateral connection)
-log10(p value)
300
200
100
0
Log10(connection strength)
-4
-2
0

Figure S4

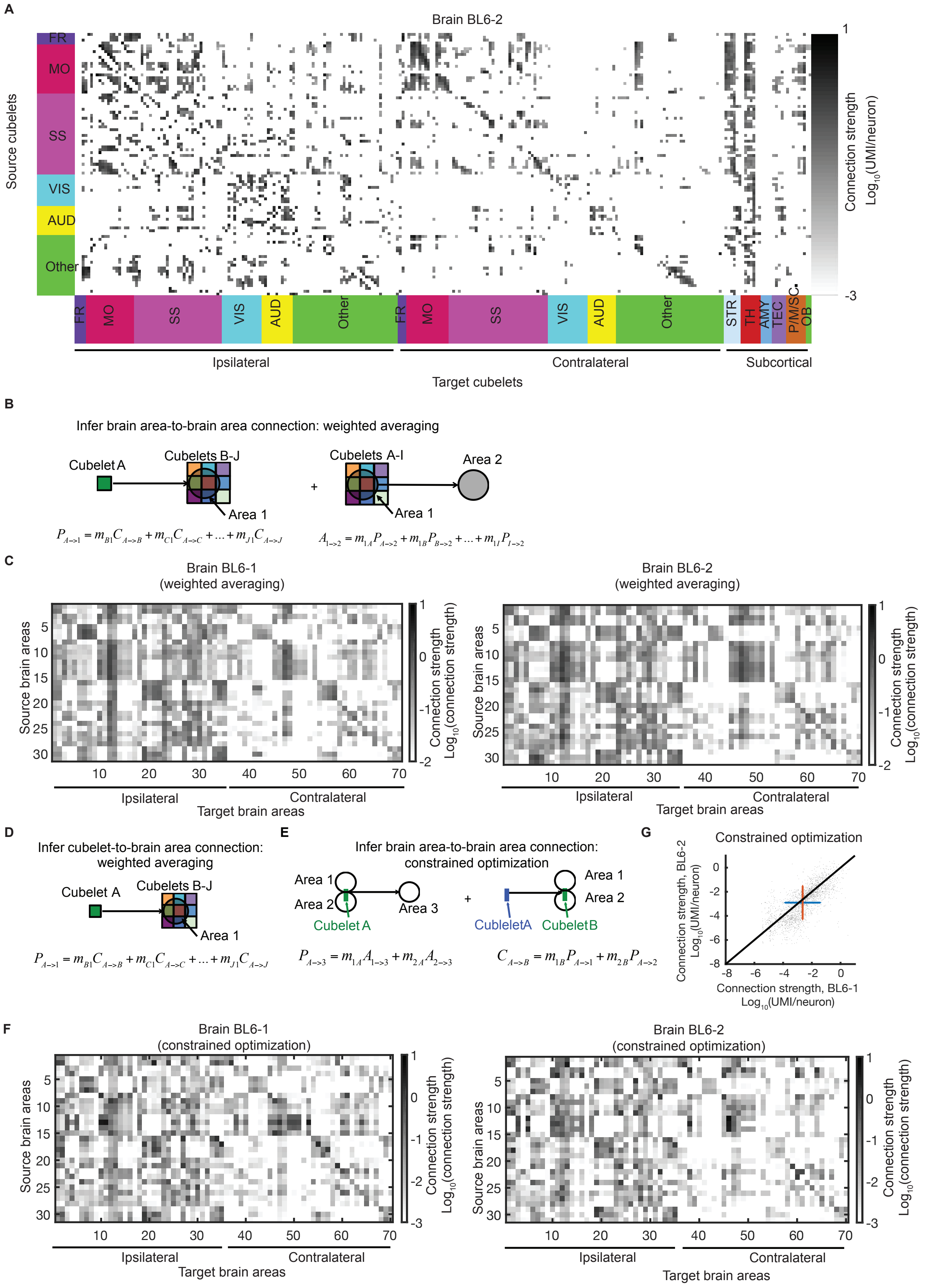

A
Brain BL6-2
Source cubelets
FR
MO
SS
VIS
AUD
Other
Connection strength
Log10(UMI/neuron)
1
-3
STR
TH
AMY
TEC
P/M/SC
OB
Ipsilateral
Contralateral
Subcortical
Target cubelets
B
Infer brain area-to-brain area connection: weighted averaging
Cubelet A
Cubelets B-J
Area 1
Cubelets A-I
Area 2
$P_{A\to1} = m_{B1}C_{A\to B} + m_{C1}C_{A\to C} + \ldots + m_{J1}C_{A\to J}$
$A_{1\to2} = m_{1A}P_{A\to2} + m_{1B}P_{B\to2} + \ldots + m_{1I}P_{I\to2}$
C
Brain BL6-1
(weighted averaging)
Brain BL6-2
(weighted averaging)
Source brain areas
Target brain areas
Ipsilateral
Contralateral
Connection strength
Log10(connection strength)
D
Infer cubelet-to-brain area connection: weighted averaging
Cubelet A
Cubelets B-J
Area 1
$P_{A\to1} = m_{B1}C_{A\to B} + m_{C1}C_{A\to C} + \ldots + m_{J1}C_{A\to J}$
E
Infer brain area-to-brain area connection: constrained optimization
Area 1
Area 2
Area 3
Cubelet A
CubleletA
CubeletB
$P_{A\to3} = m_{1A}A_{1\to3} + m_{2A}A_{2\to3}$
$C_{A\to B} = m_{1B}P_{A\to1} + m_{2B}P_{A\to2}$
G
Constrained optimization
Connection strength, BL6-2
Log10(UMI/neuron)
Connection strength, BL6-1
Log10(UMI/neuron)
F
Brain BL6-1
(constrained optimization)
Brain BL6-2
(constrained optimization)
Source brain areas
Target brain areas
Ipsilateral
Contralateral
Connection strength
Log10(connection strength)

Figure S5

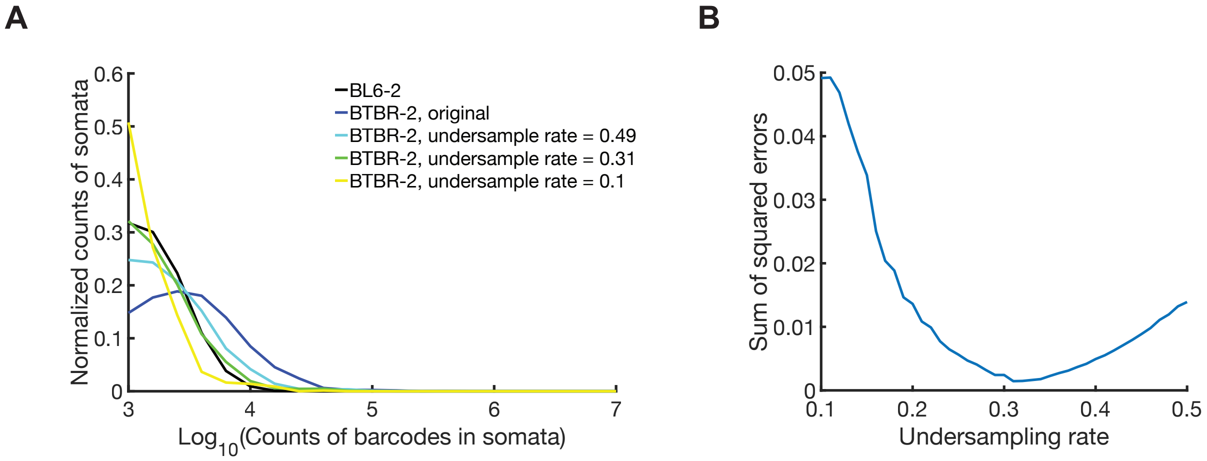
A
0.6
0.5
0.4
0.3
0.2
0.1
0
Normalized counts of somata
BL6-2
BTBR-2, original
BTBR-2, undersample rate = 0.49
BTBR-2, undersample rate = 0.31
BTBR-2, undersample rate = 0.1
3
4
5
6
7
Log10(Counts of barcodes in somata)
B
0.05
0.04
0.03
0.02
0.01
0
Sum of squared errors
0.1
0.2
0.3
0.4
0.5
Undersampling rate

Figure S6

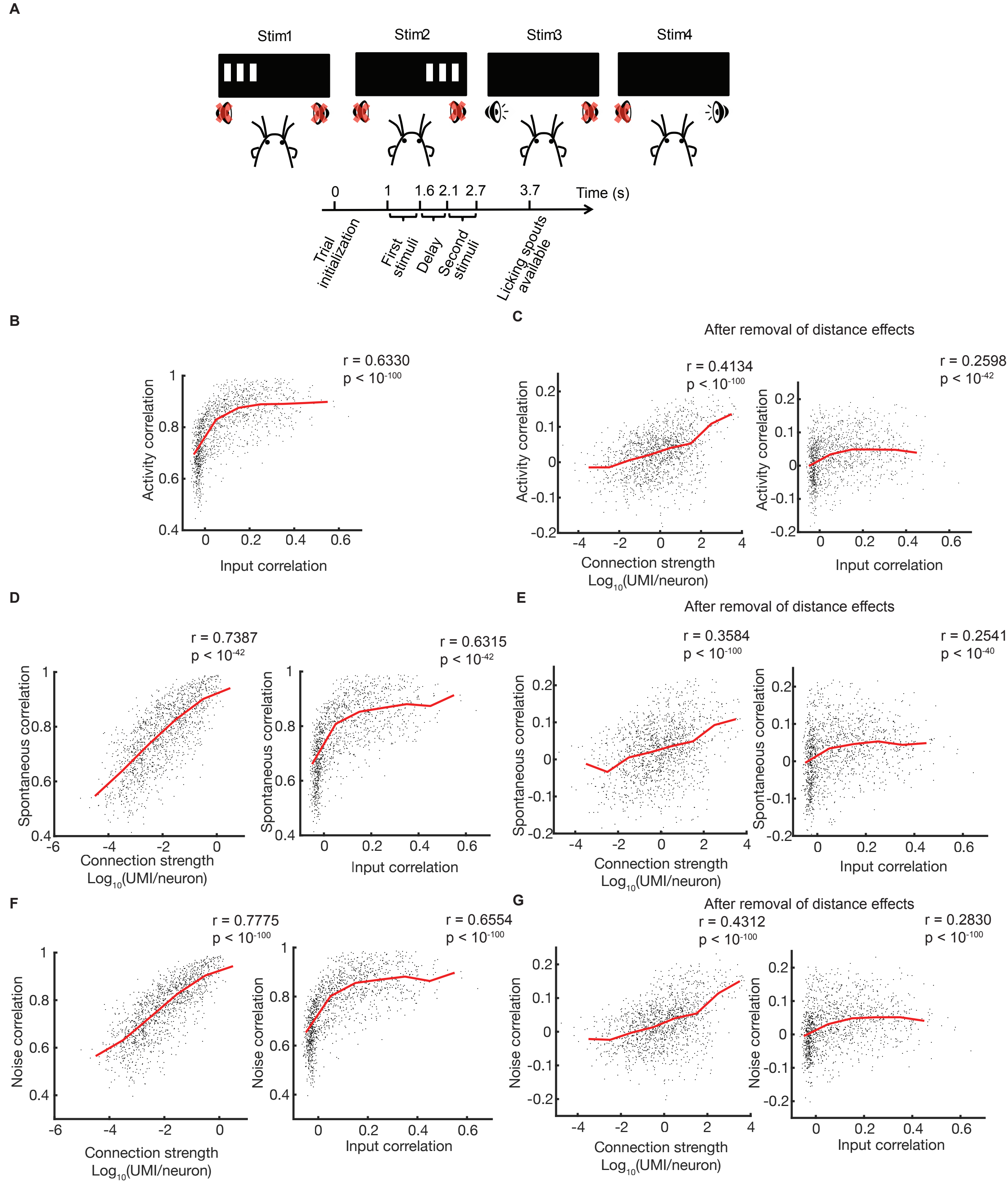

A
Stim1
Stim2
Stim3
Stim4
0
1
1.6
2.1
2.7
3.7
Time (s)
Trial initialization
First stimuli
Delay
Second stimuli
Licking spouts available
B
r = 0.6330
p < 10^-100
Activity correlation
Input correlation
C
After removal of distance effects
r = 0.4134
p < 10^-100
r = 0.2598
p < 10^-42
Activity correlation
Connection strength
Log10(UMI/neuron)
Input correlation
D
r = 0.7387
p < 10^-42
r = 0.6315
p < 10^-42
Spontaneous correlation
Connection strength
Log10(UMI/neuron)
Input correlation
E
After removal of distance effects
r = 0.3584
p < 10^-100
r = 0.2541
p < 10^-40
Spontaneous correlation
Connection strength
Log10(UMI/neuron)
Input correlation
F
r = 0.7775
p < 10^-100
r = 0.6554
p < 10^-100
Noise correlation
Connection strength
Log10(UMI/neuron)
Input correlation
G
After removal of distance effects
r = 0.4312
p < 10^-100
r = 0.2830
p < 10^-100
Noise correlation
Connection strength
Log10(UMI/neuron)
Input correlation

Figure S7

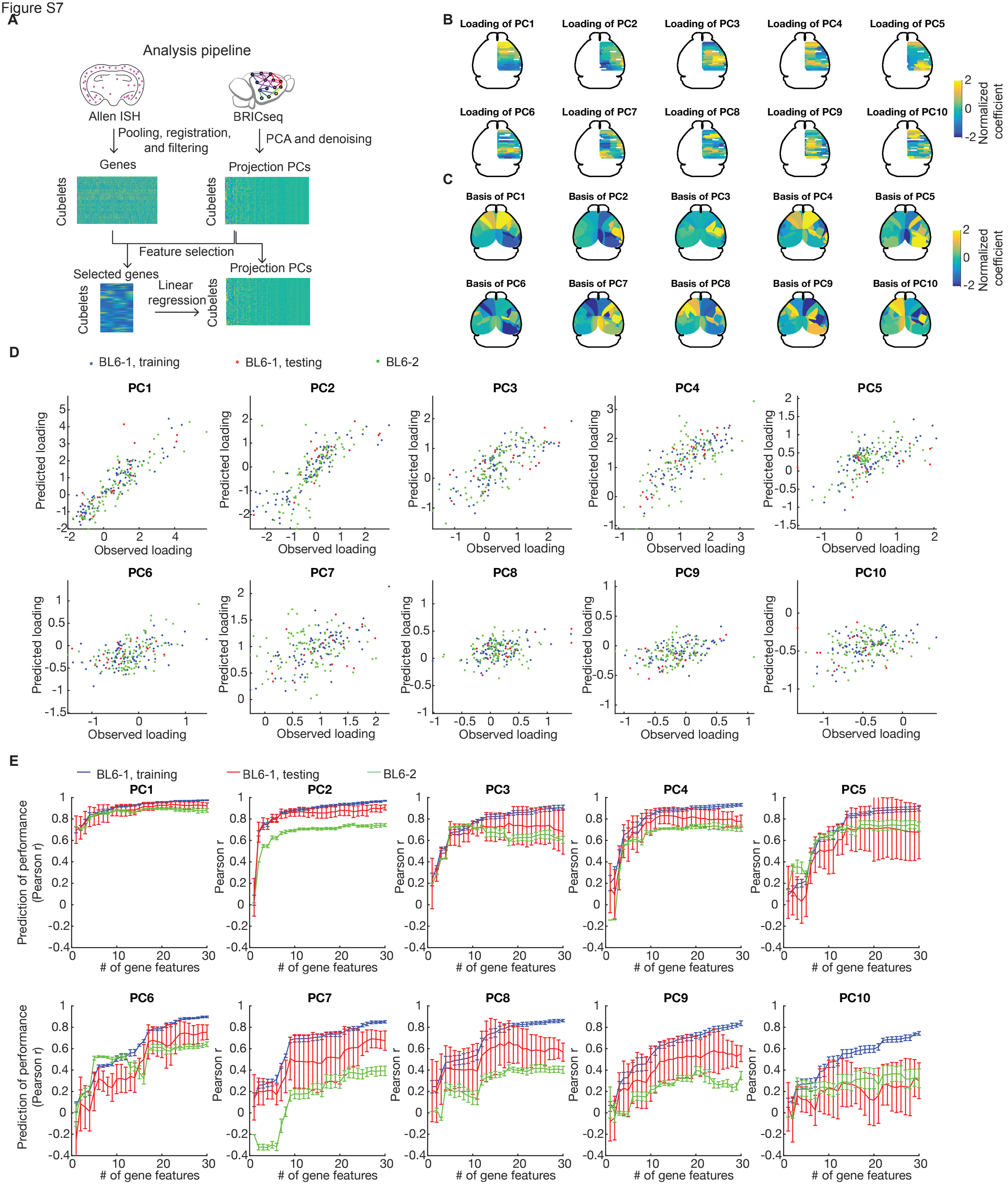

A
Analysis pipeline
Allen ISH
BRICseq
Pooling, registration, and filtering
PCA and denoising
Genes
Projection PCs
Cubelets
Feature selection
Selected genes
Linear regression
B
Loading of PC1
Loading of PC2
Loading of PC3
Loading of PC4
Loading of PC5
Loading of PC6
Loading of PC7
Loading of PC8
Loading of PC9
Loading of PC10
Normalized coefficient
C
Basis of PC1
Basis of PC2
Basis of PC3
Basis of PC4
Basis of PC5
Basis of PC6
Basis of PC7
Basis of PC8
Basis of PC9
Basis of PC10
D
BL6-1, training
BL6-1, testing
BL6-2
PC1
PC2
PC3
PC4
PC5
PC6
PC7
PC8
PC9
PC10
Predicted loading
Observed loading
E
BL6-1, training
BL6-1, testing
BL6-2
Prediction of performance (Pearson r)
Pearson r
### of gene features

Figure S8

**A**

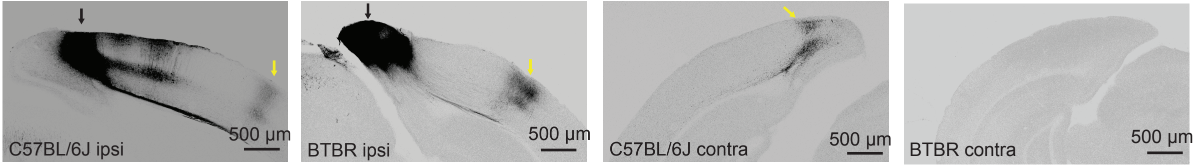

C57BL/6J ipsi
500 µm
BTBR ipsi
500 µm
C57BL/6J contra
500 µm
BTBR contra
500 µm

Figure S9

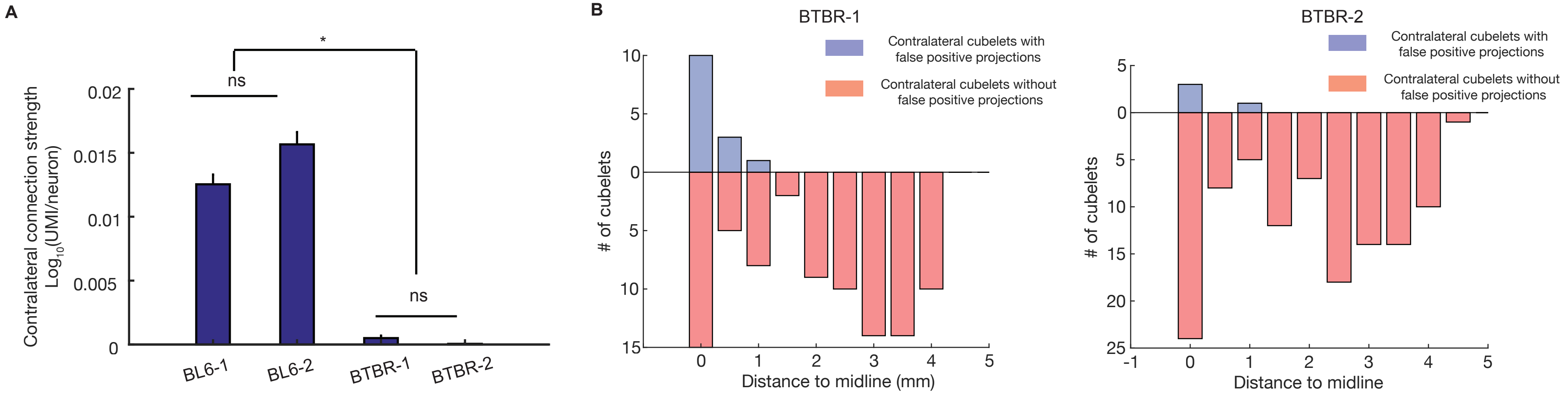
A
B
*
ns
ns
Contralateral connection strength
Log10(UMI/neuron)
0.02
0.015
0.01
0.005
0
BL6-1
BL6-2
BTBR-1
BTBR-2
BTBR-1
Contralateral cubelets with false positive projections
Contralateral cubelets without false positive projections
### of cubelets
Distance to midline (mm)
BTBR-2
Contralateral cubelets with false positive projections
Contralateral cubelets without false positive projections
### of cubelets
Distance to midline

Figure S10

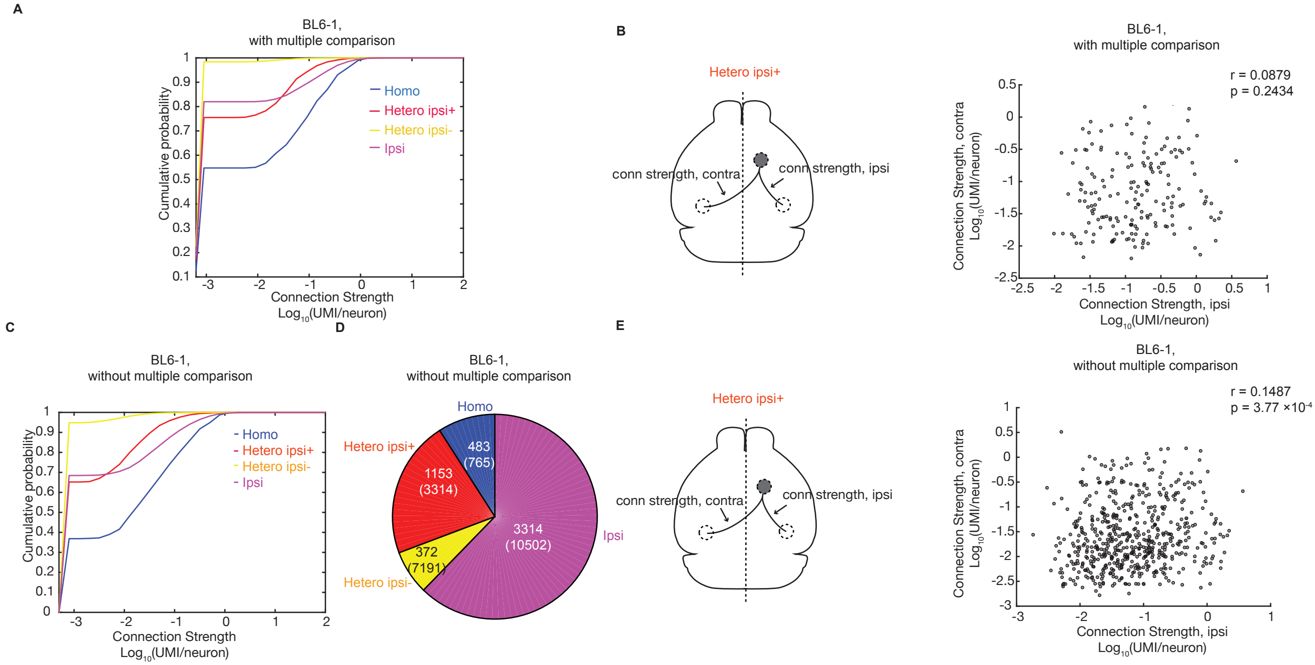

A
BL6-1,
with multiple comparison
Cumulative probability
Connection Strength
Log10(UMI/neuron)
Homo
Hetero ipsi+
Hetero ipsi-
Ipsi
B
Hetero ipsi+
conn strength, contra
conn strength, ipsi
BL6-1,
with multiple comparison
r = 0.0879
p = 0.2434
Connection Strength, contra
Log10(UMI/neuron)
Connection Strength, ipsi
Log10(UMI/neuron)
C
BL6-1,
without multiple comparison
Cumulative probability
Connection Strength
Log10(UMI/neuron)
Homo
Hetero ipsi+
Hetero ipsi-
Ipsi
D
BL6-1,
without multiple comparison
Homo
483
(765)
Hetero ipsi+
1153
(3314)
Hetero ipsi-
372
(7191)
Ipsi
3314
(10502)
E
Hetero ipsi+
conn strength, contra
conn strength, ipsi
BL6-1,
without multiple comparison
r = 0.1487
p = 3.77 ×10-4
Connection Strength, contra
Log10(UMI/neuron)
Connection Strength, ipsi
Log10(UMI/neuron)

Figure S11

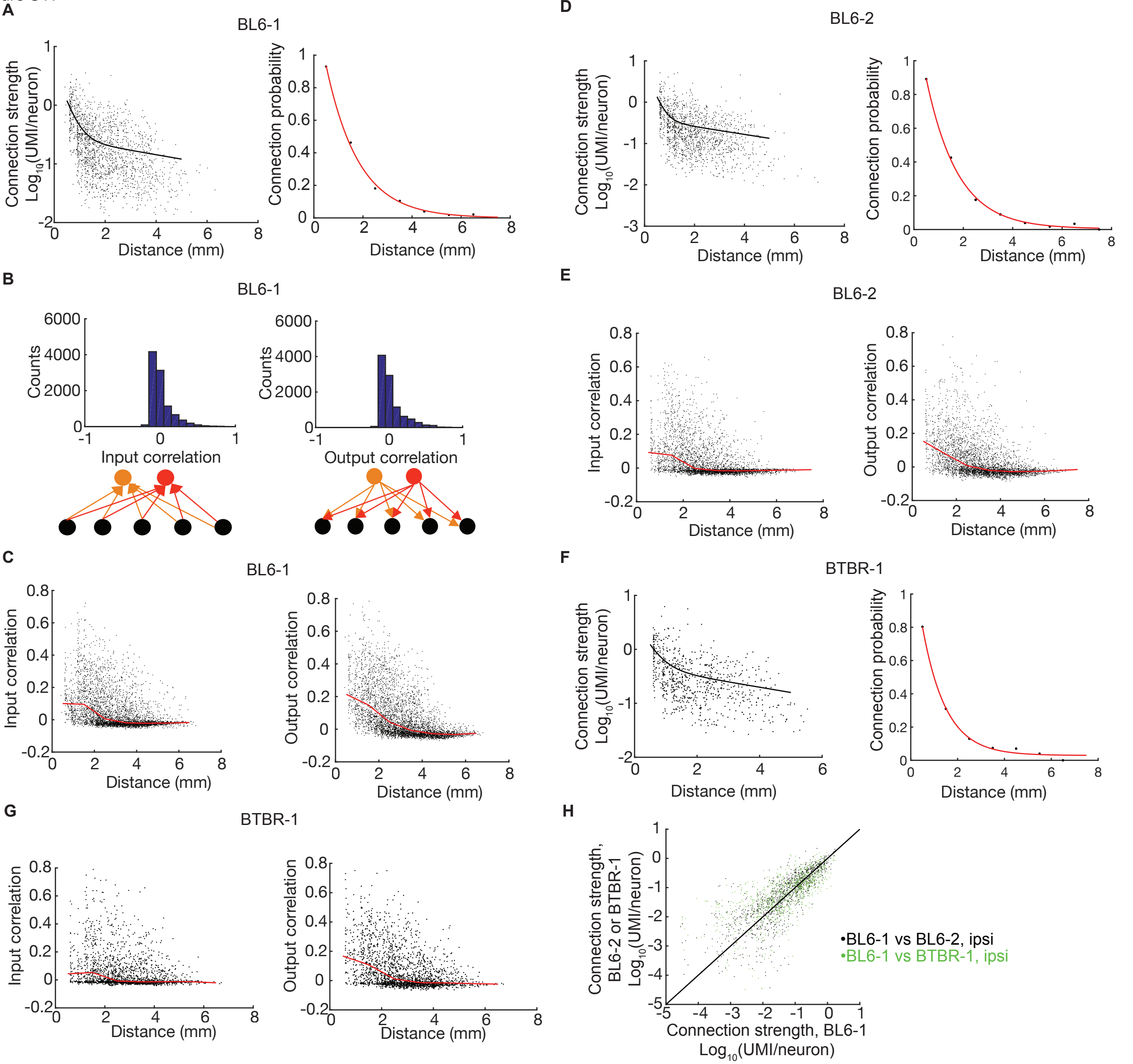

A
BL6-1
Connection strength
$Log_{10}$(UMI/neuron)
Connection probability
Distance (mm)
B
BL6-1
Counts
Input correlation
Output correlation
C
BL6-1
Input correlation
Output correlation
Distance (mm)
D
BL6-2
Connection strength
$Log_{10}$(UMI/neuron)
Connection probability
Distance (mm)
E
BL6-2
Input correlation
Output correlation
Distance (mm)
F
BTBR-1
Connection strength
$Log_{10}$(UMI/neuron)
Connection probability
Distance (mm)
G
BTBR-1
Input correlation
Output correlation
Distance (mm)
H
Connection strength,
BL6-2 or BTBR-1
$Log_{10}$(UMI/neuron)
•BL6-1 vs BL6-2, ipsi
•BL6-1 vs BTBR-1, ipsi
Connection strength, BL6-1
$Log_{10}$(UMI/neuron)

Figure S12

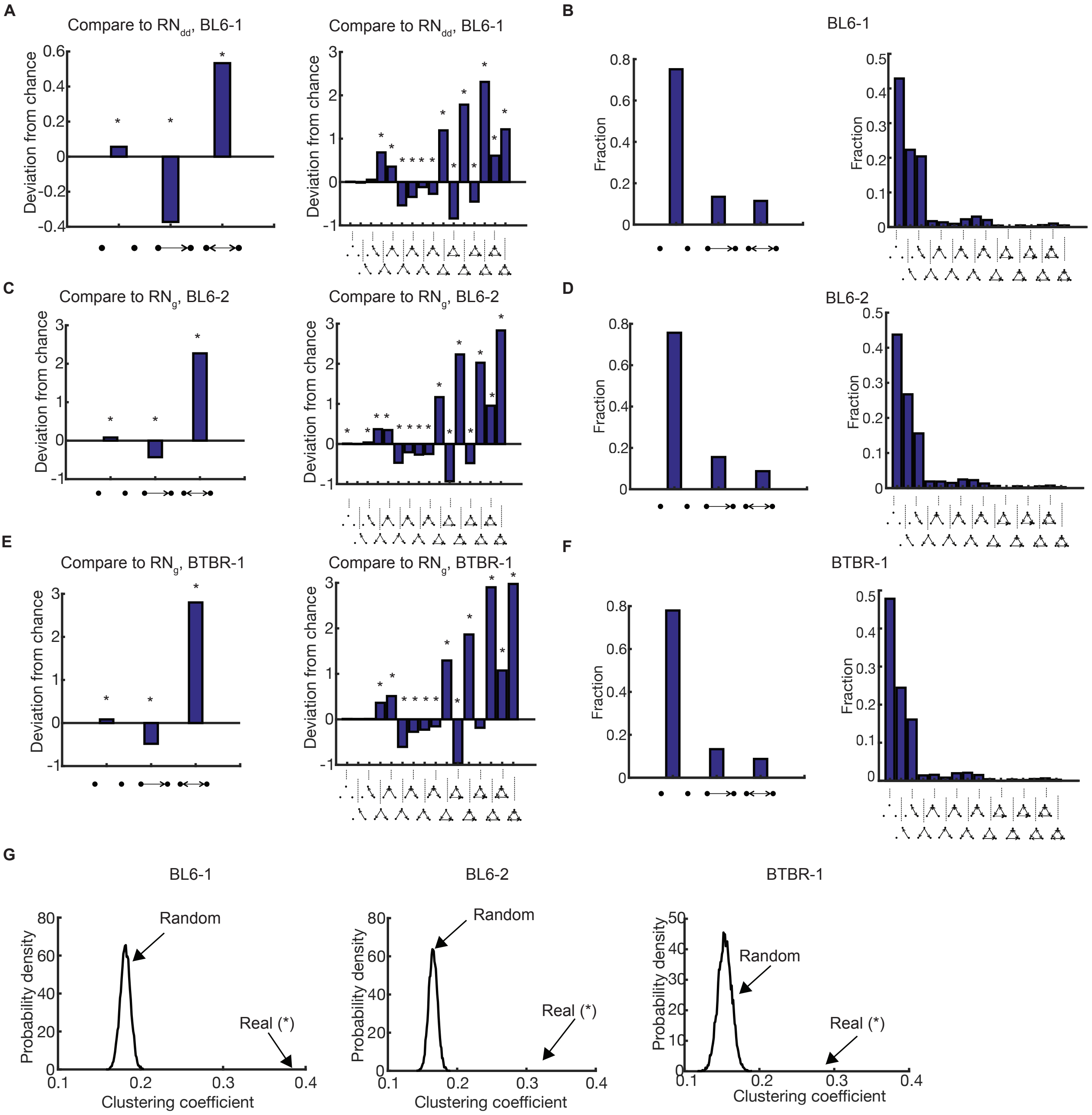

A
Compare to RN$_{dd}$, BL6-1
Deviation from chance
B
BL6-1
Fraction
C
Compare to RN$_g$, BL6-2
D
BL6-2
E
Compare to RN$_g$, BTBR-1
F
BTBR-1
G
BL6-1
BL6-2
BTBR-1
Random
Real (*)
Probability density
Clustering coefficient

Figure S13

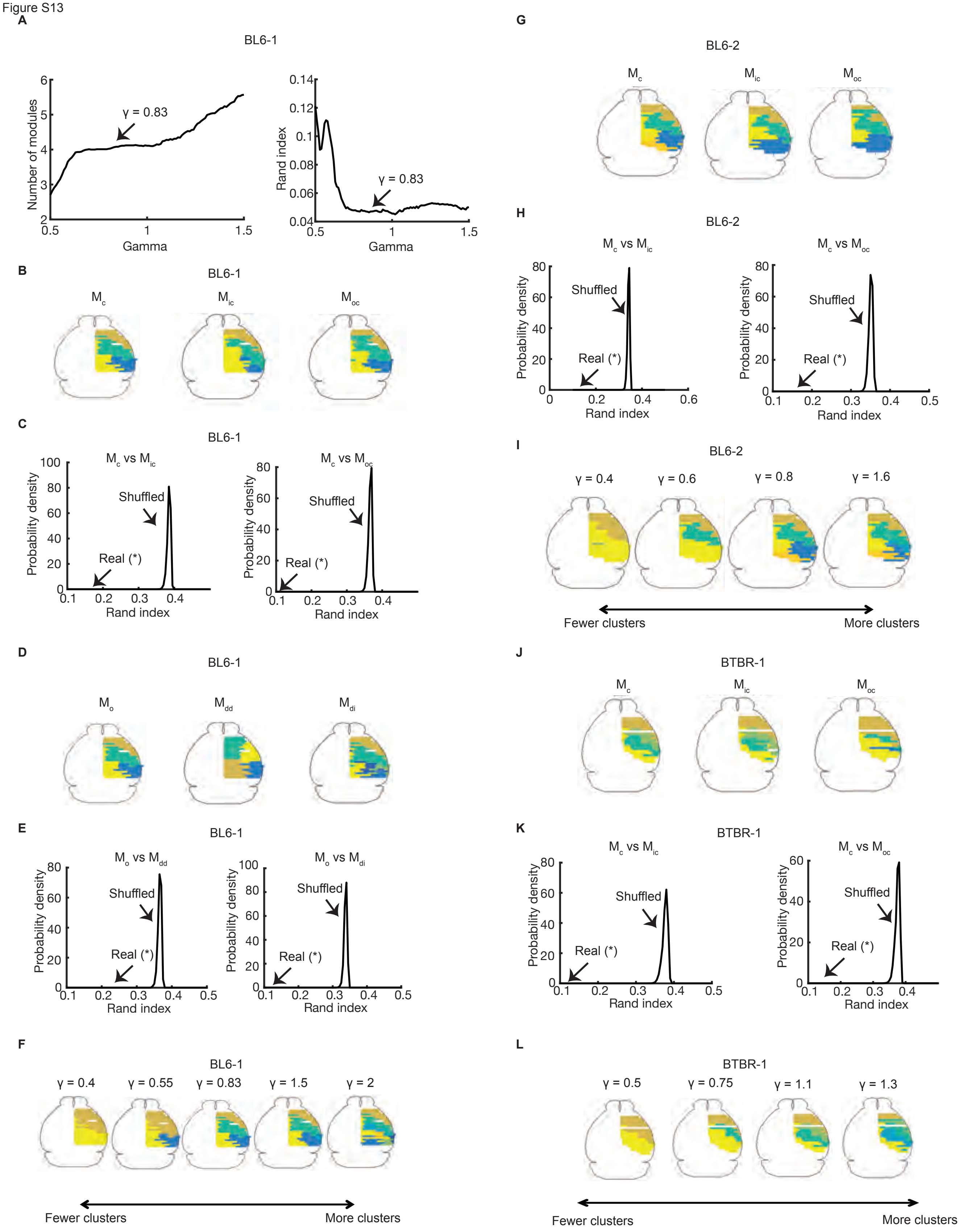
A
BL6-1
Number of modules
Gamma
γ = 0.83
Rand index
Gamma
γ = 0.83
B
BL6-1
$M_{c}$
$M_{ic}$
$M_{oc}$
C
BL6-1
$M_{c}$ vs $M_{ic}$
$M_{c}$ vs $M_{oc}$
Probability density
Shuffled
Real (*)
Rand index
D
BL6-1
$M_{o}$
$M_{dd}$
$M_{di}$
E
BL6-1
$M_{o}$ vs $M_{dd}$
$M_{o}$ vs $M_{di}$
Probability density
Shuffled
Real (*)
Rand index
F
BL6-1
γ = 0.4
γ = 0.55
γ = 0.83
γ = 1.5
γ = 2
Fewer clusters
More clusters
G
BL6-2
$M_{c}$
$M_{ic}$
$M_{oc}$
H
BL6-2
$M_{c}$ vs $M_{ic}$
$M_{c}$ vs $M_{oc}$
Probability density
Shuffled
Real (*)
Rand index
I
BL6-2
γ = 0.4
γ = 0.6
γ = 0.8
γ = 1.6
Fewer clusters
More clusters
J
BTBR-1
$M_{c}$
$M_{ic}$
$M_{oc}$
K
BTBR-1
$M_{c}$ vs $M_{ic}$
$M_{c}$ vs $M_{oc}$
Probability density
Shuffled
Real (*)
Rand index
L
BTBR-1
γ = 0.5
γ = 0.75
γ = 1.1
γ = 1.3
Fewer clusters
More clusters
